# The multi-peak adaptive landscape of crocodylomorph body size evolution

**DOI:** 10.1101/405621

**Authors:** Pedro L. Godoy, Roger B. J. Benson, Mario Bronzati, Richard J. Butler

**Affiliations:** School of Geography, Earth and Environmental Sciences, University of Birmingham, UK.; Department of Earth Sciences, University of Oxford, UK.; Laboratório de Paleontologia de Ribeirão Preto, FFCLRP, Universidade de São Paulo, Ribeirão Preto, Brazil.

**Keywords:** Crocodylomorpha, Crocodyliformes, body size evolution, adaptive landscape, phylogenetic comparative methods, Ornstein–Uhlenbeck models

## Abstract

**Background:** Little is known about the long-term patterns of body size evolution in Crocodylomorpha, the > 200-million-year-old group that includes living crocodylians and their extinct relatives. Extant crocodylians are mostly large-bodied (3–7 m) predators. However, extinct crocodylomorphs exhibit a wider range of phenotypes, and many of the earliest taxa were much smaller (< 1.2 m). This suggests a pattern of size increase through time that could be caused by multi-lineage evolutionary trends of size increase or by selective extinction of small-bodied species. In this study, we characterise patterns of crocodylomorph body size evolution using a model fitting-approach (with cranial measurements serving as proxies). We also estimate body size disparity through time and quantitatively test hypotheses of biotic and abiotic factors as potential drivers of crocodylomorph body size evolution.

**Results:** Crocodylomorphs reached an early peak in body size disparity during the Late Jurassic, and underwent essentially continually decreases in disparity since then. A multi-peak Ornstein-Uhlenbeck model outperforms all other evolutionary models fitted to our data (including both uniform and non-uniform), indicating that the macroevolutionary dynamics of crocodylomorph body size are better described within the concept of an adaptive landscape, with most body size variation emerging after shifts to new macroevolutionary regimes (analogous to adaptive zones). We did not find support for a consistent evolutionary trend towards larger sizes among lineages (i.e., Cope’s rule), or strong correlations of body size with climate. Instead, the intermediate to large body sizes of some crocodylomorphs are better explained by group-specific adaptations. In particular, the evolution of a more aquatic lifestyle (especially marine) correlates with increases in average body size, though not without exceptions.

**Conclusions:** Shifts between macroevolutionary regimes provide a better explanation of crocodylomorph body size evolution than do climatic factors, suggesting a central role for lineage-specific adaptations rather than climatic forcing. Shifts leading to larger body sizes occurred in most aquatic and semi-aquatic groups. This, combined with extinctions of groups occupying smaller body size regimes (particularly during the Late Cretaceous and Cenozoic), gave rise to the upward-shifted body size distribution of extant crocodylomorphs compared to their smaller-bodied terrestrial ancestors.

## Background

Body size influences many aspects of ecology, physiology and evolutionary history [1, 2, 3, 4, 5, 6], and patterns of animal body size evolution are a long-standing subject of macroevolutionary investigation (e.g., [7, 8, 9, 10, 11]). As a major focus of natural selection, it is expected that significant variation should occur in the body size of animals, although confined within biological constraints, such as skeletal structure, thermoregulation and resource availability [4, 5, 12]. Furthermore, body size can often be easily measured or estimated from both fossil and modern specimens, and has therefore been widely used in phenotypic macroevolutionary studies [5, 7, 8, 9, 11, 13, 14, 15, 16, 17].

With few exceptions (e.g., [18, 19]), previous studies of tetrapod body size evolution have focused on mammals (e.g., [14, 15, 16, 20, 21, 22, 23, 24]) and dinosaurs or birds (e.g., [25, 26, 27, 28, 29, 30, 31, 32, 33]). Little is known, however, about other diverse and morphologically disparate clades. Among those, Crocodylomorpha represents an excellent group for studying large-scale evolutionary patterns, with a rich and well-studied fossil record covering more than 200 million years, as well as living representatives [34, 35, 36]. Previous work has investigated multiple aspects of crocodylomorph macroevolution, including spatial and temporal patterns of diversity [35, 36, 37], as well as morphological variation, disparity, and evolution, with a particular focus on the skull [38, 39, 40, 41, 42, 43, 44].

Nevertheless, studies quantitatively investigating macroevolutionary patterns of body size in crocodylomorphs have been restricted to particular time periods (e.g., Triassic-Jurassic body size disparity [45]) or clades (e.g., metriorhynchids [46]), limiting broader interpretations. For instance, the impact of environmental temperature on the growth and adult body size of animals has long been acknowledged as an important phenomenon [4] and has been considered to have a significant influence on the physiology and distribution of crocodylians [47, 48]. There is also strong evidence for climate-driven biodiversity patterns in the group (e.g., [36, 37]). Nevertheless, it remains unclear whether extrinsic factors, such as temperature and geographic distribution, have impacted long-term patterns of crocodylomorph body size evolution [49].

Most of the earliest crocodylomorphs, such as *Litargosuchus* and *Hesperosuchus*, were small-bodied animals (with estimated total lengths of less than 1 metre [50, 51]), contrasting with some giant forms that appeared later, such as *Sarcosuchus* and *Deinosuchus* (possibly more than 10 metres long [52, 53]), as well as with the intermediate to large sizes of extant crocodylians (1.5–7 m [54, 55]). The absence of small-bodied forms among extant species raises questions about what long-term macroevolutionary process (or processes) gave rise to the prevalence of larger body sizes observed in the present. Directional trends of increasing body size through time (see [56]), differential extinction of small bodied taxa, or other factors, such as climate- or environment-driven evolutionary change could explain this. However, because patterns of body size evolution along phylogenetic lineages of crocodylomorphs have not been characterised, its causes are unaddressed.

### Model-fitting approach

Since the end of the last century, palaeontologists have more frequently used quantitative comparative methods to investigate the tempo and mode of evolution along phylogenetic lineages [57, 58, 59], including studies of body size evolution [5, 14, 27, 29, 15, 60]. More recently, numerous studies have employed a phylogeny-based model-fitting approach, using a maximum-likelihood or Bayesian framework to identify the best-fitting statistical macroevolutionary model for a given phylogenetic comparative dataset [31, 33, 61, 62, 63, 64, 65]. Many of those works have tested the fit of a uniform macroevolutionary model, with a single set of parameters applied across all branches of a phylogeny (e.g., [46, 64, 66, 67]). Uniform models are important for describing many aspects of phenotypic evolution and are often the null hypothesis in such studies. However, if the dynamics of evolutionary trends vary in more complex ways through time and space and among clades and environments [e.g., 68, 69, 70, 71, 72], then uniform models might not be adequate to characterise this variation.

Because we aim to characterise variation in body size among many subgroups inhabiting different environments and encompassing substantial variation in morphology, we approach the study of crocodylomorph body size evolution using non-uniform models. We focus on the concept of a Simpsonian Adaptive Landscape [73, 74], which has proved to be a fruitful conceptual framework for characterizing macroevolutionary changes, encompassing ideas such as adaptive zone invasion and quantum evolution [71, 75, 76]. Macroevolutionary landscapes provide a conceptual bridge for dialogues between studies of micro- and macroevolution, and have benefitted from the subsequent advancements of molecular biology and genetics [77]. Within this paradigm, uniform models primarily represent static macroevolutionary landscapes, with unchanged peaks (or maximum adaptive zones [11]) persisting through long time intervals and across the phylogeny [71, 74, 75].

Incorporating biological realism into statistical models of evolution is challenging [78]. Many existing models are based on a Brownian motion (BM) process resulting from random walks of trait values along independent phylogenetic lineages [57, 75, 79]. Uniform Brownian motion has many interpretations. For example, it can be used as a model of drift, or of adaptive evolution towards lineage-specific selective optima that undergo random walks through time, and seems reasonable for describing undirected and unconstrained stochastic change [57]. Elaborations of BM models include the “trend” model, which incorporates a tendency for directional evolution by adding a parameter μ [80. The multi-regime “trend-shift” model has also been proposed, in which the trend parameter (μ) undergoes clade-specific or time-specific shifts (G. Hunt in [33]).

The Ornstein–Uhlenbeck (OU) process [58, 61, 64, 81, 82] is a modification of Brownian motion that incorporates attraction (α) to a trait ‘optimum’ (θ). OU models describe the evolution of a trait towards or around a stationary peak or optimum value, at a given evolutionary rate. Thus, multi-regime OU models can account for the existence of multiple macroevolutionary regimes (similar to adaptive zones, in the Simpsonian Adaptive Landscape paradigm). Even though many OU-based models typically require *a priori* adaptive hypotheses for inferring the trait optima of regimes [61], more recent methods attempt to solve this problem by estimating location, values and magnitudes of regime shifts without *a priori* designation of selective regimes [71, 78, 83]. In particular, the SURFACE method [83] aims to identify shifts in macroevolutionary regimes, identified using AICc (Akaike’s information criterion for finite sample sizes [84]). Originally designated to identify “convergent” trait evolution across phylogenetic lineages, the SURFACE algorithm makes use of a multi-peak OU- model and can be a tool to determine heterogeneity of macroevolutionary landscapes [33, 85, 86]. In this work, we employ a model-fitting approach, using non-uniform macroevolutionary OU-based models (SURFACE), to characterize the adaptive landscape of body size evolution in Crocodylomorpha. This represents the first comprehensive investigation of large-scale patterns of body size evolution across the entire evolutionary history of crocodylomorphs.

## Methods

### Proxy for body size

Extinct Crocodylomorpha are morphologically diverse, and frequently known from incomplete remains. Therefore, precise estimation of their body sizes, and those of comparable fossil groups, can be challenging (see [87, 88] for related considerations). There are many methods and equations for estimating crocodylomorph body size (either body mass or length) available in the literature. The most frequently used equations are derived from linear regressions based on specimens of modern species, using both cranial [53, 89, 90, 91, 92, 93] and postcranial [94, 95] measurements as proxies, even though some inaccuracy is expected (see Additional file 1 for further discussion).

We sought an appropriate proxy for studying body size across all crocodylomorph evolutionary history that also maximised available sample size, to allow as comprehensive a study of evolutionary history as possible. Thus, we decided to use two cranial measurements as proxies for total body length: total dorsal cranial length (DCL) and dorsal orbito-cranial length (ODCL), which is measured from the anterior margin of the orbit to the posterior margin of the skull. By using cranial measurements instead of estimated total body length, we are ultimately analysing patterns of cranial size evolution in crocodylomorphs. Nevertheless, by doing this we also avoid the addition of errors to our model-fitting analyses, since previous works have reported problems when estimating total body length from cranial measurements, particularly skull length (e.g., [46, 88, 96, 97]), as the equations were formulated using modern species and different crocodylomorph clades are likely to have body proportions distinct from those of living taxa (see Additional file 1). Furthermore, the range of body sizes among living and extinct crocodylomorphs is considerably greater than variation among size estimates for single species. Therefore, we expect to recover the most important macroevolutionary body size changes in our analyses even when using only cranial measurements. The use of ODCL, in addition to DCL, is justified as it allows us to examine the sensitivity of our results to changes in proportional snout length, as a major aspect of length change in crocodylomorph skulls results from proportional elongation or shortening of the snout [98, 99, 100]. Also, more taxa could be included in our analyses when doing so, because ODCL can be measured from some incomplete skulls.

The DCL dataset includes 219 specimens (representing 178 taxa), whereas the ODCL dataset includes 240 specimens (195 taxa). In total, measurements from 118 specimens (83 taxa) were collected via first-hand examination from specimens, using callipers and measuring tape. The remaining information was collected from the literature (98 specimens) or photographs (21 specimens) supplied by other researchers, and measurements were estimated using the software ImageJ (see Additional file 2 for the complete list of sampled specimens). We used mean values in those cases where we had cranial measurements for multiple specimens of the same taxon. For both the model-fitting and correlation analyses, we used log-transformed skull measurements in millimetres. However, to help us further interpret and discuss our results, total body length was subsequently estimated using the equations presented by [91].

### Phylogenetic framework

For the phylogenetic framework of Crocodylomorpha, the aim was to maximise taxon inclusion and to use a phylogenetic hypothesis that best represents the current consensus. We primarily used an informally modified version of the supertree presented by Bronzati et al. [35], which originally contained 245 taxa. We added recently published species, and removed taxa that have not yet received a formal description and designation. Also, species not previously included in phylogenetic studies but for which we had body size data were included based on the phylogenetic positions of closely related taxa (see Additional file 1 for more information on the supertree construction). Thus, our updated version of the supertree contains 296 crocodylomorph species, as well as nine closely related taxa used as outgroups for time-scaling the trees (see below).

To accommodate major uncertainties in crocodylomorph phylogeny, we also constructed two other supertrees, with alternative topologies, varying the position of Thalattosuchia. Thalattosuchians are Jurassic–Early Cretaceous aquatic crocodylomorphs, some of which were probably fully marine [101]. They have classically been placed within Neosuchia, as the sister taxon of Tethysuchia [99]. Nevertheless, some authors have argued that this close relationship may result from the convergent acquisition of longirostrine snouts in both groups [98, 102], and some recent works have suggested alternative positions for Thalattosuchia, within or as the sister group of Crocodyliformes (i.e., only distantly related to Neosuchia [100, 103, 104, 105]). Accordingly, to test the influence of uncertainty over the phylogenetic position of Thalattosuchia, we performed our macroevolutionary analyses using three distinct phylogenetic scenarios of Crocodylomorpha (Fig. 1). In the first, the more classic position of Thalattosuchia was maintained (Thalattosuchia as the sister taxon of Tethysuchia and within Neosuchia; as in the original supertrees of Bronzati et al. [34, 35]). In the two alternative phylogenetic scenarios, Thalattosuchia was placed as the sister group of either Crocodyliformes (as non-crocodyliform crocodylomorphs), following the position proposed by Wilberg [100], or Mesoeucrocodylia (as the sister group of the clade formed by Neosuchia + Notosuchia in our topologies), following Larsson & Sues [106] and Montefeltro et al. [104]. Discrepancies among competing phylogenetic hypotheses do not concern only the “thalattosuchian problem” described above. However, our decision to further investigate only the impact of the different positions of Thalattosuchia is based on its high taxic diversity and the impact that its phylogenetic position has on branch lengths across multiple parts of the tree, factors that can substantially alter macroevolutionary patterns detected by our analyses.

**Fig. 1.**
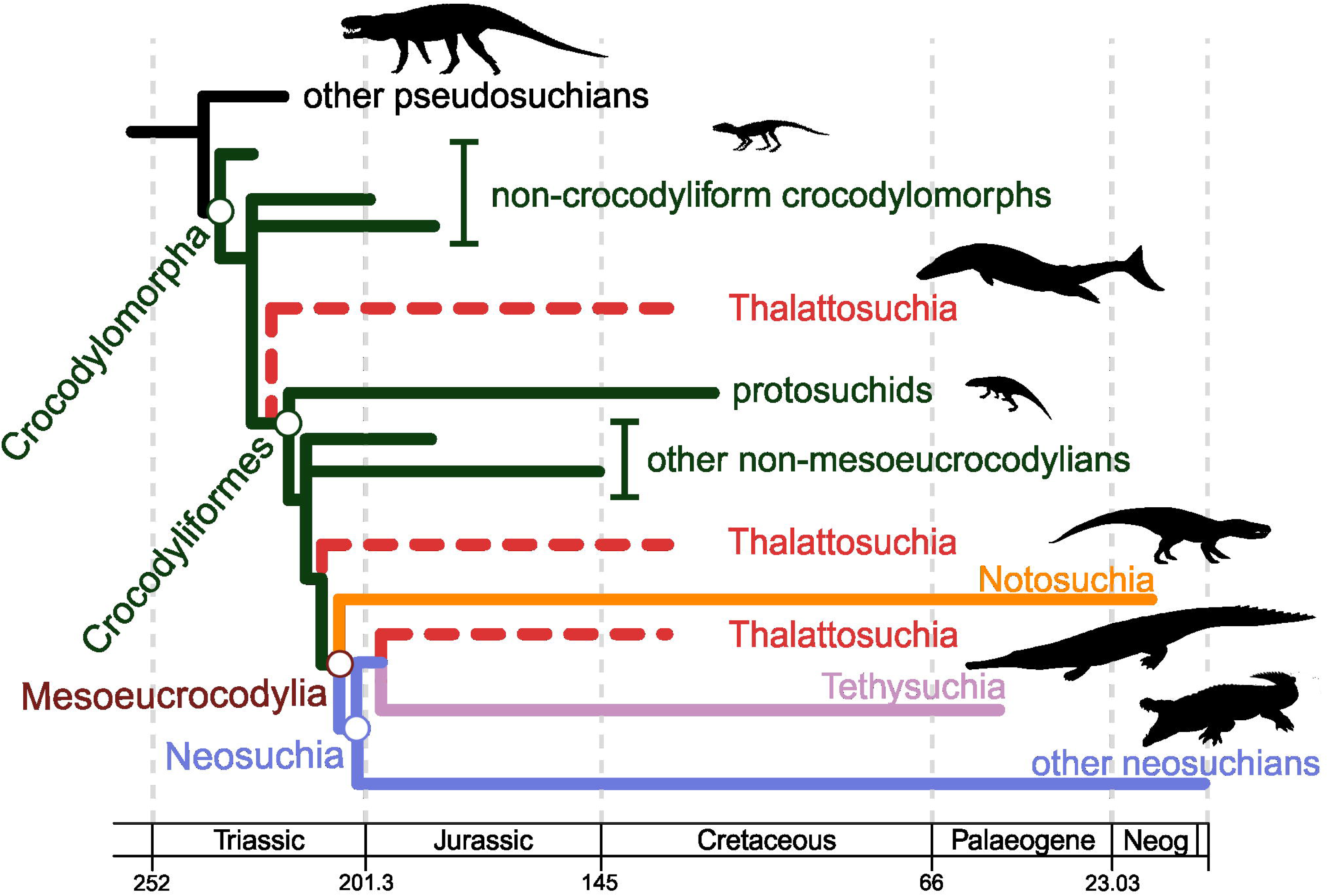
Simplified cladogram showing the phylogenetic relationships among crocodylomorphs and the alternative positions of Thalattosuchia (dashed red lines), following hypotheses proposed by [34, 35, 109, 113, 115]. Silhouettes are from phylopic.org.

### Time-scaling method

Calibration of the phylogeny to time [107] is a crucial step in comparative analyses of trait evolution, and the use of different methods may impact upon the inference of evolutionary models and the interpretation of results [108, 109]. As such, we decided to use a tip-dating approach using the fossilised birth-death (FBD) model [110]. The FBD method is a Bayesian total-evidence dating approach which uses a birth-death process that includes the probability of fossilization and sampling to model the occurrence of fossil species in the phylogeny and estimate divergence times (=node ages) [111, 112, 113, 114]. Information on occurrence times of all species in the supertree (=tip ages) were initially obtained from the Paleobiology Database, but were then checked using primary sources in the literature. Fossil ages were represented by uncertainty bounds of their occurrences. We then generated an “empty” morphological matrix for performing Bayesian Markov chain Monte Carlo (MCMC) analyses in MrBayes version 3.2.6 [115], following the protocol within the R package *paleotree* version 3.1.3 [116]. We used our supertree topologies (with alternative positions of Thalattosuchia) as topological constraints and set uniform priors on the age of tips based on the occurrence dates information. We used a uniform prior for the root of the tree (for all three topologies/phylogenetic scenarios), constrained between 245 and 260 Myr ago. This constraint was used because the fossil record indicates that a crocodylomorph origin older than the Early Triassic is unlikely [117, 118]. For each topology, 10,000,000 generations were used, after which the parameters indicated that both MCMC runs seemed to converge (i.e., the Potential Scale Reduction Factor approached 1.0 and average standard deviation of split frequencies was below 0.01).

For each topology, we randomly sampled 20 trees (henceforth: FBD trees) from the posterior distribution after a burn-in of 25%. This resulted in 60 time-scaled, completely resolved crocodylomorph trees that were used in our macroevolutionary model comparisons. Similar numbers of trees were used in previous work on dinosaurs [33], mammals [24] and early sauropsids [87]. Analyses across these 60 trees allowed us to characterise the influence of topological and time-scale uncertainty on our results.

Previous studies have demonstrated that time-calibration approaches can impact phylogenetic comparative methods (e.g., [119]). Therefore, we also used other three different time-scaling methods (*minimum branch length*, *cal3* and *Hedman* methods). Differently from the FBD tip-dating method, these three methods belong to the category of *a posteriori* time-scaling (APT) approaches (*sensu* Lloyd et al. [120]), and were used as a sensitivity analysis (see Additional file 1 for further details on the employment of these methods). These additional time-scaling approaches were used only for our initial model comparisons (see below). APT methods were performed in R version 3.5.1 [121], using package *paleotree* [116] (*mbl* and *cal3* methods) and the protocol published by Lloyd et al. [120] (*Hedman* method). Results from macroevolutionary analyses using these APT methods were similar to those using the FBD trees (see the “Results” section) and are therefore not discussed further in the main text (but are included in Additional file 1).

### Macroevolutionary models

We applied a model-fitting approach to characterize patterns of body size evolution in Crocodylomorpha, with an emphasis on evolutionary models based on the Ornstein-Uhlenbeck (OU) process [33, 58, 61, 64, 83]. The first formulation of an OU-based model was proposed by Hansen (1997), based on Felsenstein’s [81] suggestion of using the Ornstein-Uhlenbeck (OU) process as a basis for comparative studies [61, 82]. OU- based models (also known as “Hansen” models) express the dynamics of a quantitative trait evolving along the branches of a phylogeny as the result of stochastic variation around a trait “optimum” (expressed as theta: θ), towards which trait values are deterministically attracted (the strength of attraction is given by alpha: α). The constant σ^2^, describes the stochastic spread of the trait values over time (i.e., under a Brownian motion process). Accordingly, the OU model can be formulated as:

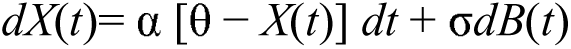

This equation expresses the amount of change in trait *X* during the infinitesimal time interval from *t* to *t* + *dt*. As expressed above, the formulation includes a term describing trait attraction towards θ, which is the product of α and the difference between *X*(*t*) and θ. The term σ*dB*(*t*) describes stochastic evolution in the form of Brownian motion (BM), with random variables of mean zero and variance of *d*t (thus, σ^2^ is the rate of stochastic evolution). In this sense, if α is zero, the attraction term becomes zero, and the result is evolution by BM as a special case of OU [33, 61, 64]. The OU model can also simulate trait evolution patterns similar to that observed under other evolutionary models, such as BM with a trend incorporated, and “white noise” or stasis [33, 58, 64]. Thus, examination of the fitted parameters of the OU model is crucial for interpreting the mode of evolution [58, 61]. For example, the estimated ancestral trait value (i.e., the value of θ at the root of the tree) is given by the parameter Z_0_. Also, by obtaining ln (2)/α, we are calculating the time taken for a new macroevolutionary regime to become more influential than the ancestral regime (i.e., how long it takes to θ to be more influential than Z_0_). This parameter is often called the phylogenetic half-life (or t_0.5_) [58].

Among the methods that attempt to model adaptive evolution under the framework of an Ornstein-Uhlenbeck process (e.g., [78, 82, 122]), the SURFACE algorithm [83] estimates the fit of a non-uniform OU-based model by allowing shifts in trait optima (θ) among macroevolutionary regimes. SURFACE locates regime shifts using stepwise AICc (Akaike’s information criterion for finite sample sizes [84, 123, 124]), with a forward phase (that searches for all regime shifts in the phylogeny) and a backward phase (in which improvements of AICc scores merge similar regimes, detecting “convergent” evolution). Although it allows θ to vary among regimes, SURFACE assumes fixed whole-tree values of σ^2^ and α [83].

We compared the performance of two different OU-based models, one with a single trait optimum or a single macroevolutionary regime (“OU model”) and another non-uniform model with multiple regimes (“SURFACE model”). To test if other macroevolutionary models could provide a better description of the observed patterns of crocodylomorph body size evolution, we also compared the OU-based models with other models. First, a uniform Brownian motion (BM model), which describes diffusive, unconstrained evolution via random walks along independent phylogenetic lineages, resulting in no directional trend in trait mean, but with increasing trait variance (=disparity) through time [57, 62, 63, 64]. Second, the “early burst” (EB model; also known as “ACDC model” [125]), in which the lineages experience an initial maximum in evolutionary rate of change, that decreases exponentially through time according to the parameter *r* [126]. This results in a rapid early increase in trait variance followed by deceleration [125, 126].

We also fitted a uniform (single-regime) and non-uniform (multi-regime) trend-like models. In the uniform “trend” model the parameter μ is incorporated into the BM model to describe directional multi-lineage increase or decrease in trait values through time in the entire clade [62, 63, 80]. Non-uniform “trend” models allow for shifts in the parameter μ, which can be explored in two different ways according to the non-uniform trend models formulated by G. Hunt and presented in Benson et al. [33]: temporal shifts in μ across all contemporaneous lineage (“time-shift trend models”), or shifts at specific nodes of the tree, modifying μ in the descendent clade (“node-shift trend models”). In time-shift trend models, shifts to a new value of μ occurs at time-horizons and are applied to all lineages alive at that time. In node-shift trend models, values of μ can vary among contemporaneous lineages. In a similar approach to the forward phase of SURFACE, the shifts in these non-uniform trend-like models are detected via stepwise AICc. In both time-shift and node-shift models, the Brownian variance (σ^2^) is constant across all regimes [33]. For our macroevolutionary analyses with the entire crocodylomorph phylogeny, we fitted trend-like models that allowed up to three time-shifts and 10 node-shifts to occur, given that analyses with more shifts are computationally intensive and often receive significantly weaker support (following results presented by Benson et al. [33]).

### Initial model comparison

Our initial model comparison involved a set of exploratory analyses to test which evolutionary models (SURFACE, OU, BM, EB and trend-like models) offered the best explanation of our data, using log-transformed cranial measurements (for both DCL and ODCL). To reduce computational demands, we used only one position of Thalattosuchia (i.e., with the group positioned within Neosuchia). The aim here was to compare the performance of the OU-based models, particularly the SURFACE model, against the other BM-based evolutionary models, but also to evaluate possible influences of the different time-scaling methods (we used four different approaches as a sensitivity analysis) and body size proxies. Maximum-likelihood was employed to fit these models to our body size data and the phylogeny of Crocodylomorpha, and we compared the AICc scores of each model.

### Appraisal of spurious model support

Previous works suggested caution when fitting OU models in comparative analyses, since intrinsic difficulties during maximum-likelihood fits can lead to false positives and spurious support to overly complex models [e.g., 127, 128]. This issue may be reduced when using non-ultrametric trees (as done here), as it improves identifiability of the parameters of OU models [64, 127]. We also addressed this by using the phylogenetic Bayesian information criterion (pBIC: proposed by Khabbazian et al. [72]) during the backward phase of model simplification in all our analyses (using the implementation for SURFACE from Benson et al. [33]). The pBIC criterion is more conservative than AICc, in principle favouring simpler models with fewer regimes with lower rates of false positive identification of regime shifts. Although these models were fit using pBIC, they were compared to other models (such as BM, EM and trend-like models) using AICc because pBIC is not implemented for most other models of trait evolution.

Furthermore, to evaluate the influence of spurious support for complex OU models, we simulated data under BM once on each of our 20 phylogenies, using the parameter estimates obtained from the BM model fits to those phylogenies. We then fitted both BM and SURFACE models to the data simulated under BM, and compared several aspects of the results to those obtained from analysis of our empirical body size data (using the ODCL dataset). Specifically, we compared delta-AICc (i.e., the difference between AICc scores received by BM and SURFACE models for each tree), the number of regime shifts obtained by SURFACE, and the values of α obtained by SURFACE. This allowed us to assess whether the results of SURFACE analyses of our empirical data could be explained by overfitting of a highly-parameterised non-uniform model to data that could equally be explained by an essentially uniform process.

### Further SURFACE analyses

Our initial model comparisons provided strong support for the SURFACE model (see the “Results” section). Subsequent analyses therefore focussed on SURFACE, which is particularly useful because it identifies macroevolutionary regimes that provide a simplified map of the major patterns of body size evolution in crocodylomorphs. This second phase of analyses made use of all three alternative phylogenetic scenarios (varying the position of Thalattosuchia) to test the influence of phylogeny in interpretations of evolutionary regimes for body size in Crocodylomorpha. We fitted SURFACE to 20 FBD trees, of each alternative topology, using body size data from the ODCL dataset (our initial model comparisons indicated that both our size indices yielded essentially identical results, and ODCL is available for more taxa). As before, we performed our SURFACE analyses using pBIC [72] during the backward-phase of the algorithm.

### Clade-specific analyses with Notosuchia and Crocodylia

Two well-recognized crocodylomorph subclades, Notosuchia and Crocodylia, returned a relatively high frequency of macroevolutionary regime shifts, representing an apparently complex evolutionary history under the SURFACE paradigm. However, the SURFACE algorithm fits a single value of α to all regimes, and therefore could overestimate the strength of evolutionary constraint within regimes, and consequently miscalculate the number of distinct regimes within clades showing more relaxed patterns of trait evolution. We investigated this possibility by fitting the initial set of evolutionary models (SURFACE, OU, BM, EB and trend-like models) to the phylogenies of these two subclades (using 50 FBD trees for each clade, sampled from the posterior distribution of trees time-scaled with the FBD method) and their body size data (using only the ODCL dataset, since it includes more species). Differently from what was done for the entire crocodylomorph phylogeny, for Notosuchia we fitted trend-like models with up to 2 time-shifts and 5 node-shifts, whereas for Crocodylia we allowed up to 3 time-shifts and 7 node-shifts to occur, given that these two clades include fewer species (70 crocodylians and 34 notosuchians were sampled in our ODCL dataset) and fewer shifts are expected.

In addition, for these same clades, we also employed the OUwie model-fitting algorithm [82], fitting different BM and OU-based models which allow all key parameters to vary freely (since SURFACE allows only θ to vary, whereas it assumes fixed values of σ^2^ and α for the entire tree). However, differently from SURFACE, OUwie needs *a priori* information on the location of regime shifts in order to be implemented. Thus, we incorporated the regime shifts identified by SURFACE into our phylogenetic and body size data (by extracting, for each tree, the regime shifts from previous SURFACE results) to fit four additional evolutionary models using the OUwie algorithm: BMS, which is a multi-regime BM model that allows the rate parameter σ^2^ to vary; OUMV, a multi-regime OU-based model that allows σ^2^and the trait optimum θ to vary; OUMA, also a multi-regime OU model, in which θ and the constraint parameter α can vary; and OUMVA, in which all three parameters (θ, α and σ^2^) can vary. Since computing all these parameter estimates can be an intensively demanding task [82], some of the model fits returned nonsensical values and were, therefore, discarded. Nonsensical values were identified by searching for extremely disparate parameter estimates, among all 50 model fits (e.g., some model fits found σ^2^ values higher than 100,000,000 and α lower than 0.00000001).

All macroevolutionary analyses were performed in R version 3.5.1 [121]. Macroevolutionary models BM, trend, EB, and OU with a single regime were fitted using the R package *geiger* [122]. The SURFACE model fits were performed with package *surface* [83]. Implementation of pBIC functions in the backward-phase of SURFACE model fits, as well as the functions for fitting non-uniform trend-like models, were possible with scripts presented by Benson et al. [33]. Simulated data under BM (for assessing the possibility of spurious support to the SURFACE model) was obtained with package *mvMORPH* [129]. The additional clade-specific model-fitting analyses, using the OUwie algorithm, were implemented with the package *OUwie* [130].

### Correlation with abiotic and biotic factors

To test whether abiotic environmental factors could be driving the evolution and distribution of body sizes in crocodylomorphs, we extracted environmental information from the literature. As a proxy for palaeotemperature, we used δ^18^O data from two different sources. The dataset from Zachos et al. [131] assembles benthic foraminifera isotopic values from the Late Cretaceous (Maastrichtian) to the Recent. The work of Prokoph et al. [132] compiled sea surface isotopic values from a range of marine organisms. Their dataset is divided into subsets representing palaeolatitudinal bands. For our analyses, we used the temperate palaeolatitudinal subset, which extends from the Jurassic to the Recent, but also the tropical palaeolatitudinal subset, which extends back to the Cambrian. For the correlation analyses, we used 10 Myr time bins (see Additional file 1 for information on time bins), by taking the time-weighted mean δ^18^O for data points that fall within each time bin. For the body size data used in the correlation tests, we calculated maximum and mean size values for each time bin, using both DCL and ODCL datasets. Correlations between our body size data and the proxies for palaeotemperature were first assessed using ordinary least squares (OLS) regressions. Then, to avoid potential inflation of correlation coefficients created by temporal autocorrelation (the correlation of a variable with itself through successive data points), we used generalised least squares (GLS) regressions with a first-order autoregressive model incorporated (see e.g., [36, 133, 134, 135]). Furthermore, to test the possible differential influence of temperature on marine versus continental (terrestrial and freshwater) animals, we also created two additional subsets of our data, one with only marine and another with only non-marine crocodylomorphs (ecological information for each taxon was obtained from the PBDB and the literature, e.g., [36, 136]).

We also collected palaeolatitudinal data for every specimen in our dataset from the Paleobiology Database (PBDB) and the literature, and tested the correlation between these and our body size data (DCL and ODCL datasets). To test whether our body size data is correlated with palaeolatitudinal data, we first applied OLS regressions to untransformed data. Then, to deal with possible biases generated by phylogenetic dependency, we used phylogenetic generalized least squares regressions (PGLS [137]), incorporating the phylogenetic information from the maximum clade credibility (MMC) tree, with Thalattosuchia placed within Neosuchia, obtained from our MCMC tip-dating results. For this, branch length transformations were optimised between bounds using maximum-likelihood using Pagel’s λ [138] (i.e., argument λ= “ML” within in the function pgls() of the R package *caper* [139]). As for the correlation analyses between our body size data and palaeotemperature, we also analysed marine and only non-marine taxa separately. To explore the effects of these two abiotic factors on the distribution of body sizes at more restricted levels (temporal and phylogenetic), we repeated our regression analyses using subsets of both ODCL and DCL datasets, including body size data only for species of Crocodylia, Notosuchia, Thalattosuchia, and Tethysuchia. For crocodylians, correlations with paleotemperature were restricted to the Maastrichtian until the Recent (i.e., data from [131]).

We also explored the possible impact of clade-specific evolutionary transitions between the environments on crocodylomorph body size evolution. For that, we obtained ecological information for each taxon from the PBDB and the literature (e.g., [36, 136]), subdividing our body size data (from the ODCL dataset, since it included more taxa) into three discrete categories to represent different generalised ecological lifestyles: terrestrial, semi-aquatic/freshwater, and aquatic/marine. We then used analysis of variance (ANOVA) for pairwise comparisons between different lifestyles. We also accounted for phylogenetic dependency by applying a phylogenetic ANOVA [140], incorporating information from the MCC tree with Thalattosuchia placed within Neosuchia. For both ANOVA and phylogenetic ANOVA, Bonferroni-corrected p-values (q-values) for *post-hoc* pairwise comparisons were calculated. Phylogenetic ANOVA was performed with 100,000 simulations.

All correlation analyses (with abiotic and biotic factors) used log-transformed cranial measurements (DCL or ODCL) in millimetres and were performed in R version 3.5.1 [121]. GLS regressions with an autoregressive model were carried out using the package *nlme* [141], PGLS regressions used the package *caper* [139], whereas phylogenetic ANOVA was performed using the package *phytools* [142].

### Disparity estimation

Important aspects of crocodylomorph body size evolution can be revealed by calculating body size disparity through time. There are different methods and metrics for quantifying morphological disparity (e.g., [143, 144, 145, 146]), and in the present study disparity is represented by the standard deviation of log-transformed body size values included in each time bin. We also plotted minimum and maximum sizes for comparison. Our time series of disparity used the same time bins as for the correlation analyses, with the difference that only time bins with more than three taxa were used for calculating disparity (time bins containing three or fewer taxa were lumped to adjacent time bins; see Additional file 1 for information on time bins). Disparity through time was estimated based on the ODCL dataset (since it includes more taxa).

## Results

### Initial model comparison

Comparisons between the AICc scores for all the evolutionary models fitted to our crocodylomorph body size data (Fig. 2a and b; see Additional file 1: Figures S5 for results of the sensitivity analysis using different time-scaling methods) show extremely strong support (i.e. lower AICc values) for the SURFACE model. This is observed for both body size proxies (DCL and ODCL) and independently of the time-scaling method used. All uniform models exhibit relatively similar AICc scores, including the OU model with a single macroevolutionary regime, and all of these are poorly supported compared to the SURFACE model. For trees calibrated with the FBD methods, all trend-like models (i.e., either uniform or multi-trend models) received very similar support, using both size proxies, and have AICc values that are more comparable to the set of uniform models than to the SURFACE model. Even the best trend-like model (usually the models with two or three node-shifts, which are shown as the “best trend” model in Fig. 2a and b) have significantly weaker support than SURFACE, regardless of the time-calibration method used (see Additional file 3 for a complete list of AICc scores, including for all trend-like models).

**Fig. 2.**
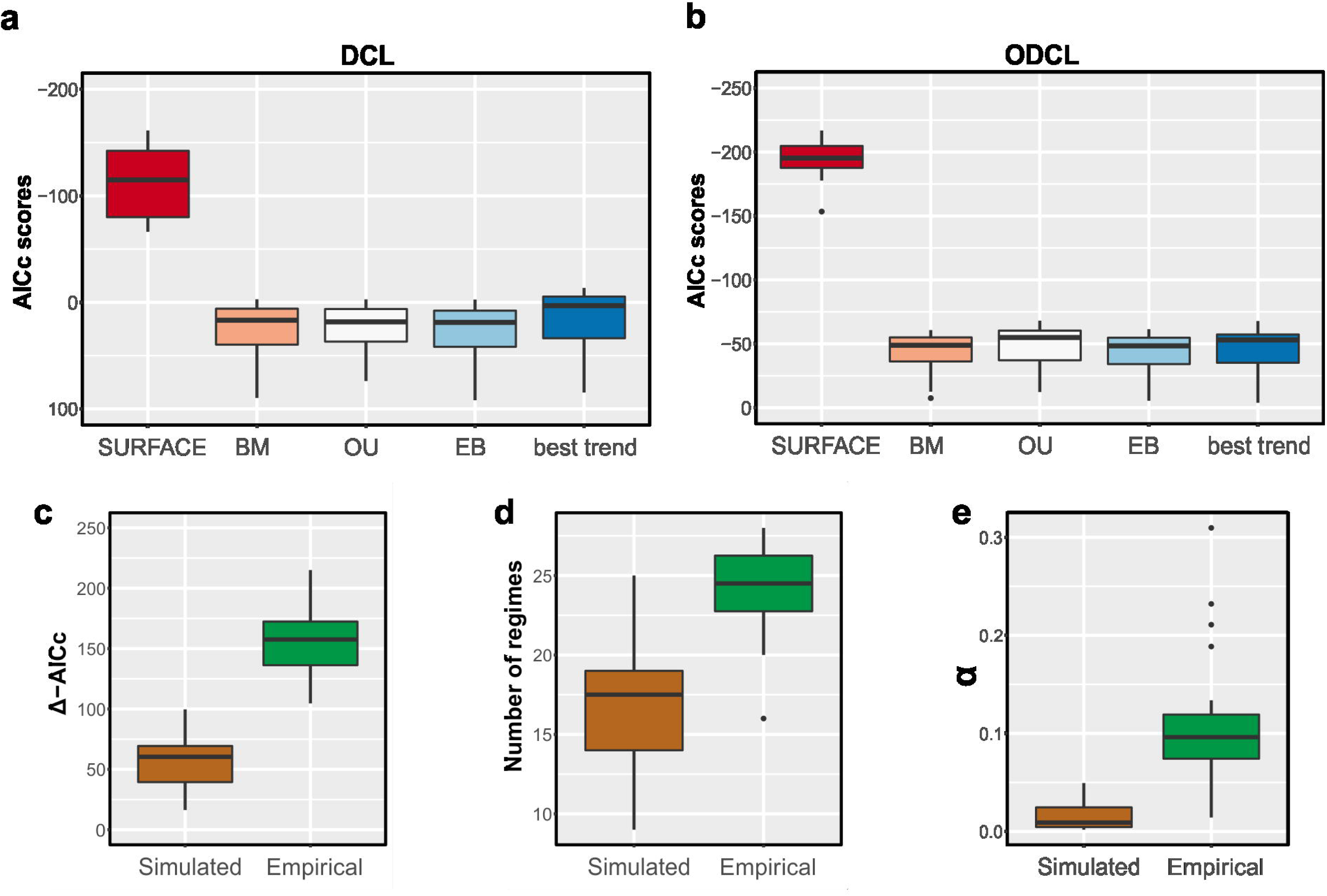
(**a** and **b**) Boxplots showing AICc scores of the evolutionary models fitted to crocodylomorph phylogeny and body size data (using 20 trees time-calibrated with the FBD method). Results shown for two cranial measurements datasets: ODCL (**a**) and DCL (**b**). For the trend-like models, only the AICc of the best model (“best trend”) is shown. See Additional files 1 and 3 for further results. (**c**-**e**) Comparative results of evolutionary models fitted to simulated data (under Brownian Motion) and our empirical body size data (using the ODCL dataset). Data was simulated for 20 crocodylomorph time-scaled trees, and the same trees were used for fitting the evolutionary models. **c** Δ-AICc is the difference between AICc scores received by BM and SURFACE models. **d** Number of regime shifts detected by the SURFACE algorithm. **e** Values of α estimated by the SURFACE algorithm. Results shown for simulated and empirical data.

### Appraising spurious support to the SURFACE model

SURFACE models were generally favoured by AICc compared to a single-regime BM model for our simulated trait data, even though these data were simulated under BM. This is consistent with previous observations of spurious support and high false positive rates for SURFACE models based on stepwise AICc methods [127, 128]. Nevertheless, our results indicate substantially stronger support for SURFACE models based on our empirical data compared to that for the data simulated under BM (Fig. 2a and b). Median delta-AICc between SURFACE and BM models for the simulated data were 60.38, compared to 157.93 for the empirical data, and the distributions of these delta-AICc values are significantly different according to a Wilcoxon–Mann–Whitney test (p < 0.001). Furthermore, the number of regime shifts detected and the values of α estimated are significantly higher (p < 0.001) when using the empirical data (Fig. 2c-e; median values of α estimated of 0.009 and 0.09, for simulated and empirical data, respectively; median number of regimes detected: 17.5 compared to 24.5).

These results suggest that the support for SURFACE models as explanations of our empirical data goes beyond that anticipated simply due to false positives expected for these complex, multi-regime models [127]. Furthermore, the SURFACE model fits represent a useful simplification of major patterns of body size evolution in a group, and particularly the shifts of average body sizes among clades on the phylogeny. Thus, although we acknowledge that some model fits might be suboptimal or could be returning some unrealistic parameter estimates, we use our SURFACE results to provide an overview of crocodylomorph body size evolution that is otherwise lacking from current literature.

### Describing the body size macroevolutionary patterns in Crocodylomorpha

The use of alternative positions of Thalattosuchia (see the “Methods” section) allowed us to further examine the impact of more significant changes to tree topologies on our SURFACE results. In general, similar model configurations were found for all tree topologies (Figs. 3, 4, and 5; see Additional file 4 for all SURFACE plots), with numerous regime shifts detected along crocodylomorph phylogeny. However, simpler model fits (i.e., with significantly less regime shifts) are relatively more frequent when Thalattosuchia is placed as the sister group of Crocodyliformes. To further investigate this, we recalibrated the same tree topologies with other time-scaling methods (i.e., *mbl* and *cal3* methods), and applied SURFACE to those recalibrated trees. Some of these trees returned more complex models, with a greater number of regime shifts and better pBIC scores. This indicates that some of the simpler model configurations might be suboptimal, given that AIC procedures might face difficulties [147], which have previously demonstrated for other datasets (e.g., in dinosaurs [33]).

Overall, most SURFACE model fits identified more than five main macroevolutionary regimes (i.e., “convergent” regimes, identified during the backward-phase of SURFACE), independently of the position of Thalattosuchia (Figs. 3, 4, and 5). Those are distributed along crocodylomorph phylogeny by means of numerous regime shifts, usually more than 20. Trait optima values for these regimes varied significantly among different crocodylomorph subclades and are described in detail below. Overall, regime shifts are frequently detected at the bases of well-recognised clades, such as Thalattosuchia, Notosuchia and Crocodylia. Nevertheless, shifts to new regimes are not restricted to the origins of these diverse clades, since many other regime shifts are observed across crocodylomorph phylogeny, including regimes containing only a single species.

Our SURFACE results indicate an ancestral regime of small body sizes for Crocodylomorpha, regardless of the position of Thalattosuchia (Figs. 3, 4, and 5). This is consistent with the small body sizes of most non-crocodyliform crocodylomorphs such as *Litargosuchus leptorhynchus* and *Hesperosuchus agilis* [50, 51]. The vast majority of the model fits show trait optima for this initial regime (Z_0_) ranging from 60 to 80 cm (total body length was estimated only after the SURFACE model fits, based on the equation from [91]; see the “Methods” section). Very few or no regime shifts are observed among non-crocodyliform crocodylomorphs (Figs. 3, 4, and 5b). The possible exception to this is in Thalattosuchia, members of which occupy large body sized regimes (θ = 500–1000 cm), and which is placed outside Crocodyliformes in some of our phylogenies (Fig. 5a). Regardless of the position of Thalattosuchia, the ancestral regime of all crocodylomorphs (Z_0_) was inherited by protosuchids (such as *Protosuchus*, *Orthosuchus*, and *Edentosuchus*) and some other non-mesoeucrocodylian crocodyliforms (e.g., *Shantungosuchus*, *Fruitachampsa*, *Sichuanosuchus* and *Gobiosuchus*).

Mesoeucrocodylia and *Hsisosuchus* share a new evolutionary regime of slightly larger body sizes (θ = 130–230 cm) in most model fits. This is usually located at the end of the Late Triassic (Rhaetian), and the recovery of this shift is independent of the phylogenetic position of Thalattosuchia (Figs. 3, 4, and 5). The regime that originates at the base of Mesoeucrocodylia (θ = 130–230 cm) is often inherited by Notosuchia and Neosuchia, even though many regime shifts are observed later on during the evolution of these two clades. Within Notosuchia, although some taxa inherit the same regime of smaller sizes present at the base of the clade (θ = 130–230 cm), many regime shifts are also observed (often more than four). Regime shifts to smaller sizes (θ = 60–100 cm) are often seen in uruguaysuchids (including all *Araripesuchus* species), *Anatosuchus*, *Pakasuchus* and *Malawisuchus*. Shifts towards larger sizes are seen among peirosaurids (θ = 210–230 cm) and, more conspicuously, in sebecosuchids and sometimes in the armoured sphagesaurid *Armadillosuchus arrudai* (θ = 330–350 cm).

**Fig. 3.**
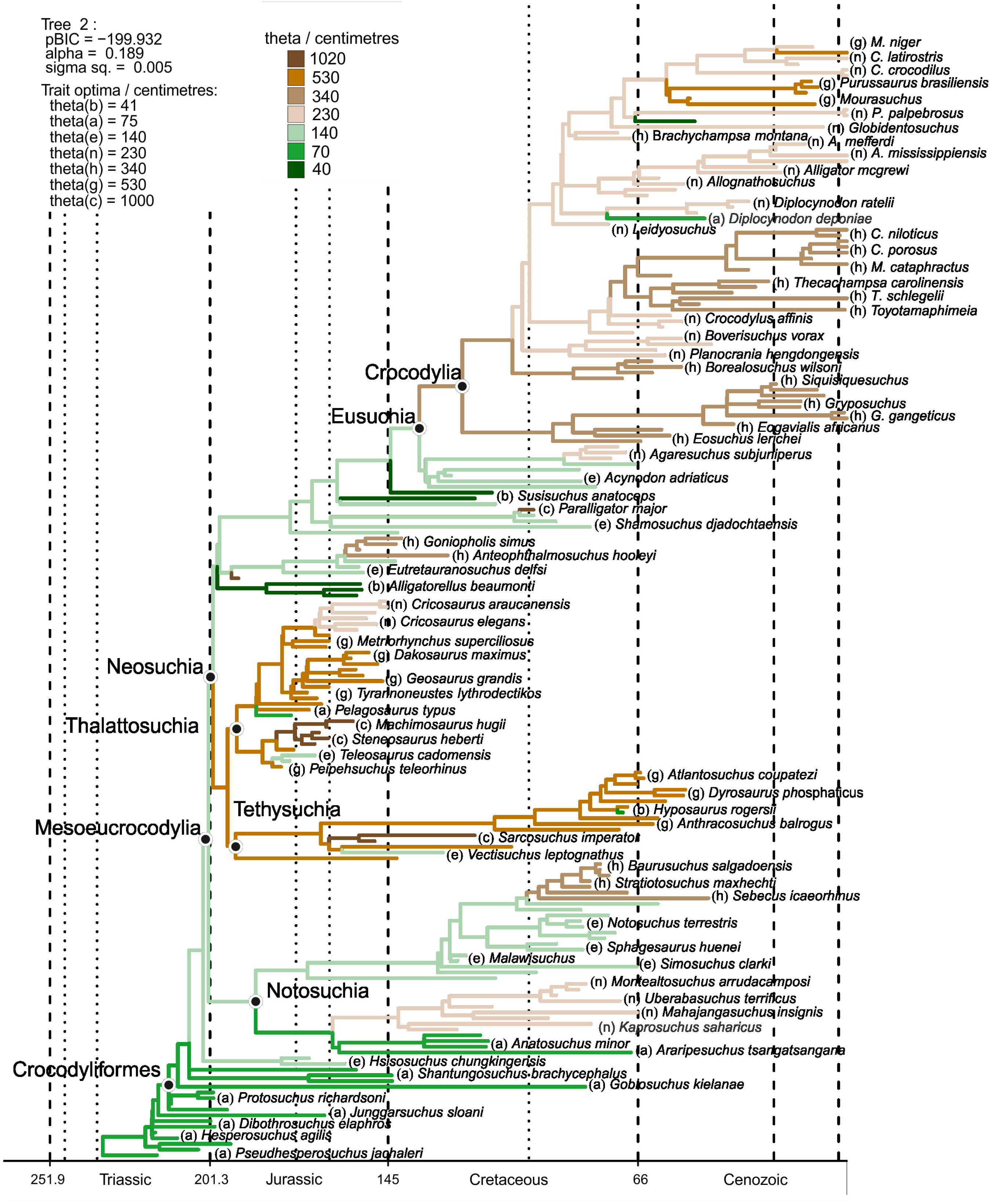
SURFACE model fit (using pBIC searches in the backward-phase) of tree number 2 among crocodylomorph topologies with Thalattosuchia placed within Neosuchia, using the ODCL dataset and time-calibrated with the FBD method. Attraction to unrealized low or high trait optima are highlighted in blue and red, respectively. Model fits of trees sharing the same position of Thalattosuchia show very similar regime configurations, regardless of the dataset used (ODCL or DCL) and the time-calibration method (see Additional file 4 for all SURFACE plots).

**Fig. 4.**
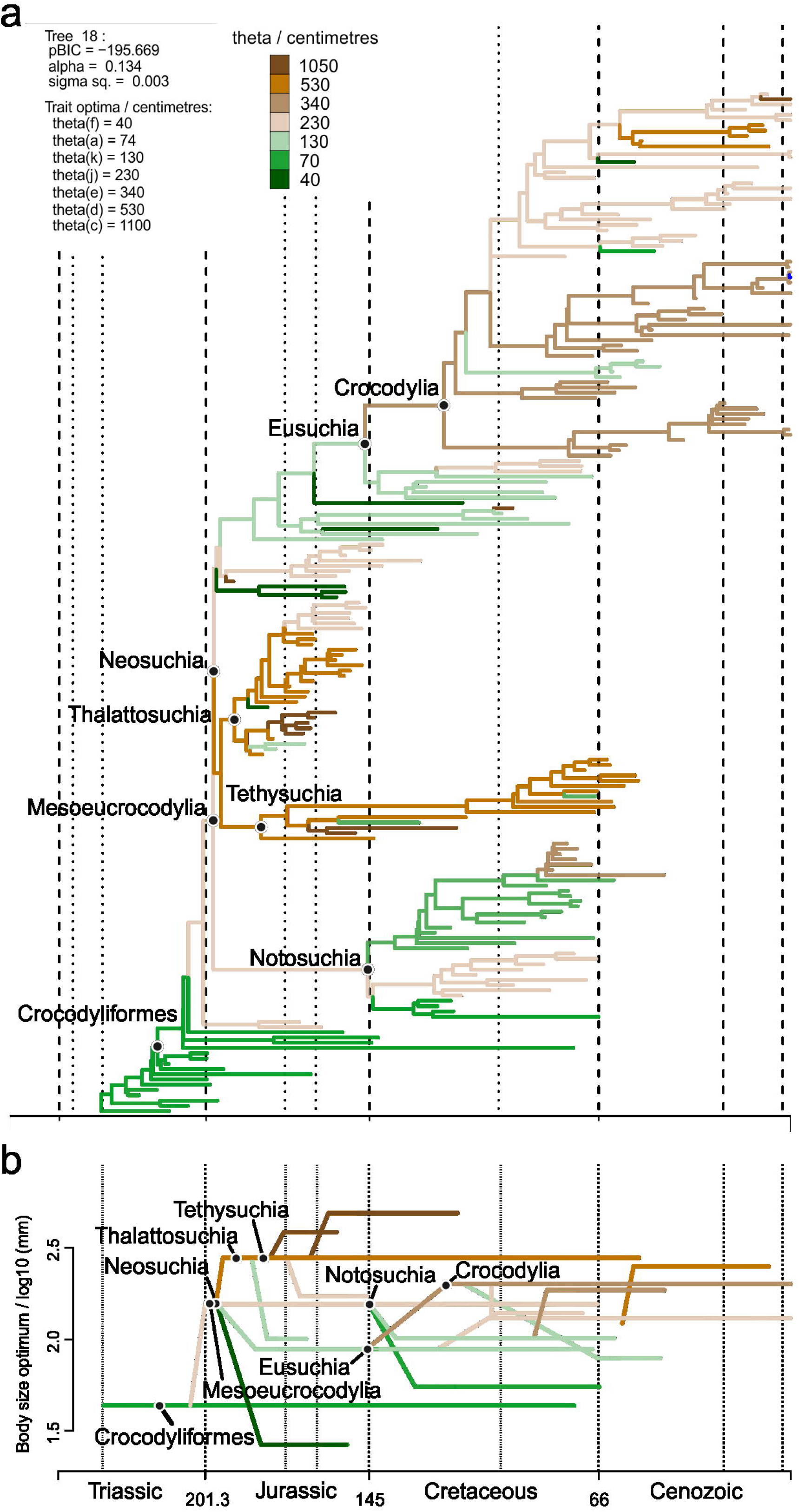
(**a**) SURFACE model fit (using pBIC searches in the backward-phase) of tree number 18 among crocodylomorph topologies with Thalattosuchia placed within Neosuchia, using the ODCL dataset and time-calibrated with the FBD method. Attraction to unrealized low or high trait optima are highlighted in blue and red, respectively. (**b**) Simplified version of **a**, with independent multi-taxon regimes collapsed to single branches.

**Fig. 5.**
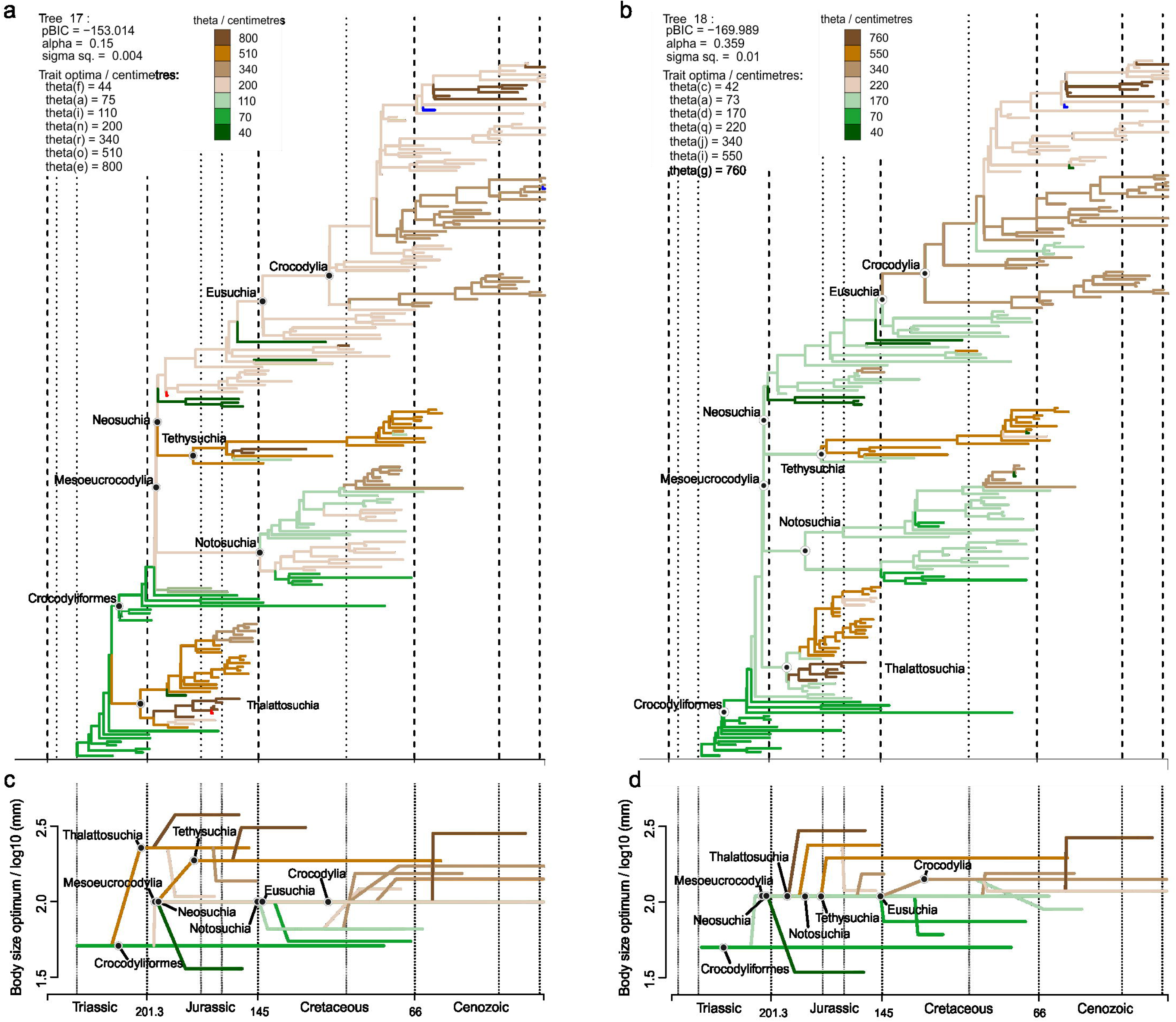
SURFACE model fits of trees time-calibrated with the FBD method, using the ODCL dataset. (**a**) Model fit of tree number 17 with Thalattosuchia as the sister group of Crocodyliformes. Some model fits of trees sharing this same position of Thalattosuchia show simpler model configurations, with significantly fewer regimes (see text for details and Additional file 4 for all SURFACE plots). (**b**) Model fit of tree number 18 with Thalattosuchia as the sister group of Mesoeucrocodylia. (**c** and **d**) Simplified versions of **a** and **b**, respectively, with independent multi-taxon regimes collapsed to single branches.

Independent regime shifts to much smaller sizes (θ = 40–60 cm) are present among non-eusuchian neosuchians (excluding Thalattosuchia and Tethysuchia), particularly in atoposaurids, *Susisuchus*, and *Pietraroiasuchus*, whereas shifts to larger sizes (θ = 300–850 cm) are also detected, often in *Paralligator major* and in some goniopholidids. Within both Tethysuchia and Thalattosuchia, most taxa occupy a regime of relatively large body sizes (θ = 500–1000 cm). When these two clades are sister taxa (Figs. 3 and 4) they usually inherit a same body size regime (θ = 500–550 cm), which originated during the Early Jurassic (Hettangian). In contrast, when Thalattosuchia is placed as sister to Crocodyliformes or Mesoeucrocodylia (Fig. 5), the regime shifts to larger sizes are often independent, and occur at the base of each clade (also with θ values around 500 cm) or later on during their evolutionary history (e.g., some model fits show Tethysuchia with regime shifts to larger sizes only at the base of Dyrosauridae [θ ≈ 500 cm] and the clade formed by *Chalawan* and *Sarcosuchus* [θ = 800–1000 cm]). Both groups also exhibit regime shifts to smaller sizes (θ = 100–150 cm) in some lineages, such as those leading to *Pelagosaurus typus* and *Teleosaurus cadomensis* within Thalattosuchia, and *Vectisuchus* within Tethysuchia. Among thalattosuchians, a conspicuous shift towards larger body sizes (θ = 800–1000 cm) is frequently observed in the teleosaurid clade formed by *Machimosaurus* and *Steneosaurus*, whereas within Metriorhynchidae, a shift to smaller sizes (θ = 230–350 cm) is often detected in Rhacheosaurini.

Crocodylia is also characterized by a predominance of macroevolutionary regimes of relatively large sizes, such as in Thalattosuchia and Tethysuchia. Indeed, regimes of larges sizes are frequently associated with clades of predominantly aquatic or semi-aquatic forms, although not strictly restricted to them. Regarding Crocodylia, a Cretaceous regime shift is usually detected at the base of the clade (Figs. 3, 4, and 5), changing from the macroevolutionary regime of smaller sizes (θ = 130–180 cm) found for non-crocodylian eusuchians (such as hylaeochampsids and some allodaposuchids) to a regime of larger trait optimum (θ = 280–340 cm). This same ancestral regime to all crocodylians is inherited by many members of the clade, particularly within Crocodyloidea and Gavialoidea. Although some model fits show Crocodylia inheriting the same regime as closely related non-crocodylian eusuchians (more frequently when Thalattosuchia is placed outside Neosuchia), shifts towards larger body sizes are seen in members of Crocodyloidea and Gavialoidea, but they only occur later in time and arise independently. In comparison to the other two main lineages of Crocodylia, Alligatoroidea is characterized by a regime of lower trait optima values (θ = 210–230 cm), which frequently occurs as a Late Cretaceous shift at the base of the clade. But Alligatoroidea is also distinct from the other two clades by exhibiting more regime shifts, reflecting its great ecological diversity and body size disparity (ranging from very small taxa, such as the caimanine *Tsoabichi greenriverensis*, to the huge *Purussaurus* and *Mourasuchus*).

### Modes of body size evolution within Notosuchia and Crocodylia

The significant number of regime shifts that occur within both Notosuchia and Crocodylia led us to more deeply scrutinise the modes of body size evolution in these two clades. We therefore conducted another round of model-fitting analyses, initially fitting the same evolutionary models (SURFACE, OU, BM, EB and trend-like models) to subtrees representing both groups. In addition, we used the same regime shifts identified by the SURFACE algorithm to fit four additional models using the OUwie algorithm (BMS, OUMV, OUMA and OUMVA), which allow more parameters to vary, but need regime shifts to be set *a priori*.

The results of these analyses indicate different modes of body size evolution during the evolutionary histories of these two groups. In Crocodylia (Fig. 6; see Additional file 3 for a complete list of AICc scores), AICc scores indicate a clear preference for OU-based models, with highest support found for the SURFACE model, but also strong support for the uniform OU model, as well as OUMA and OUMVA models. The SURFACE algorithm frequently identified at least three main (i.e. “convergent”) macroevolutionary regimes for crocodylians (with θ values around 200, 350 and 750 cm, respectively), usually with α ranging from 0.02 to 0.2 and σ^2^ between 0.0007 and 0.02. When allowed to vary among regimes (i.e., in models OUMA and OUMVA), ranges of both parameters increase significantly, with some model fits displaying extremely unrealistic parameter values, which might explain the stronger support found for SURFACE compared to these latter models. Even though the relatively small number of taxa included in these analyses (i.e. N = 70) suggests caution when interpreting the higher support for OU-based models [128], BM-based models received consistently worse support than any of the four OU-based models mentioned above, even the best trend-like model (usually the one with the best AICc scores among BM-based models).

**Fig. 6.**
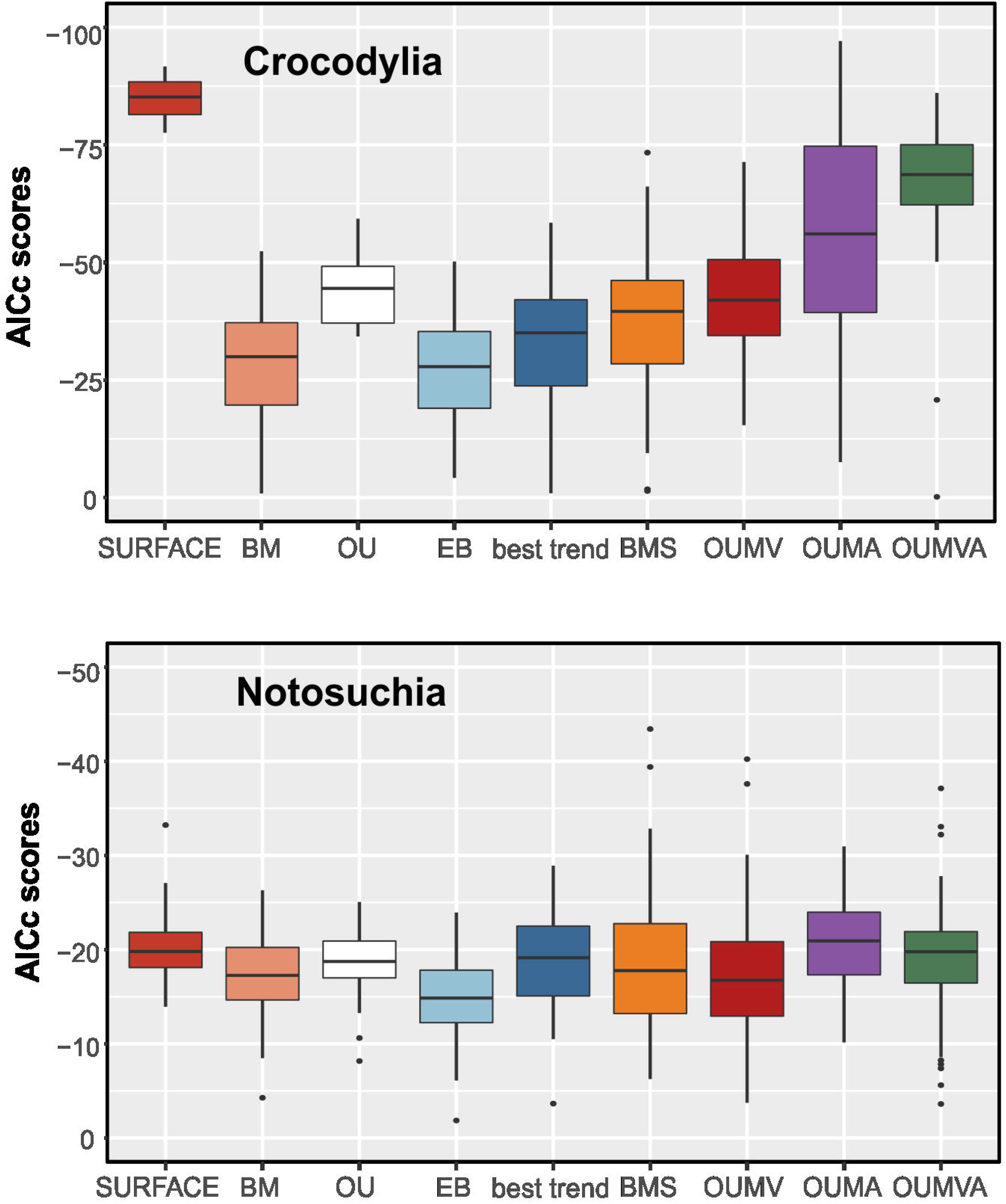
AICc scores of all evolutionary models fitted to the phylogenies and body size data of Crocodylia (top) and Notosuchia (bottom). For the trend-like models, only the AICc of the best model (“best trend”) is shown.

Our results show a different scenario for Notosuchia, for which we found comparable support for all evolutionary models analysed (Fig. 6). Among OU-based models, slightly better AICc scores were found for the SURFACE model. However, this model received virtually the same support as the BMS model, the best of the BM-based models. BMS is a multi-regime BM model that allows the rate parameter (σ^2^) to vary, and, as α is effectively set to zero, represents diffusive model of evolution. The support found for this model might suggest a more relaxed mode of body size evolution in notosuchians, which is consistent with the wide range of body sizes observed in the group, even among closely-related taxa. Although OU-based models (including SURFACE) are not favoured over other evolutionary models, we can use some SURFACE model to further explore body size evolutionary patterns among Notosuchia. For example, even though we sampled twice as many crocodylians (N = 70) as notosuchians (N = 34), many SURFACE model fits found three main macroevolutionary regimes for notosuchians, similar to what was found for Crocodylia (although model fits with less regimes were more frequent for Notosuchia than Crocodylia). For these, θ values were usually around 80, 150 and 320 cm, with α usually ranging from 0.008 to 0.05 and σ^2^ between 0.0007 and 0.005. When the same regimes detected by the SURFACE algorithm were used by the OUwie algorithm to fit the BMS model, values of σ^2^ rarely varied significantly from the range of whole-tree σ^2^ estimated for the SURFACE model fits. The few exceptions were usually related to regimes with unrealised θ values, as in the case of the armoured sphagesaurid *Armadillosuchus arrudai* (probably with more than 2 metres in total length, whereas other sampled sphagesaurids reach no more than 1.2 m [148]), and sebecosuchians (top predators of usually more than 2.5 metres [97]), even though these values might still be realistic when simulating trend-like dynamics (i.e., in a single lineage with extremely disparate trait values [19, 57]).

### The influence of palaeolatitude and palaeotemperature

Most of the correlation analyses between our body size data and the different datasets of the abiotic factors palaeotemperature and palaeolatitude yielded weak (coefficient of determination R^2^ usually smaller than 0.2) or non-significant correlations (see Additional file 1 for all regressions and further results). This is consistent with the distribution of crocodylomorph body size through time (Fig. 7), as well as with the results from our macroevolutionary analyses, which found strong support for a multi-regime OU model (SURFACE). This suggests that shifts between macroevolutionary regimes (which we interpret as “maximum adaptive zones” sensu Stanley [11]) are more important in determining large-scale macroevolutionary patterns of crocodylomorph body size evolution than these abiotic factors, at least when analysed separately.

**Fig. 7.**
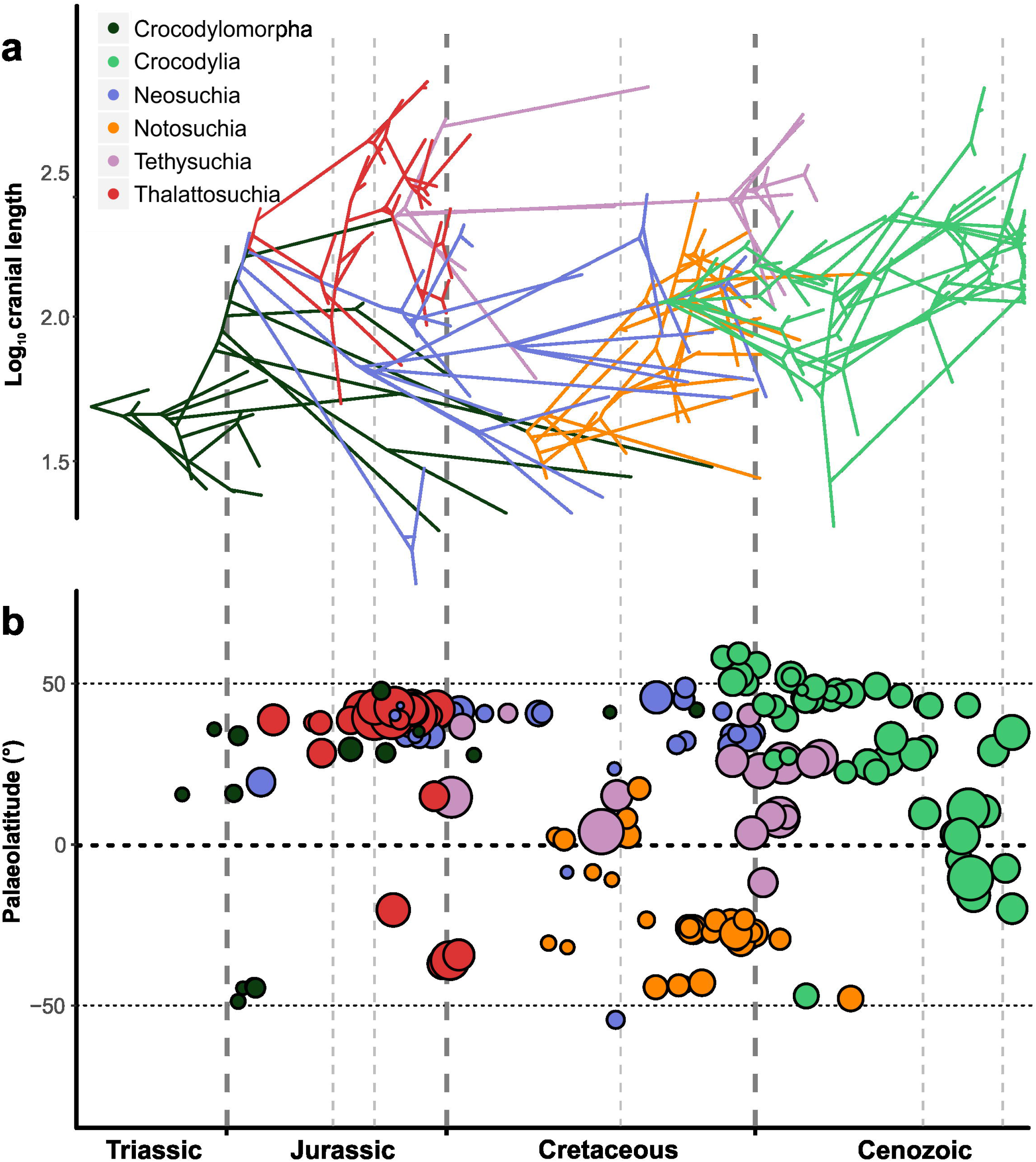
Crocodylomorph body size through time, with colours representing different mono- or paraphyletic (i.e., Crocodylomorph = non-mesoeucrocodylian crocodylomorphs, excluding Thalattosuchia; Neosuchia = non-crocodylian neosuchians) crocodylomorph groups. Body size represented by log_10_ ODCL (orbito-cranial dorsal length) in millimetres. (**a**) Phenogram with body size incorporated into crocodylomorph phylogeny. (**b**) Palaeolatitudinal distribution of extinct crocodylomorphs through time, incorporating body size information (i.e., different-sized circles represent variation in body size).

However, one important exception was found: a correlation between mean body size values and palaeotemperatures from the Late Cretaceous (Maastrichtian) to the Recent (data from [131]). Using either all taxa in the datasets or only non-marine species, we found moderate to strong correlations (R^2^ ranging from 0.376 to 0.635), with higher mean body size values found in time intervals with lower temperatures (i.e., positive slopes, given that the δ^18^O proxy is inversely proportional to temperature). The correlation was present even when we applied GLS regressions with an autoregressive model (Table 1), which returned near-zero or low autocorrelation coefficients (phi = 0.01–0.15). This suggests that temperature might have had an influence in determining the body size distribution of crocodylomorphs at smaller temporal and phylogenetic scales. For this reason, we decided to further scrutinise the relationships between the distribution of body sizes and these abiotic factors at these smaller scales, repeating our regression analyses using only data for Crocodylia, Notosuchia, Thalattosuchia, and Tethysuchia (see the “Methods” section).

**Table 1.** Regression results of mean values of body size values on palaeotemperature.

| Dataset | GLS |  |  |  | OLS (untransformed) |  |  |  |
| --- | --- | --- | --- | --- | --- | --- | --- | --- |
|  | Phi | Intercept | Slope | AIC | R<sup>2</sup> | Intercept | Slope | AIC |
| **ODCL**<br>**with all taxa** | -0.046 | 2.022 | 0.055<br>(0.002) | -31.576 | 0.635 | 2.023 | 0.054<br>(0.003) | -33.557 |
| **DCL**<br>**with all taxa** | 0.014 | 2.433 | 0.081<br>(0.011) | -19.577 | 0.527 | 2.433 | 0.081<br>(0.01) | -21.575 |
| **ODCL**<br>**non-marine** | -0.157 | 1.964 | 0.06<br>(0.007) | -24.96 | 0.502 | 1.965 | 0.06<br>(0.013) | -26.706 |
| **DCL**<br>**non-marine** | -0.089 | 2.345 | 0.07<br>(0.027) | -16.045 | 0.376 | 2.346 | 0.07<br>(0.034) | -18.272 |

These additional regressions corroborate the hypothesis that at least some crocodylomorph subclades show a correspondence between body size and global palaeotemperature. Although most of the regressions provided non-significant or weak/very weak correlations (see Additional file 1 for all regression results), including all regressions of body size on palaeolatitudinal data, both maximum and mean body size values of Crocodylia are moderately to strongly correlated to palaeotemperature through time (Table 2). The positive slopes and coefficients of determination (R^2^ ranging from 0.554 to 0.698) indicate that the lowest temperatures are associated with the highest body size values in the crown-group. However, correlations with data from other subclades (Notosuchia, Thalattosuchia and Tethysuchia) were mostly non-significant, suggesting that this relationship between body size and temperature was not a widespread pattern among all groups.

**Table 2.**
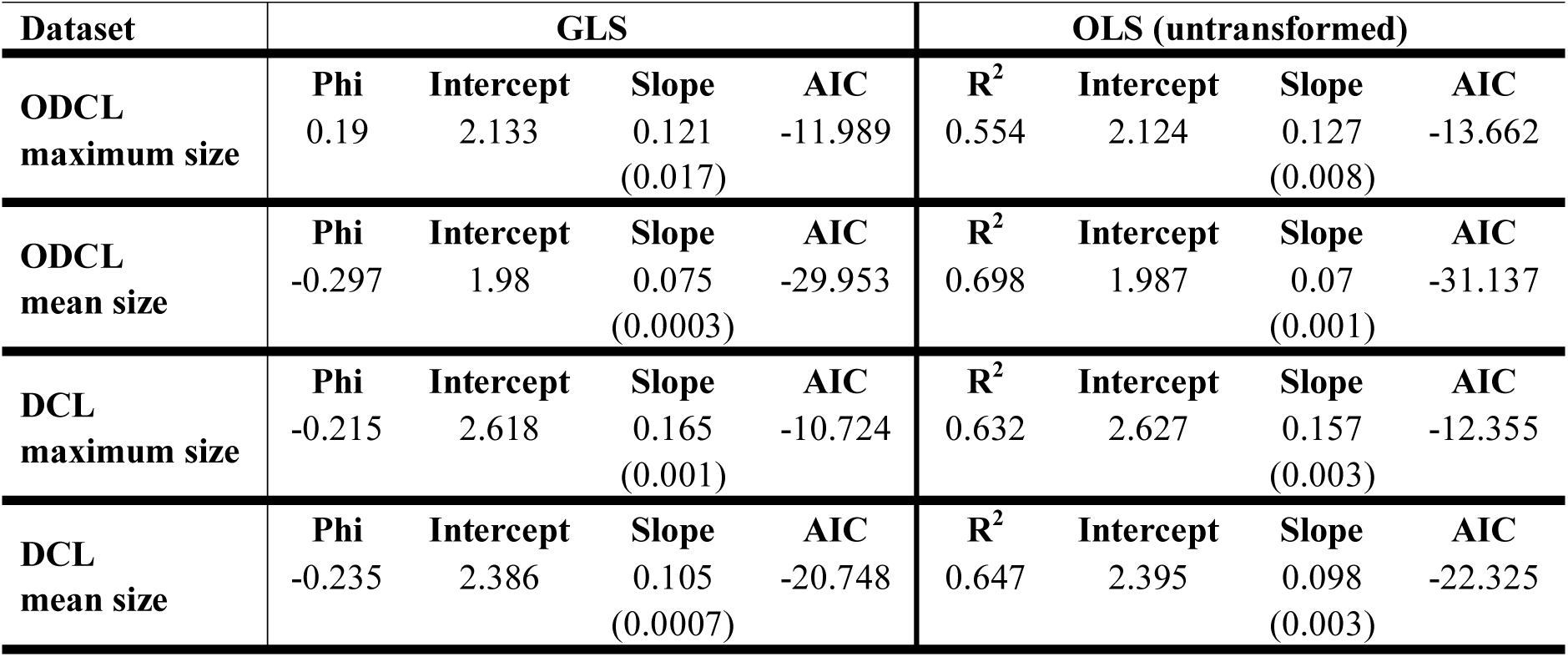
Regression results of maximum and mean crocodylian body size values on palaeotemperature.

| Dataset | GLS |  |  |  | OLS (untransformed) |  |  |  |
| --- | --- | --- | --- | --- | --- | --- | --- | --- |
|  | Phi | Intercept | Slope | AIC | R<sup>2</sup> | Intercept | Slope | AIC |
| **ODCL**<br>**maximum size** | 0.19 | 2.133 | 0.121<br>(0.017) | -11.989 | 0.554 | 2.124 | 0.127<br>(0.008) | -13.662 |
| **ODCL**<br>**mean size** | -0.297 | 1.98 | 0.075<br>(0.0003) | -29.953 | 0.698 | 1.987 | 0.07<br>(0.001) | -31.137 |
| **DCL**<br>**maximum size** | -0.215 | 2.618 | 0.165<br>(0.001) | -10.724 | 0.632 | 2.627 | 0.157<br>(0.003) | -12.355 |
| **DCL**<br>**mean size** | -0.235 | 2.386 | 0.105<br>(0.0007) | -20.748 | 0.647 | 2.395 | 0.098<br>(0.003) | -22.325 |

Results of GLS (with an autoregressive model) and OLS (untransformed data) regressions. Mean body size represented by mean values of log-transformed cranial measurements (DCL and ODCL), in millimetres. Data from both ODCL and DCL datasets was divided into subsets with all crocodylomorphs or only non-marine species. N = 10 in all four subsets (number of time bins analysed). Palaeotemperature data from [131], represented by δ^18^O data from the Late Cretaceous to Recent. Only significant correlations (p < 0.05) are shown.

Results of GLS (with an autoregressive model) and OLS (untransformed data) regressions. Mean and maximum body size only for members of the crown-group Crocodylia, represented by mean and maximum values of log-transformed cranial measurements (DCL and ODCL), in millimetres. N = 10 in all four datasets (number of time bins analysed). Palaeotemperature data from [131], represented by δ^18^O data from the Late Cretaceous to Recent. Only significant correlations (p < 0.05) are shown.

### Correlation between body size and habitat choice

We initially found a relationship between lifestyle (i.e., terrestrial, semi-aquatic/freshwater, and aquatic/marine) and body size using ANOVA. However, a phylogenetic ANOVA [140] returned non-significant results (Table 3). Phylogenetic ANOVA asks specifically whether evolutionary habitat transitions are consistently associated with particular body size shifts as optimised on the phylogeny. This indicates that, although crocodylomorphs with more aquatic lifestyles (particularly marine species) tend to be large-bodied, the evolutionary transitions between these lifestyle categories were probably not accompanied by immediate non-random size changes. Furthermore, the smaller body sizes of some aquatic or semi-aquatic lineages (e.g., atoposaurids, *Tsoabichi* and *Pelagosaurus*) show that adaptive peaks of smaller sizes are also viable among aquatic/semi-aquatic species. This suggests that, even though there seems to be an ecological advantage for larger-sized freshwater/marine crocodylomorphs, the lower limit of body size in aquatic species was comparable to that of terrestrial species.

**Table 3.**
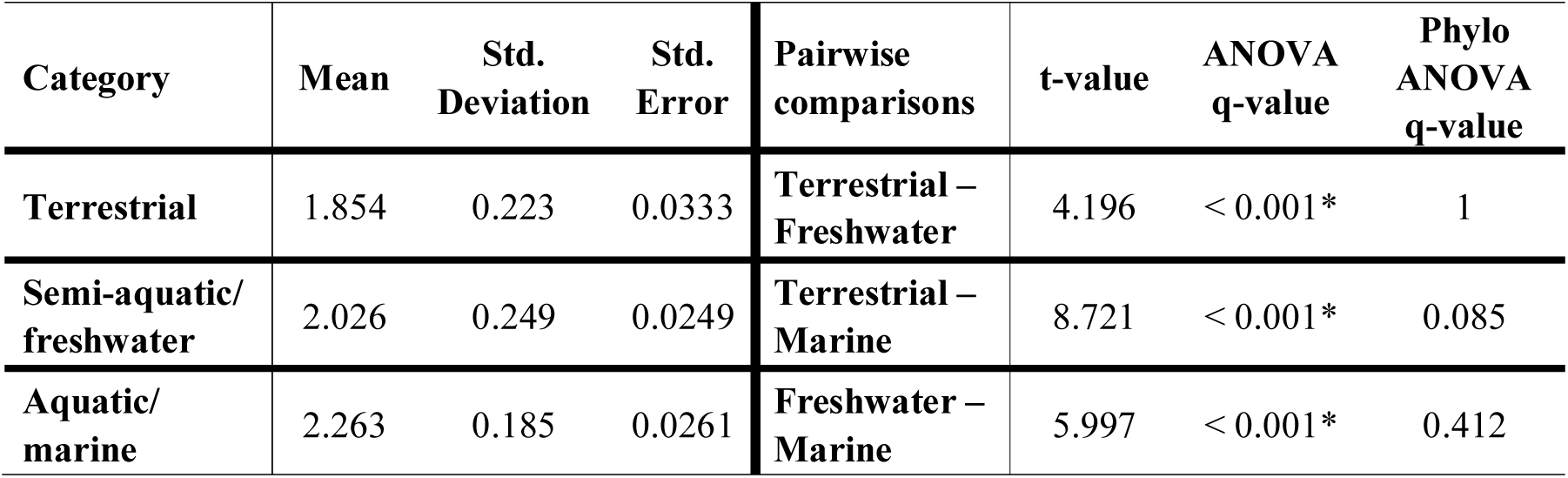
Pairwise comparison between body size of crocodylomorphs subdivided into three lifestyle categories.

| Category | Mean | Std. Deviation | Std. Error | Pairwise comparisons | t-value | ANOVA q-value | Phylo ANOVA q-value |
| --- | --- | --- | --- | --- | --- | --- | --- |
| Terrestrial | 1.854 | 0.223 | 0.0333 | Terrestrial – Freshwater | 4.196 | < 0.001* | 1 |
| Semi-aquatic/freshwater | 2.026 | 0.249 | 0.0249 | Terrestrial – Marine | 8.721 | < 0.001* | 0.085 |
| Aquatic/marine | 2.263 | 0.185 | 0.0261 | Freshwater – Marine | 5.997 | < 0.001* | 0.412 |

Body size data from the ODCL dataset (log-transformed cranial measurement, in millimetres). Number of species in each category: 45 (terrestrial), 100 (semi-aquatic/freshwater), and 50 (aquatic/marine). Results from ANOVA, without accounting for phylogenetic dependency, and phylogenetic ANOVA [140] with 100,000 simulations. *Bonferroni-corrected p-values (q- values) significant at alpha = 0.05

## Discussion

### The adaptive landscape of crocodylomorph body size evolution

Crocodylomorph body size disparity increased rapidly during the early evolution of the group, from the Late Triassic to the Early Jurassic (Hettangian–Sinemurian), which is mostly a result of the appearance of the large-bodied thalattosuchians (Fig. 8b). After a decline in the Middle Jurassic, body size disparity reaches its maximum peak in the Late Jurassic, with the appearance of atoposaurids, some of the smallest crocodylomorphs, as well as large teleosaurids (such as *Machimosaurus*). This increase in disparity may have occurred earlier than our results suggest, given that Middle Jurassic records of atoposaurids [149] could not be included in our analyses due to their highly incomplete preservation.

**Fig. 8.**
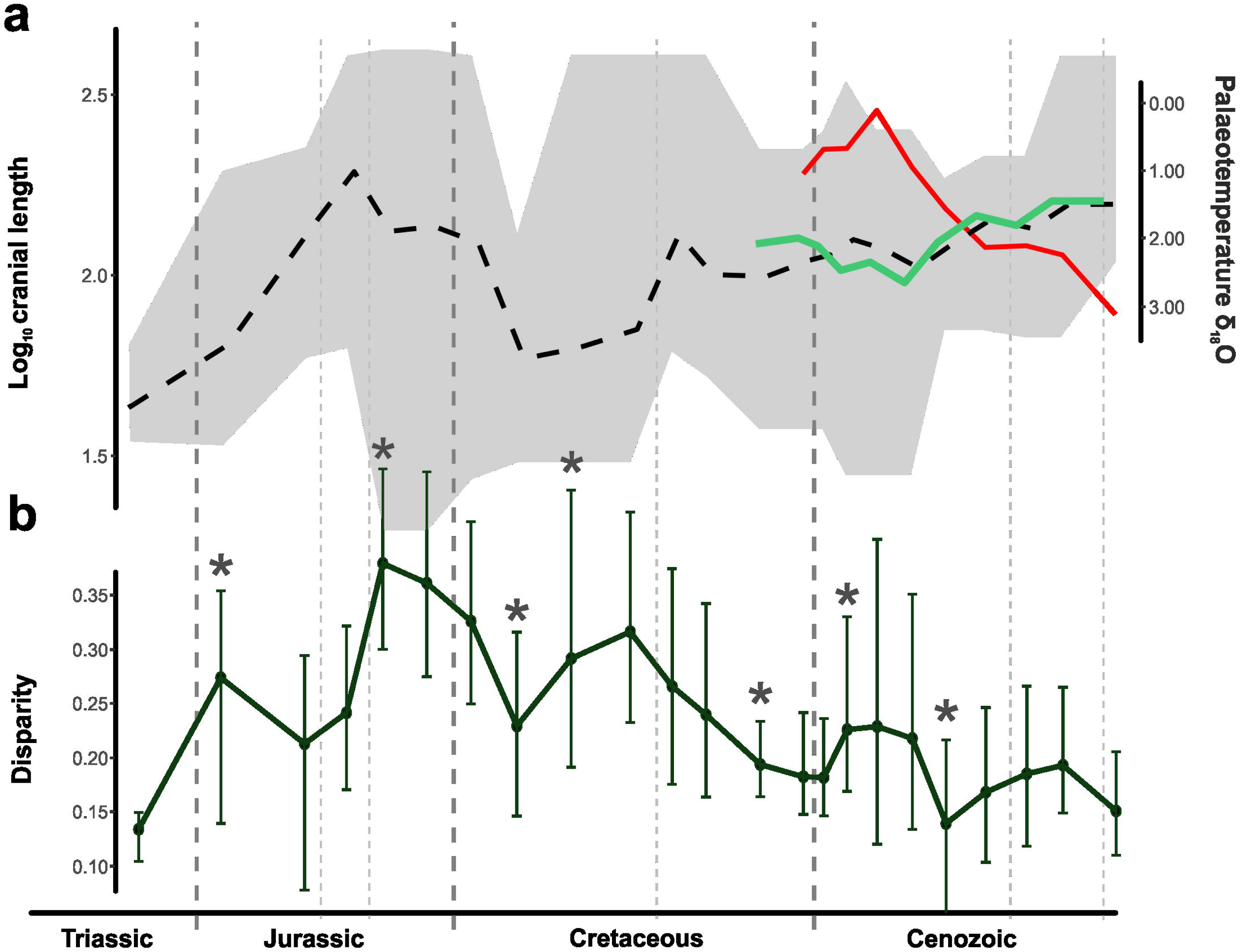
(**a**) Crocodylomorph body size and palaeotemperature through time. Mean log_10_ ODCL represented by dashed black line, shaded polygon shows maximum and minimum values for each time bin. Continuous light green displays mean log_10_ ODCL values only for Crocodylia. Palaeotemperature (δ^18^O) illustrated by red line (data from [132]). (**b**) Body size disparity through time. Disparity is represented by the standard deviation of log_10_ ODCL values for each time bin (only time bins with more than 3 taxa were used for calculating disparity). Error bars are accelerated bias-corrected percentile limits (BCa) of disparity from 1,000 bootstrapping replicates. Asterisks mark the events of largest interval-to-interval changes in disparity.

Since this peak in the Middle/Late Jurassic, crocodylomorphs underwent an essentially continuous decline in body size disparity, with some short-term fluctuations related to the extinction or diversification of particular lineages. The Early Cretaceous witnessed the extinction of thalattosuchians, and a sharp decrease in disparity is seen from the Berriasian to the Barremian (although this time interval is also relatively poorly sampled in our dataset). A subsequent increase in disparity is seen in the Aptian, probably reflecting the appearance of small-bodied crocodylomorphs (such as susisuchid eusuchians). Nevertheless, this is followed by a continuing decline for the remainder of the Cretaceous (in spite of the occurrence of highly disparate notosuchians). The Cenozoic is also characterised by an overall decrease in disparity, even though some short-term increases in disparity do occur, mostly related to the presence of smaller-bodied crocodylians in the Palaeogene (such as *Tsoabichi* [150]).

We characterised the macroevolutionary patterns that gave rise to these patterns of body size disparity through time, by performing comparative model-fitting analyses. Our results indicate a strong support found for a multi-peak OU model (i.e., the SURFACE model; Fig. 2a and b). Within the concept of adaptive landscape [73, 74, 75], we can interpret the SURFACE regimes, with different trait optima, as similar to shifts to new macroevolutionary adaptive zones [11, 151]. Thus, the support found for the SURFACE model indicates that lineage-specific adaptations related to body size play an important role in determining the patterns of crocodylomorph body size evolution. Our comparative model-fitting analyses also indicate that uniform OU models, BM models, and both uniform and multi-regime trend models provide poor explanations for the overall patterns of crocodylomorph body size evolution.

Our findings reject the hypothesis of long-term, multi-lineage trends during the evolution of crocodylomorph body size. This is true even for Crocodylia, which shows increases in maximum, minimum and mean body sizes during the past 70 million years (Fig. 8a), a pattern that is classically taken as evidence for trend-like dynamics [56]. In fact, explicitly phylogenetic models of the dynamics along evolving lineages reject this.

We can also reject diffusive, unconstrained Brownian-motion like dynamics for most of Crocodylomorpha, although Notosuchia might be characterised by relatively unconstrained dynamics (Fig. 6). Single-regime (=uniform) models received poor support in general, which might be expected for long-lived and disparate clades such as Crocodylomorpha, which show complex and non-uniform patterns of body size evolution (see [5, 11, 58, 61]). Although multi-regime trend-like models received stronger support than uniform models for most phylogenies (Fig. 2a and b), multi-peak OU models (SURFACE) received overwhelmingly still greater support. This suggests that the macroevolutionary landscape of crocodylomorph body size evolution is best described by shifts between phylogenetically defined regimes that experience constrained evolution around distinct trait optima [61, 71, 75, 83].

The success of a multi-peak OU model indicates that, in general, a significant amount of crocodylomorph body size variance emerged through pulses of body size variation, and not from a gradual, BM-based dispersal of lineages through trait (body size) space. These pulses, represented by regime shifts, represent excursions of single phylogenetic lineages through body size space, resulting in the founding of new clades with distinct body size from their ancestors. This indicates that lineage-specific adaptations (such as those related to ecological diversification; see below) are an important aspect of the large-scale patterns of crocodylomorph body size evolution.

This can also explain the weak support found for the early burst (EB) model in our analyses. The early burst model attempts to simulate Simpson’s [73] idea of diversification through “invasion” of new adaptive zones (niche-filling). It focuses on a particular pattern of adaptive radiation, with evolutionary rates higher in the early evolution of a clade and decelerating through time [126]. Other models have also been proposed to better represent the concept of pulsed Simpsonian evolution (e.g., [152]).

Our results show that, overall, the EB model offers a poor explanation for the evolution of body size in crocodylomorphs, in agreement with previous works that suggested that early bursts of animal body size receive little support from phylogenetic comparative methods ([126], but see [153] for intrinsic issues for detecting early bursts from extant-only datasets). However, rejection of an early burst model does not reject Simpson’s hypothesis that abrupt phenotypic shifts along evolving lineages (“quantum evolution”) results from the distribution of opportunities (adaptive zones, or unfilled niches). Patterns of crocodylomorph body size evolution could still be explained by this “niche-filling” process if opportunities were distributed through time rather than being concentrated early on the evolution of the clade. This is one possible explanation of the pattern of regime shifts returned by our analyses, and might be particularly relevant for clades with long evolutionary histories spanning many geological intervals and undergoing many episodes of radiation.

Bronzati et al. [35] examined variation in rates of species diversification among clades using methods based on tree asymmetry. They found that most of crocodyliform diversity was achieved by a small number of significant diversification events that were mostly linked to the origin of some subclades, rather than via a continuous process through time. Some of the diversification shifts from Bronzati et al. [35] coincide with body size regime shifts found in many of our SURFACE model fits (such as at the base of Notosuchia, Eusuchia and Alligatoroidea; Fig. 9). However, many of the shifts in body size regimes detected by our analyses are found in less-inclusive groups (as in the case of “singleton” regimes, that contain only a single taxon).

**Fig. 9.**
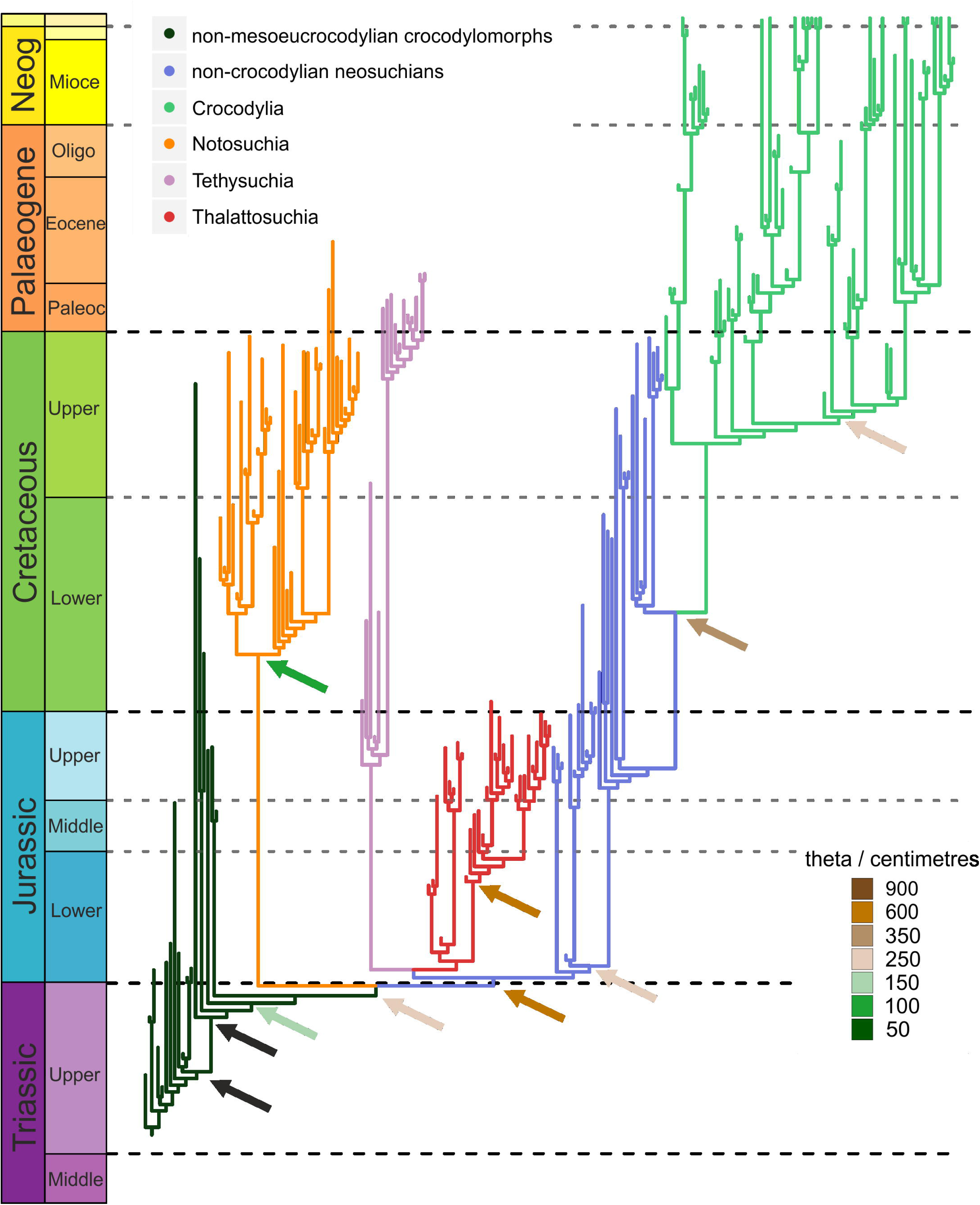
Summary of our SURFACE results combined with the crocodylomorph diversification shifts found by Bronzati et al. [35]. Nodes with diversification shifts are indicated by arrows, the colours of which represent distinct trait optima values (total body length in centimetres, after applying formula from [91]), of different body size regimes. Black arrows indicate nodes for which diversification shifts were identified, but no body size regime shift was found by any of our SURFACE model fits.

### Ecological diversification and its implications for crocodylomorph body size distribution

Ecological factors seem to be important for the large-scale patterns of body size in crocodylomorphs. Many of the regime shifts to larger sizes detected by our SURFACE analyses occur at the base of predominantly aquatic or semi-aquatic clades, such as Thalattosuchia, Tethysuchia and Crocodylia (Figs. 3, 4, and 5), although small-bodied aquatic/semi-aquatic clades also occur, such as Atoposauridae. Some terrestrial clades also display relatively large sizes (such as sebecosuchians and peirosaurids, within Notosuchia). However, most terrestrial species are small-bodied (Fig. 10b), including many of the earliest crocodylomorphs (such as *Litargosuchus leptorhynchus* and *Hesperosuchus agilis* [50, 51]; Fig. 10a), and are within body size regimes of lower values of θ (< 150 cm; Figs. 3, 4, and 5). In contrast, the regimes with the highest values of θ (> 800 cm) are almost always associated with aquatic or semi-aquatic crocodylomorphs (e.g., the tethysuchians *Sarcosuchus imperator* and *Chalawan thailandicus*, the thalattosuchians *Machimosaurus* and *Steneosaurus*, and the crocodylians *Purussaurus* and *Mourasuchus*).

**Fig. 10.**
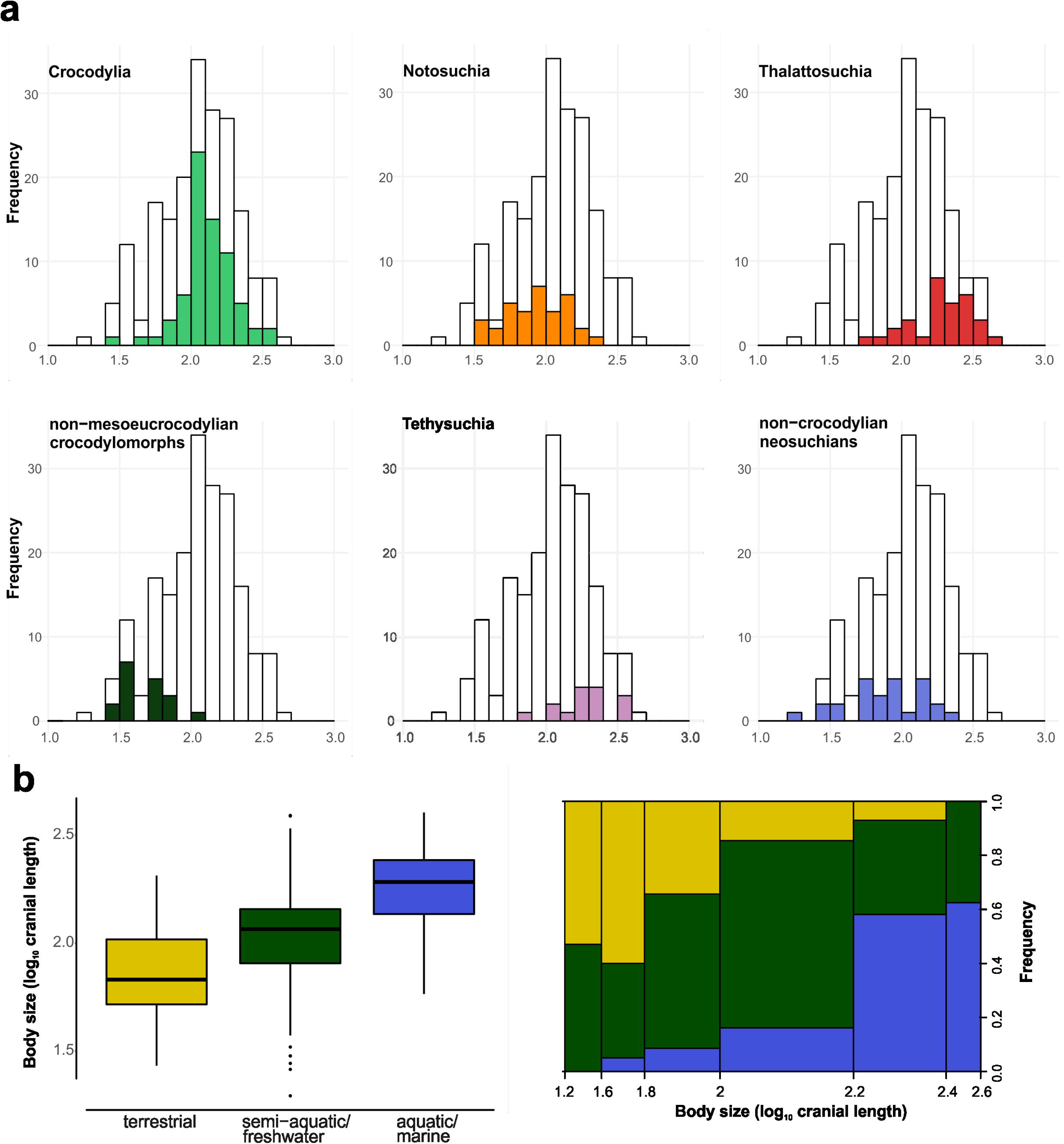
(a) Body size frequency distributions of different crocodylomorph groups (mono- or paraphyletic), constructed using the full set of 240 specimens in the ODCL dataset. Underlying unfilled bars represent values for all crocodylomorphs. Filled bars represent values for Crocodylia, Notosuchia, Thalattosuchia, non-mesoeucrocodylian crocodylomorphs (excluding thalattosuchians), Tethysuchia and non-crocodylian neosuchians (excluding tethysuchians and thalattosuchians). (**b**) Body size distributions of different crocodylomorph lifestyles, shown with box-and-whisker plots (on the left) and a mosaic plot (on the right). The 195 species from the ODCL dataset were subdivided into terrestrial, semi-aquatic/freshwater and aquatic/marine categories (N = 45, 100 and 50, respectively) based on the literature. Body size is represented by log_10_ cranial length (ODCL, orbito-cranial length, in millimetres).

Previous studies have investigated a possible link between an aquatic/marine lifestyle and larger body sizes in other animals, particularly in mammals (e.g., [17, 21, 24]). For instance, it has been previously shown that aquatic life in mammals imposes a limit to minimum body size [24, 154] and relaxes constraints on maximum size [155]. Therefore, aquatic mammals (especially marine ones) have larger body sizes than their terrestrial relatives [21, 156]. We document a similar pattern in crocodylomorphs (Table 3), although the phylogenetic ANOVA results revealed that changes in size are not abrupt after environmental invasions (as also suggested by the diminutive size of some semiaquatic lineages, such as atoposaurids and some crocodylians). Animals lose heat faster in water than in air (given the different rates of convective heat loss in these two environments), and it has demonstrated that thermoregulation plays an important role in determining the larger sizes of aquatic mammals [24, 154, 157]. Although mammals have distinct thermal physiology to crocodylomorphs (which are ectothermic poikilotherms), it has been reported that American alligators (*Alligator mississippiensis*) heat up more rapidly than cool down, and that larger individuals are able to maintain their inner temperature for longer than smaller ones [158]. Thus, given that both heating and cooling rates are higher in water than in air [158], larger aquatic/semi-aquatic animals could have advantages in terms of physiological thermoregulation. If extinct crocodylomorphs had similar physiologies, this could provide a plausible explanation for the larger sizes of non-terrestrial species.

### Cope’s rule cannot explain the evolution of larger sizes in Crocodylomorpha

Previous interpretations of the fossil record suggest a dominance of small sizes during the early evolution of crocodylomorphs [45, 117], inferred from the small body sizes of most early crocodylomorphs. Consistent with this, our SURFACE results revealed a small-bodied ancestral regime for Crocodylomorpha (Z_0_ between 66 and 100 cm), which was inherited virtually by all non-crocodyliform crocodylomorphs. Larger non-crocodyliform crocodylomorphs have also been reported for the Late Triassic (e.g., *Carnufex carolinensis* and *Redondavenator quayensis*, with estimated body lengths of approximately 3 metres [159]), but the fragmentary nature of their specimens prevented us from including them in our macroevolutionary analysis. Nevertheless, given the larger numbers of small-bodied early crocodylomorphs, taxa like *Carnufex* and *Redondavenator* probably represent derived origins of large body size and their inclusion would likely result in similar values of ancestral trait optima (=Z_0_).

The small ancestral body size inferred for crocodylomorphs, combined with the much larger sizes seen in most extant crocodylians and in some other crocodylomorph subclades (such as thalattosuchians and tethysuchians), suggests a pattern of increasing average body size during crocodylomorph evolutionary history. This idea is reinforced by the overall increase in crocodylomorph mean body size through time, particularly after the Early Cretaceous (Fig. 8a). The same pattern also occurs within Crocodylia during the past 70 million years (Fig. 8a), as some of the earliest taxa (such as *Tsoabichi*, *Wannaganosuchus* and *Diplocynodon deponiae*) were smaller-bodied (< 2m) than more recent species, such as most extant crocodylians (usually > 3m). Cope’s rule is most frequently conceived as the occurrence of multi-lineage trends of directional evolution towards larger body sizes [7, 8, 11], and this can be evaluated using BM-based models that incorporate a directional trend (parameter μ [80]; see e.g., [33, 62]).

We find little support for trend-like models as a description of crocodylomorph or crocodylian body size evolution. Therefore, we reject the applicability of Cope’s rule to crocodylomorph evolution. This reinforces previous works suggesting that multi-lineage trends of directional body-size evolution are rare over macroevolutionary time scales [33, 67, 160, 161] (but see [19]). Furthermore, our SURFACE model fits indicate that regime shifts towards smaller-bodied descendent regimes occurred approximately as frequently (12–13 times) as shifts to regimes of larger body sizes (10–14 times; Fig. 11), when considering shifts that led to both clades containing multiple and single taxa. Together, these results indicate that long-term increases in the average body size of crocodylomorphs also cannot be explained either by multi-lineage trends of directional evolution towards larger size, or by a biased frequency of transitions to large-bodied descendent regimes.

**Fig. 11.**
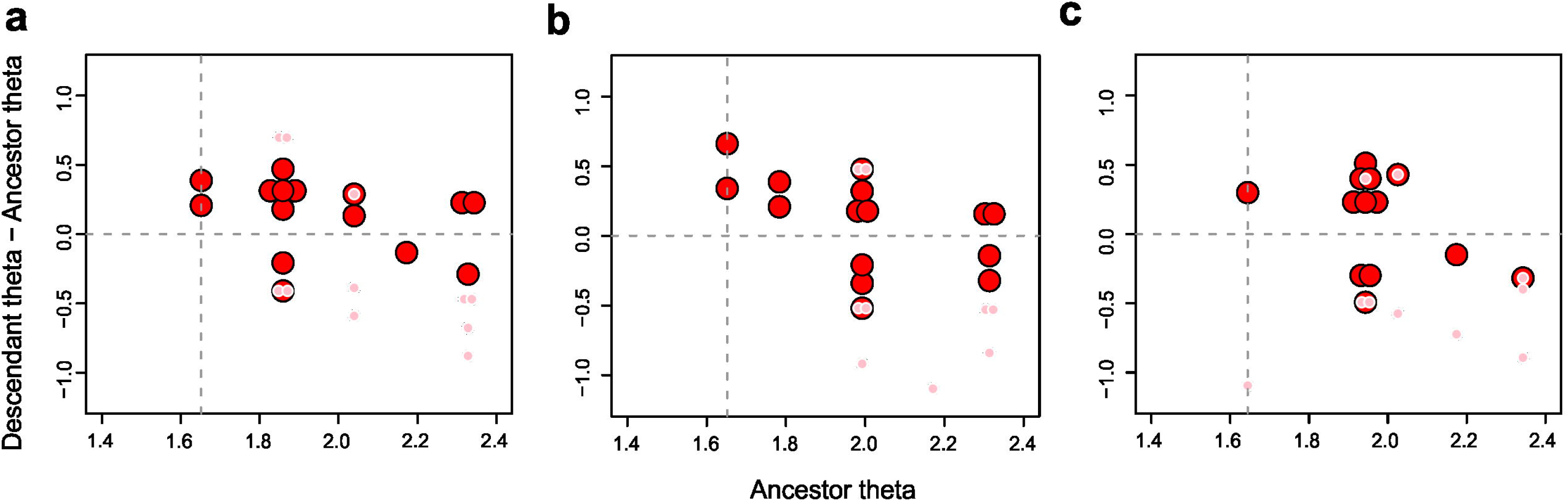
Distribution of regime shifts represented by the difference between descendant and ancestral regimes trait optima values (θ) plotted against the θ of the ancestral regime. Large red circles represent shifts that led to clades containing multiple taxa, while smaller pink circles represent “singleton” regimes, containing only single taxa. Vertical dashed line indicates the ancestral regime for all crocodylomorphs (Z_0_), while horizontal dashed line can be used as a reference to identify regime shifts giving rise to larger (circles above the line) or smaller-bodied (circles below the line) descendants. Circles at the exact same position (i.e., shifts with the same θ values for both ancestral and descendant regimes) were slightly displaced in relation to one another to enable visualization. This plot was constructed using the θ values from trees with different positions of Thalattosuchia: (**a**) Tree number 2, with Thalattosuchia within Neosuchia; Tree number 17, with Thalattosuchia as the sister group of Crocodyliformes; (**c**) Tree number 18, with Thalattosuchia as the sister group of Mesoeucrocodylia. θ values in log_10_ mm, relative to the cranial measurement ODCL (orbito-cranial dorsal length).

Instead, the apparent trend towards larger body sizes can be explained by extinctions among small-bodied regimes. Crocodylomorph body size disparity decreased gradually through the Cretaceous (Fig. 8b). This occurred due to the decreasing abundance of small-bodied species. Despite this, our SURFACE model fits mostly indicate the survival of clades exhibiting small-bodied regimes (θ < 200 cm) until approximately the end of the Mesozoic, (e.g., gobiosuchids, uruguaysuchids, sphagesaurids, hylaeochampsids and some allodaposuchids; Figs. 3, 4, and 5). Many of these small-bodied clades became extinct at least by the Cretaceous/Palaeogene (K/Pg) boundary, resulting in a substantial reduction of small-bodied species. Further reductions among the crown-group (Crocodylia) occurred by the Neogene, from which small-bodied species are absent altogether (Figs. 3, 4, and 5).

This predominance of regimes of large sizes today results from the occurrence of large body sizes in the crown-group, Crocodylia. Our SURFACE analyses focusing on Crocodylia indicate ancestral body size regimes with relatively high values of θ (Z_0_ between 220 and 350 cm). The shift to a larger-sized regime (when compared to smaller-bodied eusuchian regimes) probably occurred at the Late Cretaceous (Figs. 3, 4, and 5), and this same regime was inherited by many members of the clade (predominantly semi-aquatic species). During the Palaeogene, however, shifts to regimes of smaller sizes also occurred (such as in *Tsoabichi greenriverensis*, *Diplocynodon deponiae* and planocraniids), increasing total body size disparity (Fig. 8b). The crocodylian body size distribution shifted upwards mainly during the latter part of the Cenozoic (from the Miocene; Fig. 8b), when even larger-bodied animals occurred (e.g., *Purussaurus* and *Mourasuchus*), combined with the disappearance of lineages of smallest species.

### Correlation of crocodylian body size with global cooling

Our time series regressions demonstrate a moderate to strong correlation between crocodylian size and palaeotemperature (from the Late Cretaceous until the Recent; Table 2). This results from the upward-shift of the crocodylian body size distribution, coinciding with cooling global climates in the second half of the Cenozoic [131, 162]. Even though this is an apparently counter-intuitive relationship, we do not interpret it as a result of direct causation. Previous studies have shown that crocodylian species richness decreased with declining global temperatures of the Cenozoic [36, 37].

Furthermore, the palaeolatitudinal ranges of both marine and continental crocodylomorphs have contracted as temperatures decreased (Fig. 7b; see also [36, 37]). Therefore, the temperatures experienced by evolving lineages of crocodylians are not equivalent to global average temperatures. We propose that the association between global cooling and increasing crocodylian body size results from a systematic reduction of available habits/niches (due to a more restricted geographical distribution), with differential extinction of smaller-bodied species. The hypothesis of selective extinction is also consistent with the decreasing in crocodylian body size disparity during the Cenozoic (Fig. 8b).

### Body size selectivity and diversification across Mesozoic boundaries

Numerous comparative studies have investigated a possible link between extinction risk and animal body size (e.g., [163, 164, 165, 166, 167]). For example, larger body sizes, in association with dietary specializations, might increase susceptibility to extinction in some animal groups, such as hypercarnivorous canids [168, 169]. On the other hand, the recovery of some animal clades after extinction events can also be associated with a subsequent increase in diversity and morphological disparity (e.g., Palaeogene mammals [14]), potentially leading to the exploration of new regions of body size space (i.e., invasions of new body size regimes). Thus, although for some groups (and for some extinctions) body size might play an important role, this is evidently not a generalised pattern across all animals.

For crocodylomorphs, little is known about possible influence of body size on differential extinction. In one of the few studies to quantitatively investigate this, Turner & Nesbitt [45], using femoral length as a proxy for total body size, recognized a drop in mean body size of crocodylomorphs across the Triassic-Jurassic (T–J) boundary. Our SURFACE results, however, indicate otherwise, as all Triassic crocodylomorphs are within a macroevolutionary regime of smaller sizes (θ < 100 cm) when Thalattosuchia is placed within Neosuchia (Fig. 3 and 4). In the other two phylogenetic scenarios, the origin of thalattosuchians (which are predominantly large-bodied animals) is placed either at the middle of the Late Triassic or closer to the T–J boundary (Fig. 5). However, as the first records of thalattosuchians only occur in the Early Jurassic, mean body size increases immediately after the boundary (Fig. 8a). The differences between our results and those found by Turner & Nesbitt [45] might be related to the distinct body size proxies used or to the different taxon sample used, as those authors also included non-crocodylomorph pseudosuchians in their analysis. In this context, we acknowledge that the inclusion in our analyses of larger non-crocodyliform crocodylomorphs, such as *Carnufex carolinensis* (∼ 3 metres [159]), might change our results. Thus, at the moment we do not have empirical or statistical evidence to demonstrate selectivity of body sizes in crocodylomorphs during the end-Triassic extinction.

The Early Jurassic was characterized by key events of crocodylomorph diversification [35] and an increase in morphological disparity [42], following the end-Triassic extinction. Similarly, our body size data suggests an increase in body size disparity after the T–J boundary (Fig. 8b). Although a decrease in disparity is observed subsequently, this is probably due to the relatively few crocodylomorphs known for the latest Early Jurassic and the Middle Jurassic (Sinemurian–Aalenian [36]). Subsequently, the diversification of thalattosuchians during the Late Jurassic, together with the occurrence of smaller- to intermediate-bodied neosuchians (such as atoposaurids and goniopholidids), created the greatest observed disparity of crocodylomorph body sizes during their evolutionary history (Fig. 8b).

Recent studies [170, 171, 172] suggested that a combination of environmental perturbations occurred during the Jurassic-Cretaceous (J/K) transition, which might have led to the extinction of some tetrapod lineages. For crocodylomorphs the boundary is characterised by a decrease in marine diversity [36, 171, 172], highlighted by declines in thalattosuchian diversity, especially among teleosaurids, which suffered widespread extinction (except, apparently, at lower palaeolatitudes [173]). Nevertheless, Wilberg [43] did not find evidence for a substantial decrease in crocodylomorph cranial disparity across the J/K boundary. Similarly, our SURFACE results do not suggest dramatic changes in body size space exploration immediately before or after the J/K boundary (Figs. 3, 4, and 5), and there seems to be no defined body size selectivity across this boundary, as the multiple survivor crocodylomorph lineages were within regimes of very disparate optima values. Furthermore, the decrease in disparity observed in the middle of the Early Cretaceous (i.e., Valanginian–Barremian) is likely due to poor sampling [174], resulting in the scarcity of more completely preserved crocodylomorphs during these stages.

The Late Cretaceous is marked by a remarkable fossil richness of notosuchians, in Gondwana [175, 176], and the diversification of eusuchian crocodylians [177]. Notosuchia exhibits a wide range of body sizes (Fig. 10a), to some extent reflecting its remarkable diversity [36, 176] and morphological disparity [43, 44]. Our model-fitting analyses using only notosuchian data suggest more relaxed modes of body size evolution in Notosuchia (Fig. 6), which is consistent with their high species richness and morphological disparity. This could be explained by a combination of intrinsic (i.e., innovations and/or adaptations, such as a highly modified feeding apparatus [178, 179]) and extrinsic factors (i.e., specific environmental conditions, such as the predominantly hot and arid climate of the Gondwanan landmasses occupied by notosuchians [36, 175]).

Even though our body size data show no specific pattern at the K/Pg boundary, a decline in body size disparity is present through the Late Cretaceous, combined with an increase in mean body size (Fig. 8), a pattern that generally continued through the Cenozoic (although with some short-term fluctuations). This supports the hypothesis that the K/Pg extinction had only minor impacts on crocodylomorphs [35, 36, 37, 43, 181]. Although subsampled estimates of genus richness suggest a decline in terrestrial crocodylomorph diversity during the Late Cretaceous, this occurred prior to the K/Pg boundary, between the Campanian into the Maastrichtian, in both Europe and North America [36]. Indeed, several crocodylomorph subclades lost several species prior to the end of the Cretaceous (in particular notosuchians and non-crocodylian eusuchians [35, 36]; Figs. 3, 4, and 5), and multiple lineages within other groups, such as dyrosaurid tethysuchians and crocodylians, crossed the boundary with little change [37, 180, 181] (Figs. 3, 4, and 5). Our data suggest a long-term pattern of selective extinctions of small-bodied crocodylomorphs, starting from the Late Cretaceous and continuing to the Recent. This may have resulted from a longstanding trend of global cooling [131, 162], resulting in more restricted geographical distributions, and reducing niche availability for crocodylomorphs. This is consistent with our SURFACE results (Figs. 3, 4, and 5), that show very few smaller-bodied regimes (θ < 150 cm) during the Palaeogene and a complete absence after the Neogene. This pattern strikingly contrasts with that proposed for mammals, which may have experienced selectivity against larger bodied taxa across the K/Pg boundary [182], although an increase in body size occurred observed subsequently, during the Palaeogene [14, 15]. The pattern of survival in crocodylomorphs also differs from that suggested for squamates (lizards and snakes), in which small-bodied taxa show evidence of preferential survival [183].

## Conclusions

After an early increase (with the highest peak in the Late Jurassic), crocodylomorph body size disparity experienced sustained decline during virtually its entire evolutionary history. This disparity decrease is combined with an increase of average body size through time, with highest peaks in the Middle Jurassic and today. In particular, the increase in mean body size seen during the Cenozoic (mostly related to crocodylians) co-occurs with an overall decrease in global temperatures.

To further characterise these patterns, we used comparative model-fitting analyses for assessing crocodylomorph body size evolution. Our results show extremely strong support for a multi-peak Ornstein-Uhlenbeck model (SURFACE), rejecting the hypothesis of evolution based on Brownian motion dynamics (including those representing the concept of Cope’s rule). This suggests that crocodylomorph body size evolution can be described within the concept of a macroevolutionary adaptive landscape, with a significant amount of crocodylomorph body size variance evolving from pulses of body size changes, represented by shifts between macroevolutionary regimes (similar to adaptive zones or “maximum adaptive zones” of Stanley [11]). This is reflected in the regime shifts frequently detected at the base of well-recognised and diverse crocodylomorph subclades such as Notosuchia, Thalattosuchia, and Crocodylia.

We did not find strong correlations between our body size data and abiotic factors, indicating that shifts between macroevolutionary regimes are more important for determining large-scale patterns of crocodylomorph body size than isolated climatic factors. However, at more refined temporal and phylogenetic scales, body size variation may track changes in climate. In the case of Crocodylia, a global cooling event might explain the long-term increases in body size, as a result of systematic reduction of available habits/niches (due to a more latitudinally-restricted geographical distribution during cooler global climates), with preferential extinction of smaller-bodied species.

Shifts towards larger sizes are often associated with aquatic/marine or semi-aquatic subclades, indicating that ecological diversification may also be relevant, and suggesting a possible link between aquatic adaptations and larger body sizes in crocodylomorphs. These shifts to larger sizes, occurred throughout crocodylomorph evolutionary history, combined with the extinction of smaller-sized regimes, particularly during the Late Cretaceous and Cenozoic, explain the overall increase in mean body size, as well as the large-bodied distribution of extant crocodylians (all of which are aquatic or semi-aquatic) compared to smaller-bodied early taxa.

## Abbreviations

**Cranial measurements**
DCL: dorsal cranial length
ODCL: orbito-cranial length

**Evolutionary models**
BM: Brownian motion
EB: Early burst
OU: Ornstein-Uhlenbeck
BMS: multi-regime BM model that allows parameter σ2 to vary
OUMV: multi-regime OU model that allows θ and σ2 to vary
OUMA: multi-regime OU model in which θ and α can vary
OUMVA: OU model in which all three parameters (θ, α and σ2) can vary

**Model parameters**
θ: trait optimum of OU-based models
α: attraction parameter of OU-based models
σ^2^: Brownian variance or rate parameter of BM or OU-based models
μ: evolutionary trend parameter of BM-based models
Z_0_: estimated trait value at the root of the tree of OU-based models

**Optimality criteria**
AIC: Akaike’s information criterion
AICc: Akaike’s information criterion for finite sample sizes
BIC: Bayesian information criterion
pBIC: phylogenetic Bayesian information criterion

## Declarations

## Acknowledgements

Access to fossil collections was possible thanks to Lorna Steel (NHMUK), Eliza Howlett (OUMNH), Matthew Riley (CAMSM), Zoltán Szentesi (MTM), Attila Ősi (MTM), Ronan Allain (MNHN), Rainer Schoch (SMNS), Erin Maxwell (SMNS), Marisa Blume (HLMD), Eberhard Frey (SMNK), Oliver Rauhut (BSPG), Max Langer (LPRP/USP), Sandra Tavares (MPMA), Fabiano Iori (MPMA), Thiago Marinho (CPP), Jaime Powell (PVL), Rodrigo Gonzáles (PVL), Martín Ezcurra (MACN), Stella Alvarez (MACN), Alejandro Kramarz (MACN), Patricia Holroyd (UCMP), Kevin Padian (UCMP), William Simpson (FMNH), Akiko Shinya (FMNH), Paul Sereno (UCRC), Tayler Keillor (UCRC), Mark Norell (AMNH),Carl Mehling (AMNH), Judy Galkin (AMNH), Alan Turner (SUNY), Liu Jun (IVPP), Corwin Sullivan (IVPP), Zheng Fang (IVPP), Anna K. Behrensmeyer (USNM), and Amanda Millhouse (USNM). Felipe Montefeltro, Andrew Jones and Giovanne Cidade also provided photographs of many crocodylomorph specimens.

We are thankful to Gene Hunt, whose R functions and scripts (particularly for fitting multi-trend models and SURFACE with pBIC) greatly benefited this study. We further thank Gemma Benevento, Luke Parry, Dave Bapst, and Alan Turner for assistance with the FBD tip-dating method. We also thank Emma Dunne, Daniel Cashmore, and Andrew Jones for help and discussion at different stages of this project, especially related to the use of R. Thorough reviews by two anonymous reviewers helped improve the manuscript. We thank the editor R. Alexander Pyron for handling the manuscript. Silhouettes of crocodylomorph representatives in figures are from illustrations by Dmitry Bogdanov, Smokeybjb, and Nobumichi Tamura, hosted at Phylopic (http://phylopic.org), where license information is available.

## Availability of data and material

The data generated and/or analysed during the current study, as well as R codes used for macroevolutionary analyses and supplementary results, are included within the article and its additional files.

## Authors’ contributions

PLG, RBJB and RJB designed the study. PLG and MB collected the data. PLG analysed the data. All authors participated in drafting the manuscript. All authors read and approved the final manuscript.

## Consent for publication

Not applicable.

## Ethics approval and consent to participate

Not applicable.

## Supplementary methods

### Proxy for total body length

Equations based on modern species, using either cranial (e.g., Webb & Messel, 1978; Hall & Portier, 1994; Sereno *et al*., 2001, Hurlburt *et al*., 2003; Platt *et al*., 2009; 2011) or postcranial measurements (e.g., Bustard & Singh, 1977; Farlow *et al*., 2005), have predominantly been used for estimating total body size of extinct crocodylomorph species. Although some of these approaches have been claimed to work well when applied to extinct taxa (e.g., Farlow *et al*., 2005), they are expected to be less accurate for extinct species that have different body proportions to those of extant species (e.g., Pol *et al*., 2012; Young *et al*., 2011; 2016; Godoy *et al*., 2016; but see Figure S1). An alternative approach that has been suggested is to use clade-specific equations that are derived from regressions using fossil specimens with complete skeletons preserved, such as the recently proposed equations for estimating body length in the highly specialised marine clade Thalattosuchia (Young *et al*., 2011; 2016). Nevertheless, using this approach for the entire Crocodylomorpha would require numerous different equations and, consequently, complete specimens for all desired subclades.

Furthermore, Campione & Evans (2012) demonstrated a universal scaling relationship between proximal (stylopodial) limb bone circumferences and the body masses of terrestrial tetrapods. For instance, their equations, using both femur and humerus circumference, have been applied to estimate body mass of fossil dinosaurs (e.g., Benson *et al*. 2014; 2018; Carballido *et al.*, 2017). However, due to a historical neglect of crocodylomorph postcranial anatomy, especially for Mesozoic taxa (Godoy *et al*., 2016), relatively less information is available on this part of the skeleton. Based on data collected for the present study, total or partial skull lengths (i.e., complete skulls or lacking only the snouts) can be measured in fossil specimens of approximately 50% of crocodylomorph species, whereas femoral and humeral shaft circumferences or lengths can only be measured in 35% of species. This greatly reduces the number of taxa that can be sampled and limits the utility of using postcranial elements as a proxy for body size. Similar problems exist for other methods, such as the “Orthometric Linear Unit” proposed by Romer & Price (1940) that uses dorsal centrum cross section (Currie, 1978), as well as volumetric reconstructions (e.g., Colbert, 1962; Hurlburt, 1999; Motani, 2001; Bates et al., 2009; Sellers et al., 2012), since relatively complete postcranial specimens are required.

Thus, aiming for a proxy (or proxies) for total body size that could maximised sample size (for a study encompassing the entire evolutionary history of Crocodylomorpha), we decided to use two cranial measurements: total dorsal cranial length (DCL) and dorsal orbito-cranial length (ODCL), which is measured from the anterior margin of the orbit to the posterior margin of the skull. By using actual cranial measurements, rather than estimated total body length, we avoid the addition of possible errors to our model-fitting analyses (Figure S1). Furthermore, the range of body sizes among living and extinct crocodylomorphs is considerably greater than variation among size estimates for single species. Therefore, we expect to recover the most important macroevolutionary body size changes in our analyses even when using only cranial measurements.

**Figure S1.**
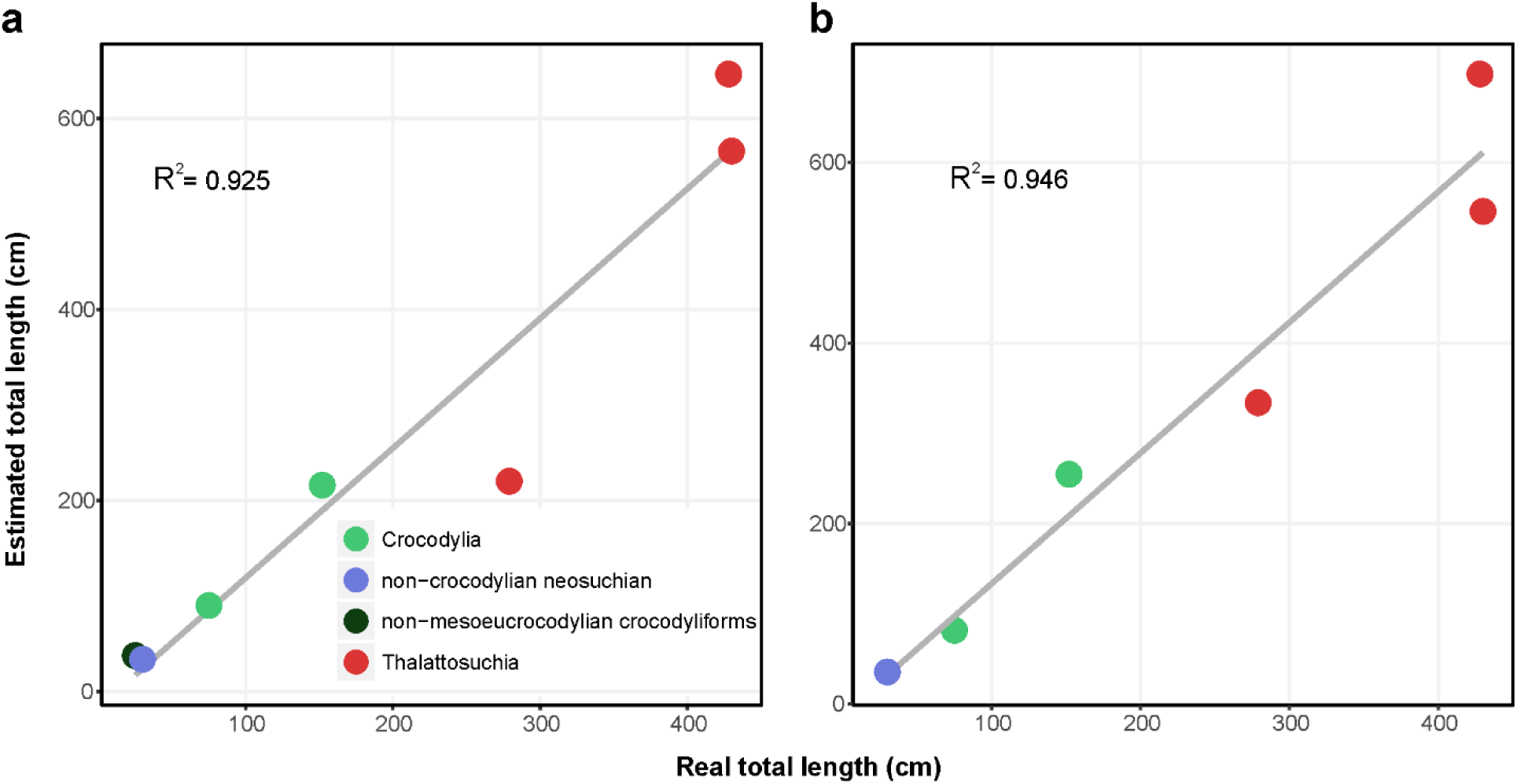
Expected error for total body length estimated from cranial measurements. Real total body length, measured from some complete fossil crocodylomorph specimens, is plotted against total length estimated from the cranial measurements DCL (**a**) and ODCL (**b**) (equations from Hurlburt *et al*., 2003), exemplifying the amount of error expected when using cranial measurements to estimate total body length of crocodylomorphs. R^2^ value illustrates the strength of the correlation between real and estimated total lengths. Colours represent different mono- or paraphyletic crocodylomorph groups. See Table S1 for information on the specimens used for the construction of these plots.

**Table S1.** List of fossil specimens with complete skeleton preserved which data was used for creating Figure S1 of the manuscript. “DCL” is the total length estimated using the cranial measurement dorsal cranial length, “ODCL” is the total length estimated when using dorsal orbito-cranial length, and “Real TL” is the real total body length measured from the specimen. All measurements in centimetres.

| Species (specimen) | DCL | ODCL | Real TL | Source of information |
| --- | --- | --- | --- | --- |
| *Shantungosuchus chuhsienensis* (IVPP V2484) | 37.66 | N/A | 25 | First-hand observation |
| *Alligatorellus beaumonti* (BSPG 1937 I 26) | 33.77 | 35.72 | 30 | First-hand observation |
| *Diplocynodon ratelii* (MNHN.F SG 13728ab) | 216.15 | 254.74 | 152 | First-hand observation |
| *Diplocynodon darwini* (HLMD-Me 10262) | 90.02 | 81.86 | 75 | First-hand observation |
| *Platysuchus multiscrobiculatus* (SMNS 9930) | 220.22 | 334.08 | 279 | Young *et al.* (2016) |
| *Steneosaurus bollensis* (SMNS 54063) | 565.97 | 545.90 | 430 | Young *et al.* (2016) |
| *Steneosaurus leedsi* (NHMUK R 3806) | 646.58 | 698.01 | 428 | Young *et al.* (2016) |

### Supertree construction and alternative topologies

The supertree used as the phylogenetic framework for the macroevolutionary analyses was constructed using an informal approach. For such, we started with the MRP (matrix representations with parsimony) supertree of Bronzati *et al*. (2015), and then used some recently published phylogenetic hypotheses to create and updated version, by manually modifying the tree using the software Mesquite (Version 3.51; Maddison & Maddison, 2018). For this updated version, we added some taxa, removed others, and also changed the position of a few more, always aiming to include as many species as possible (especially the ones for which we had body size data available), but also to incorporate more well-resolved relationships from recent studies.

The supertree presented by Bronzati *et al*. (2015) is restricted to Crocodyliformes, which is less inclusive than Crocodylomorpha. Thus, we added non-crocodyliform crocodylomorphs taxa following the phylogenetic hypotheses presented by Pol *et al*. (2013) and Leardi *et al*. (2017). Within Crocodyliformes, as in Bronzati *et al*. (2015) and other recent studies (e.g., Andrade *et al.,* 2011; Montefeltro *et al.,* 2013; Pol *et al.,* 2014; Turner & Pritchard, 2015; Buscalioni, 2017), taxa classically associated to “Protosuchia” are paraphyletic arranged in relation to Mesoeucrocodylia, with smaller subgroups displayed following Bronzati *et al*. (2015) (but see below for differences in this region of the tree in the alternative topologies). Accordingly, *Hsisosuchus* is the sister-group of Mesoeucrocodylia (as in Clark, 2011, Pol *et al.,* 2014; Buscalioni, 2017) and the following groups represent taxa successively more distant to Mesoeucrocodylia: Shartegosuchidae (following Clark 2011); an unnamed clade composed by taxa such as *Sichuanosuchus* and *Shantungosuchus;* an unnamed clade composed by *Zaraasuchus* and *Gobiosuchus* (following Pol *et al.,* 2014); Protosuchidae (following Clark 2011; Pol *et al.,* 2014; Turner & Pritchard, 2015).

Within Mesoeucrocodylia, Notosuchia corresponds to the sister group of all the other mesoeucrocodylians (= Neosuchia in our topology), similar to what is presented by Andrade *et al*. (2011), Pol *et al*. (2014), and Turner & Pritchard (2015). Yet, Notosuchia comprises forms such as baurusuchids, sebecosuchians, peirosaurids, sphagesaurids, uruguaysuchids, and *Araripesuchus.* The relationships among taxa within Notosuchia follow the general arrangement presented by Pol *et al*. (2014).

One of the branches at the basal split of Neosuchia leads to a clade composed by longirostrine forms, which includes Thalattosuchia and Tethysuchia (i.e. Dyrosauridae and “pholidosaurids”). Arrangement between these groups (i.e. sister-group relationship between Thalattosuchia and Tethysuchia) follows that recovered in the supertree of Bronzati *et al*. (2015). Within Tethysuchia, “pholidosaurids” are paraphyletic in relation to Dyrosauridae (also found in Pol *et al.,* 2014; Young *et al*., 2017 and Meunier & Larsson, 2017). Relationships among Dyrosauridae follow Hastings *et al*. (2015). Relationships among thalattosuchians follow Young (2014) and Herrera *et al*. (2015).

The sister-group of the longirostrine clade mentioned above contains Eusuchia and its closest relatives such as Atoposauridae and Goniopholididae. The latter is depicted as the sister group of Eusuchia, whereas the former corresponds to the sister group of Eusuchia + Goniopholididae. This arrangement follows that recovered in Pol *et al*. (2014) and Bronzati *et al*. (2015). Regarding the internal relationships of Goniopholididae, we follow the hypotheses of Martin *et al*. (2016) and Ristevski et al. (2018). For Atoposauridae, we follow the arrangements presented by Tennant *et al*. (2016) and Schwarz *et al*. (2017). For Paralligatoridae and Susisuchidae, we followed the phylogenetic hypotheses of Turner (2015) and Turner & Pritchard (2015).

In relation to non-crocodylian eusuchians, we mainly follow the topology of Bronzati *et al*. (2015), with modifications to accommodate the arrangements proposed by Turner (2015) and Turner & Pritchard (2015) within Paralligatoridae and Susisuchidae. Regarding the interrelationships of the crown-group, as well as the position of Hylaeochampsidae + Allodaposuchidae as the sister group of Crocodylia, we follow the topology of Narváez *et al*. (2015). For the relationships within the crown-group, we follow Brochu (2012), Brochu *et al*. (2012), Scheyer *et al*. (2013) and Narváez *et al*. (2015).

Additionally, two alternative topologies were also manually constructed, for testing the impact of alternative positions of Thalattosuchia. The “longirostrine problem”, which mostly concerns the position of Thalattosuchia, has been largely debated in phylogenetic studies of Crocodylomorpha (e.g., Clark, 1994; Pol & Gasparini, 2009; Wilberg, 2015). Because of the possible impact that a group like Thalattosuchia (i.e. of relatively old origin and many species within it) can inflict in our model-fitting analyses, we built two alternative trees to test the effects related to this phylogenetic uncertainty. Apart from the position of Thalattosuchia described above (within Neosuchia), two main alternative scenarios for the position of the group within Crocodylomorpha were proposed (see Wilberg, 2015). The first places Thalattosuchia as the sister group of all other mesoeucrocodylians (= Notosuchia + Neosuchia) (e.g., Larsson & Sues, 2007; Montefeltro *et al.,* 2013), and was depicted in one of our alternative topologies. The other alternative topology places Thalattosuchia as the sister group of Crocodyliformes (following Wilberg, 2015). Only the position of Thalattosuchia has been altered in these alternative topologies. Relationships among other taxa, including the relationship among thalattosuchians, were kept as in the first topology, described above.

### Additional time-scaling methods

For time-calibrating our trees, apart from the Bayesian tip-dating approach, we also used three different *a posteriori* time-scaling (APT) methods: the minimum branch length (*mbl*), the *cal3* and the *extended Hedman* methods. These methods were used only for the initial model comparison.

For these methods, ages (first and last occurrence dates) were initially obtained from the Paleobiology Database, but were then checked using primary sources in the literature (see Additional file 6 for ages of all taxa). To accommodate uncertainties related to the ages of terminal taxa (i.e., most taxon ages are based on single occurrences, known only within rather imprecise bounds), we treated these first and last occurrences dates as maximum and minimum possible ages and drew terminal dates for time-calibration from a uniform distribution between these.

First, the *mbl* method (Laurin, 2004), which requires a minimum branch duration to be set *a priori*, to avoid the presence of undesirable and unrealistic zero-length branches (Bapst, 2014*a*, *b*). For our analyses, the minimum of 1 Myr was set.

Second, the *cal3* method, which is a stochastic calibration method that requires estimates of sampling and diversification (branching and extinction) rates to draw likely divergence dates under a birth–death-sampling model (Bapst, 2014; Lloyd *et al*., 2016). The fact that most crocodylomorph taxa are singletons (i.e., very few genera or species have multiple occurrences in different time intervals) prevented us from directly calculating speciation, extinction and sampling rates needed as inputs to the *cal3* method. Thus, when using this time-scaling method for our analyses, we adopted the same rates estimated for dinosaurs in Lloyd *et al*. (2016) (i.e., extinction and speciation rates = 0.935; sampling rate = 0.018), which used the apparent range-frequency distribution of dinosaurs in the Paleobiology Database for these estimates. Although essentially different from that of dinosaurs, the crocodylomorph fossil record is arguably comparable enough to result in similar rates (i.e., in both groups, many species are based on only single occurrences, having therefore no meaningful range data; Benson *et al*., 2018), and *a posteriori* comparison to other time-scaling methods demonstrated that results were qualitatively reasonable.

Finally, the *extended Hedman* method was proposed by Lloyd *et al*. (2016), and is expansion of the approach presented by Hedman (2010). It is a probabilistic that uses the ages of successive outgroup taxa relative to the age of the node of interest to date this node by sampling from uniform distributions (Lloyd *et al*. 2016, Brocklehurst, 2017).

Since the input phylogenies (i.e., the three alternatives topologies of the supertree, see above) were not completely resolved, we randomly resolved the polytomies, generating 20 completely resolved trees (the same number of trees was time-scaled with the FBD method) for each alternative phylogenetic scenario (i.e., with different positions of Thalattosuchia). These trees were then time-scaled using the three time-calibration methods. Time-scaling with the *mbl* and *cal3* methods were performed using the package *paleotree* (Bapst, 2012) in R version 3.5.1 (R Core Team, 2018), whilst the *Hedman* method was implemented also in R, using the protocol published by Lloyd *et al*. (2016).

Time bins used for time series correlations and disparity calculation

| LOWER LIMIT (IN MYR) | UPPER LIMIT (IN MYR) |
| --- | --- |
| 7.246 | 0 |
| 15.97 | 7.246 |
| 23.03 | 15.97 |
| 33.9 | 23.03 |
| 40.4 | 33.9 |
| 48.6 | 40.4 |
| 55.8 | 48.6 |
| 61.7 | 55.8 |
| 66.043 | 61.7 |
| 70.6 | 66.043 |
| 84.9 | 70.6 |
| 94.3 | 84.9 |
| 99.7 | 94.3 |
| 112.6 | 99.7 |
| 125.45 | 112.6 |
| 136.4 | 125.45 |
| 145.5 | 136.4 |
| 155.7 | 145.5 |
| 164.7 | 155.7 |
| 171.6 | 164.7 |
| 183 | 171.6 |
| 189.6 | 183 |
| 201.6 | 189.6 |
| 205.6 | 201.6 |
| 221.5 | 205.6 |
| 235 | 221.5 |
| 242 | 235 |
| 252.3 | 242 |

### FBD consensus trees

**Figure S2.**
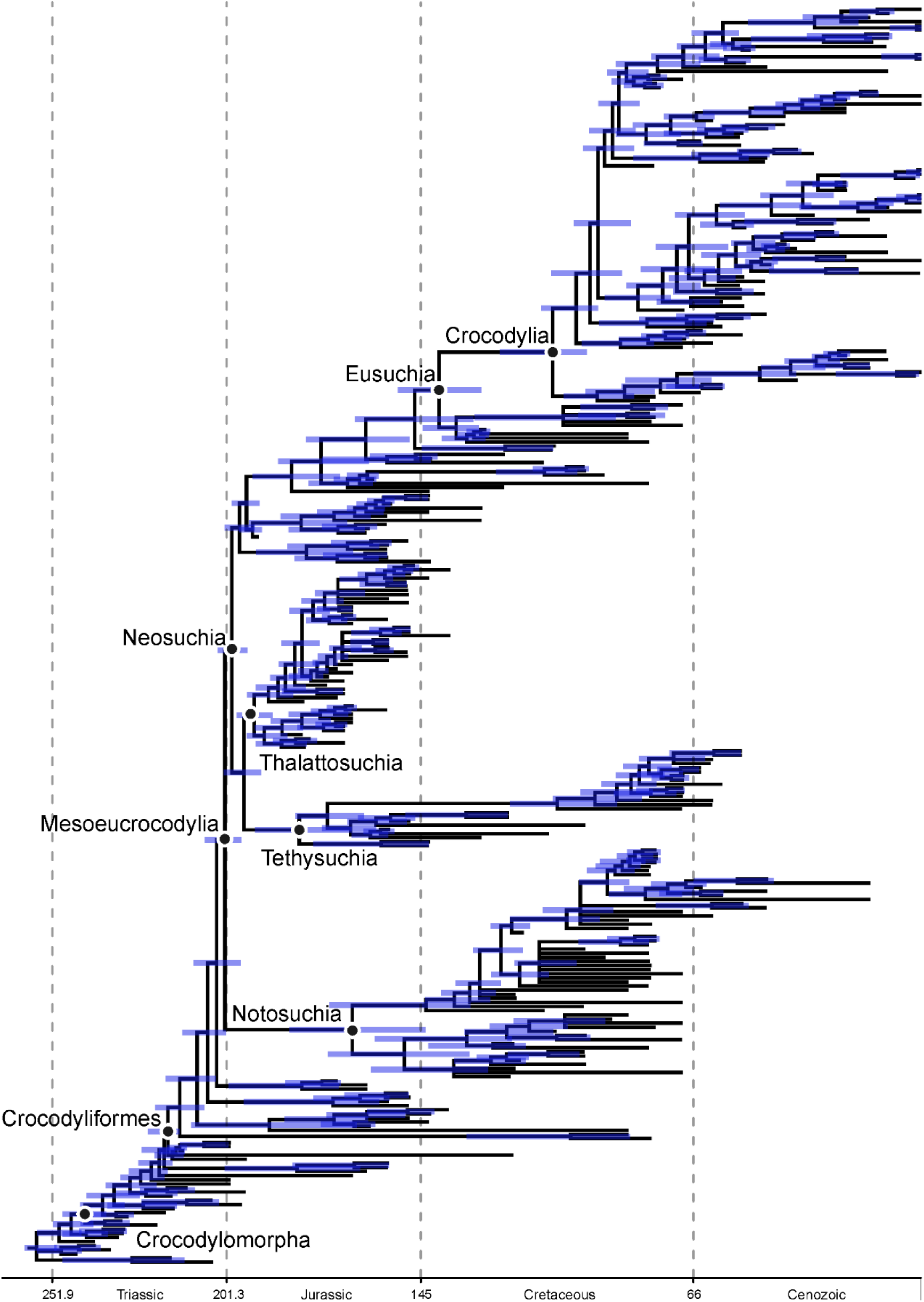
Consensus tree (50% majority rule tree) of Crocodylomorpha, with Thalattosuchia within Neosuchia. Node ages were inferred under a fossilized birth-death process, performing 10,000,000 generations of MCMC analyses. Blue bars indicate 95% Highest Posterior Density (HPD) time intervals.

**Figure S3.**
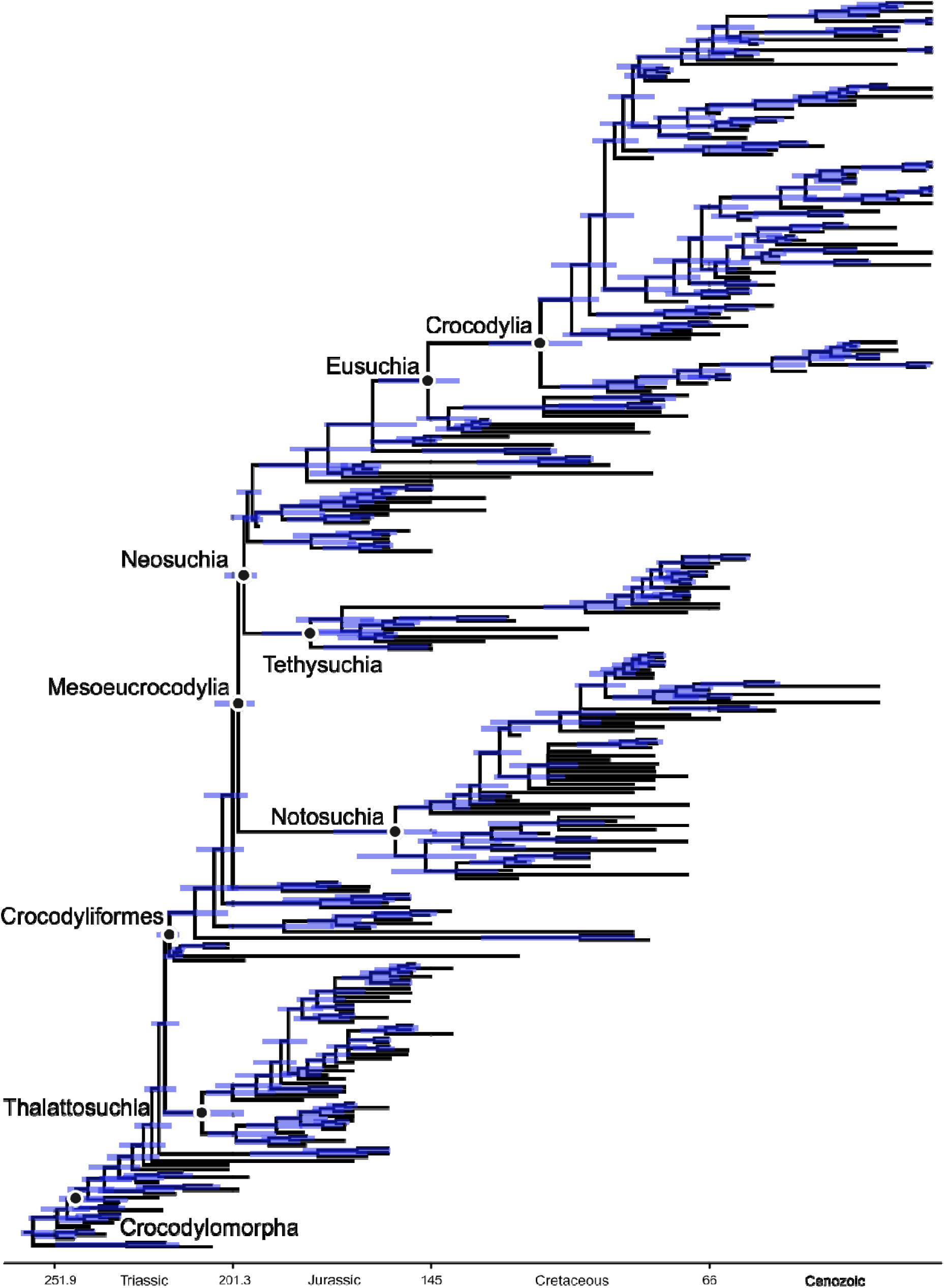
Consensus tree (50% majority rule tree) of Crocodylomorpha, with Thalattosuchia as the sister group of Crocodyliformes. Node ages were inferred under a fossilized birth-death process, performing 10,000,000 generations of MCMC analyses. Blue bars indicate 95% Highest Posterior Density (HPD) time intervals.

**Figure S4.**
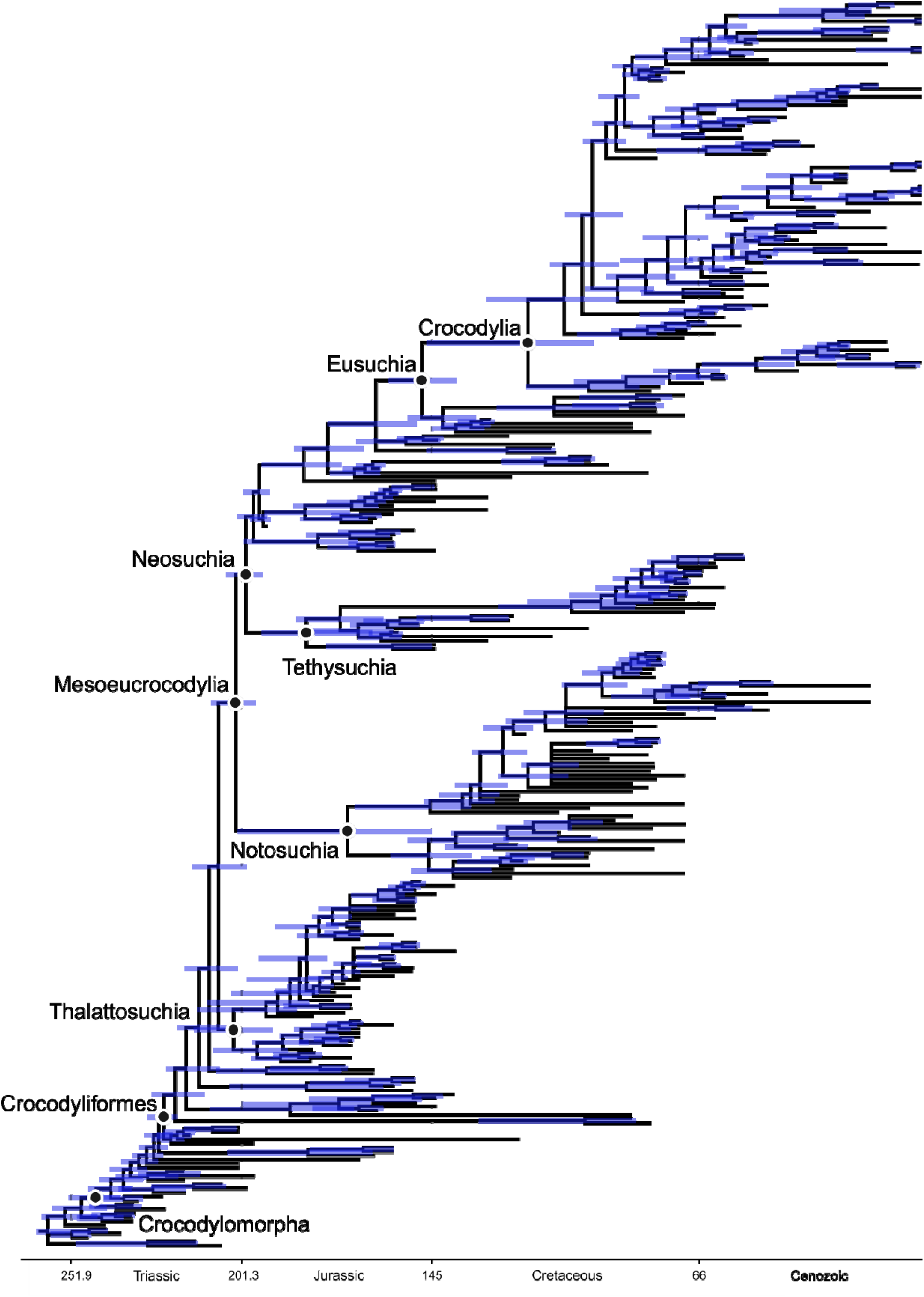
Consensus tree (50% majority rule tree) of Crocodylomorpha, with Thalattosuchia as the sister group of Mesoeucrocodylia. Node ages were inferred under a fossilized birth-death process, performing 10,000,000 generations of MCMC analyses. Blue bars indicate 95% Highest Posterior Density (HPD) time intervals.

### Initial model comparison using APT time-scaling methods

**Figure S5.**
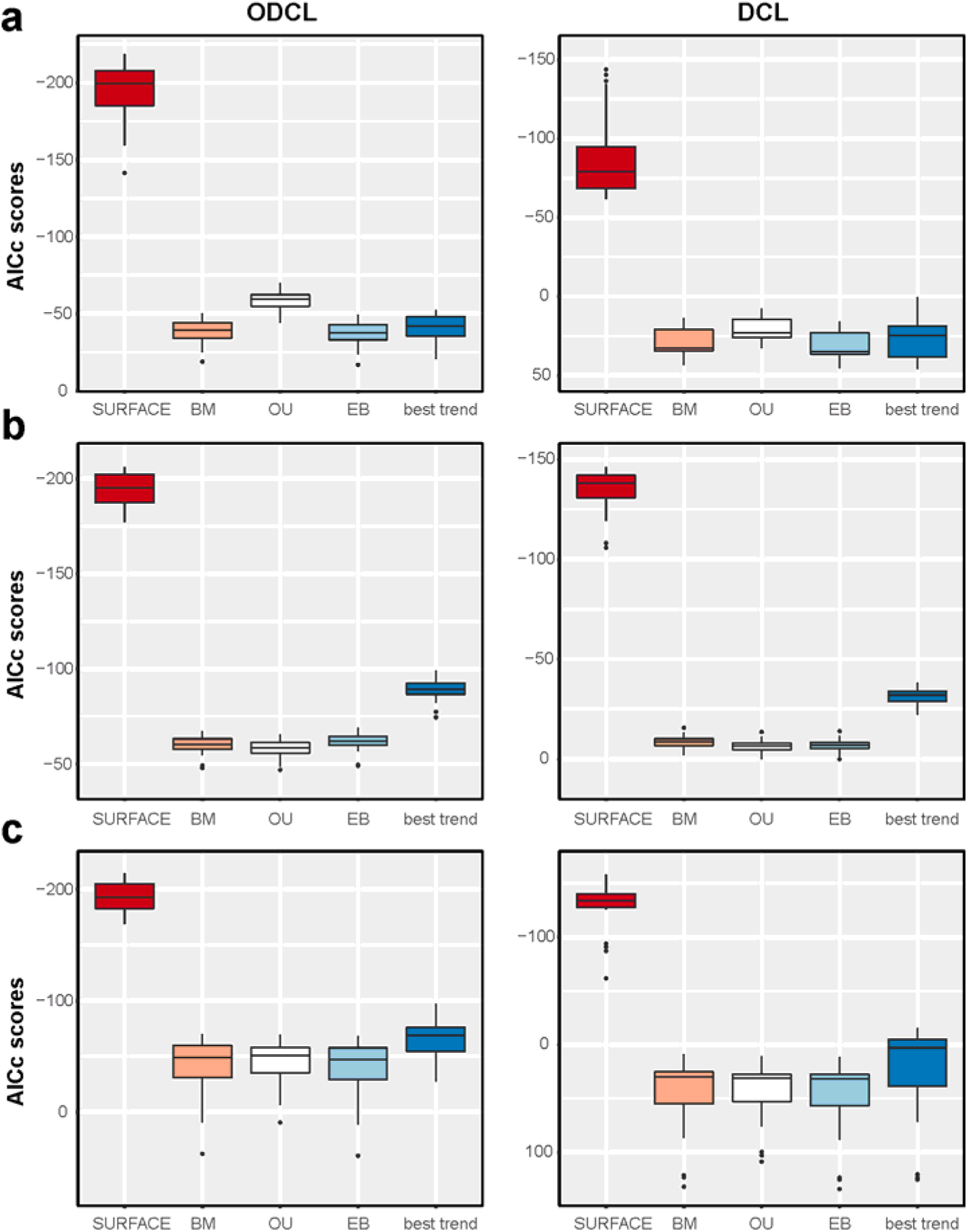
AICc scores of the evolutionary models fitted to crocodylomorph phylogeny and body size data. Results shown for two cranial measurements datasets (ODCL in the left column and DCL in the right one), as well as using three different APT time-scaling (i.e., *a posteriori*) methods to time-calibrate 20 randomly resolved phylogenies of Crocodylomorpha: (a) mbl, (b) Hedman, and (c) cal3 methods. For the trend-like models, only the AICc of the best model (“best trend”) is shown.

### Correlations with abiotic factors

Most of the regression analyses with body size and palaeotemperature data (Tables S3– S16) revealed very weak or non-significant correlations. In some cases, we did find significant correlations, but they were frequently inconsistent (i.e., correlations did not persist in both ODCL and DCL datasets or were absent when accounting for serial autocorrelation using GLS). The only conspicuous exception was found between mean body size values and palaeotemperatures from the Late Cretaceous (Maastrichtian) to the Recent (and, in particular, when using only taxa of the crown-group Crocodylia [Tables S8 and S9]).

A similar scenario was found for the correlation test between body size and paleolatitude (Tables S17–S30), with very weak or non-significant correlations. Our phylogenetic regressions of found some significant correlations, but in all cases the coefficient of determination (R^2^) was very low (always smaller than 0.1), indicating that the correlation is very weak and only a small proportion (less than 10%) of the body size variation observed can be explained by the palaeolatitudinal data.

### Palaeotemperature

**Table S3.**
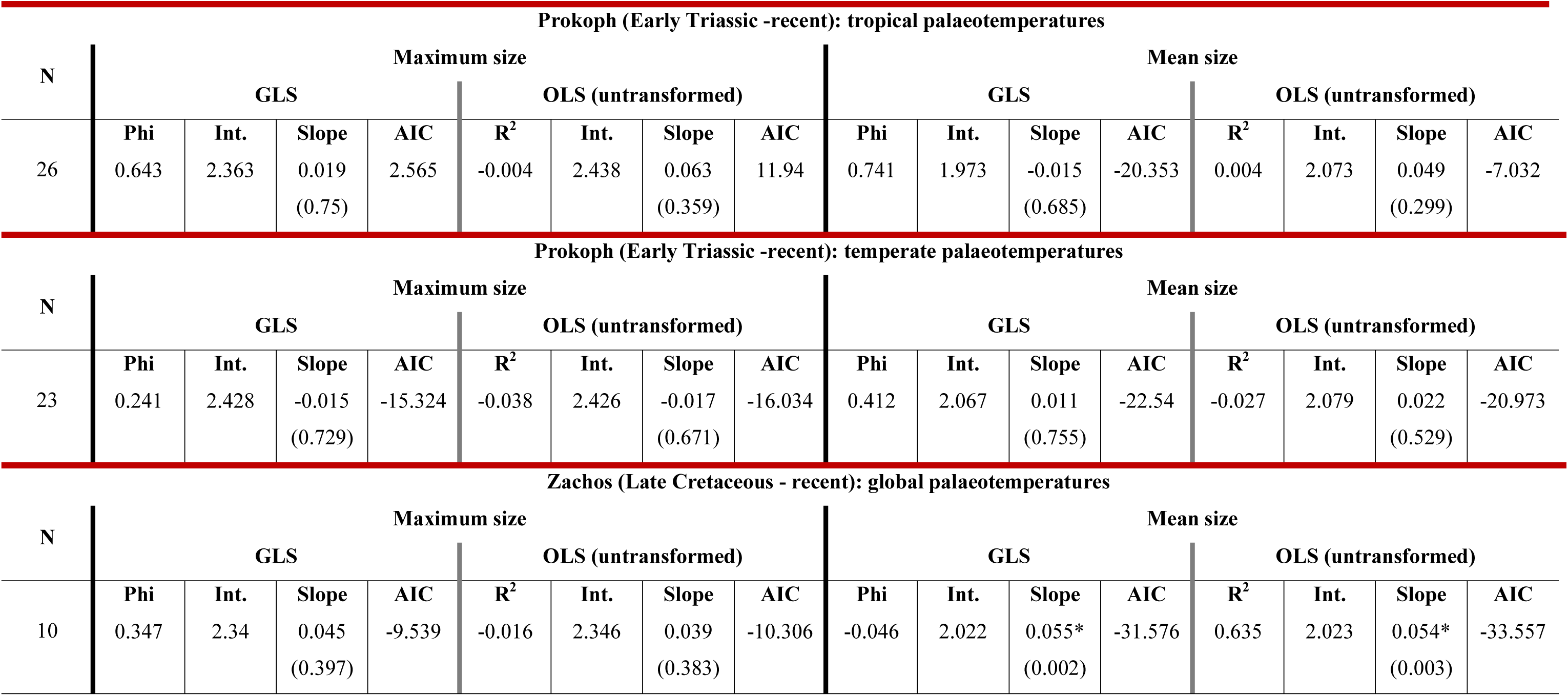
Results of regressions of body size proxy (maximum and mean log-transformed ODCL, using all species in the dataset) on the palaeotemperature proxies (δ^18^O data for tropical and temperate regions from Prokoph *et al*. (2008), and global δ^18^O data from Zachos *et al*. (2008)). Possible correlation was analysed using generalised least squares (GLS) regressions, incorporating a first-order autoregressive model, as well as ordinary least squares (OLS) regressions using untransformed data (assuming no serial correlation). *Significant at alpha = 0.05.

| Prokoph (Early Triassic -recent): tropical palaeotemperatures |  |  |  |  |  |  |  |  |  |  |  |  |  |  |  |  |
| --- | --- | --- | --- | --- | --- | --- | --- | --- | --- | --- | --- | --- | --- | --- | --- | --- |
| N | Maximum size |  |  |  |  |  |  |  | Mean size |  |  |  |  |  |  |  |
|  | GLS |  |  |  | OLS (untransformed) |  |  |  | GLS |  |  |  | OLS (untransformed) |  |  |  |
|  | Phi | Int. | Slope | AIC | R<sup>2</sup> | Int. | Slope | AIC | Phi | Int. | Slope | AIC | R<sup>2</sup> | Int. | Slope | AIC |
| 26 | 0.643 | 2.363 | 0.019<br>(0.75) | 2.565 | -0.004 | 2.438 | 0.063<br>(0.359) | 11.94 | 0.741 | 1.973 | -0.015<br>(0.685) | -20.353 | 0.004 | 2.073 | 0.049<br>(0.299) | -7.032 |
| Prokoph (Early Triassic -recent): temperate palaeotemperatures |  |  |  |  |  |  |  |  |  |  |  |  |  |  |  |  |
| N | Maximum size |  |  |  |  |  |  |  | Mean size |  |  |  |  |  |  |  |
|  | GLS |  |  |  | OLS (untransformed) |  |  |  | GLS |  |  |  | OLS (untransformed) |  |  |  |
|  | Phi | Int. | Slope | AIC | R<sup>2</sup> | Int. | Slope | AIC | Phi | Int. | Slope | AIC | R<sup>2</sup> | Int. | Slope | AIC |
| 23 | 0.241 | 2.428 | -0.015<br>(0.729) | -15.324 | -0.038 | 2.426 | -0.017<br>(0.671) | -16.034 | 0.412 | 2.067 | 0.011<br>(0.755) | -22.54 | -0.027 | 2.079 | 0.022<br>(0.529) | -20.973 |
| Zachos (Late Cretaceous - recent): global palaeotemperatures |  |  |  |  |  |  |  |  |  |  |  |  |  |  |  |  |
| N | Maximum size |  |  |  |  |  |  |  | Mean size |  |  |  |  |  |  |  |
|  | GLS |  |  |  | OLS (untransformed) |  |  |  | GLS |  |  |  | OLS (untransformed) |  |  |  |
|  | Phi | Int. | Slope | AIC | R<sup>2</sup> | Int. | Slope | AIC | Phi | Int. | Slope | AIC | R<sup>2</sup> | Int. | Slope | AIC |
| 10 | 0.347 | 2.34 | 0.045<br>(0.397) | -9.539 | -0.016 | 2.346 | 0.039<br>(0.383) | -10.306 | -0.046 | 2.022 | 0.055*<br>(0.002) | -31.576 | 0.635 | 2.023 | 0.054*<br>(0.003) | -33.557 |

**Table S4.**
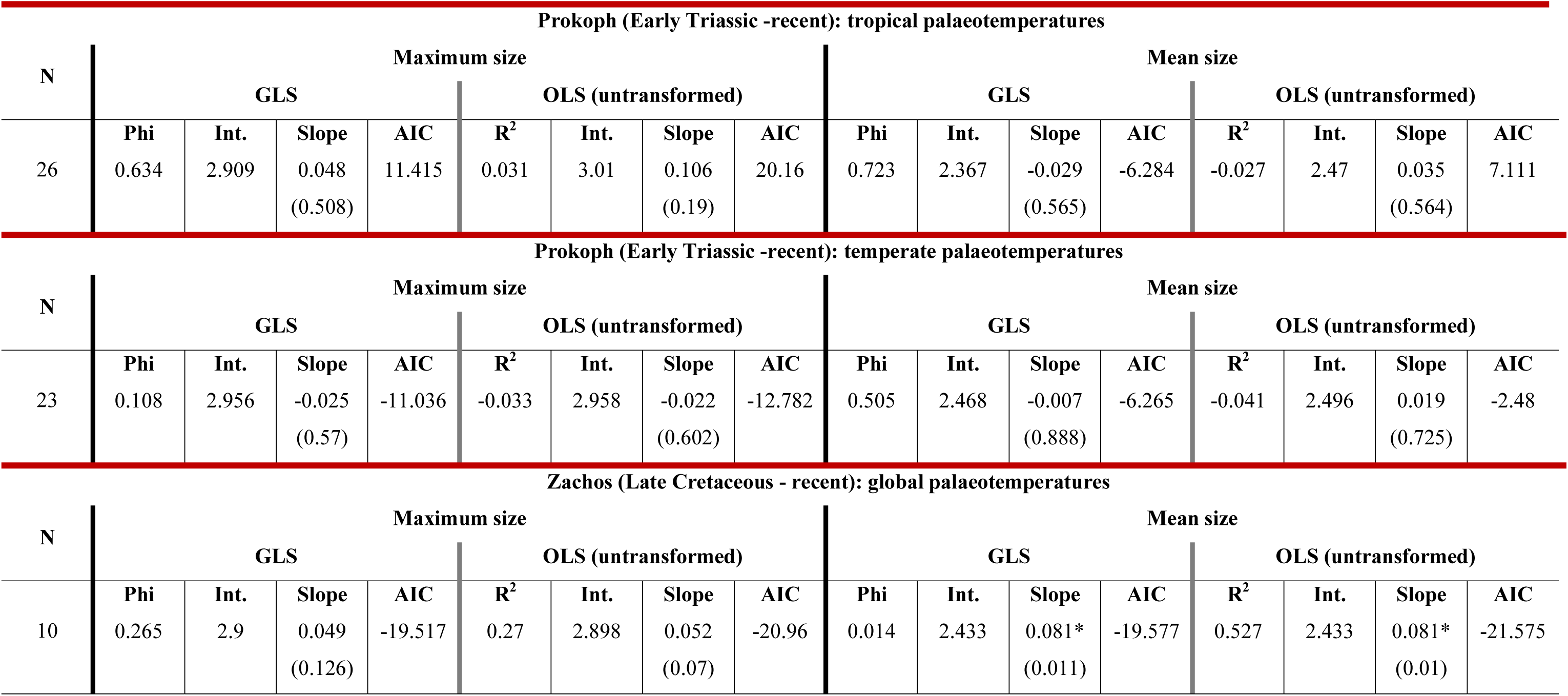
Results of regressions of body size proxy (maximum and mean log-transformed DCL, using all species in the dataset) on the palaeotemperature proxies (δ^18^O data for tropical and temperate regions from Prokoph *et al*. (2008), and global δ^18^O data from Zachos *et al*. (2008)). Possible correlation was analysed using generalised least squares (GLS) regressions, incorporating a first-order autoregressive model, as well as ordinary least squares (OLS) regressions using untransformed data (assuming no serial correlation). *Significant at alpha = 0.05.

**Table S5.** Results of regressions of body size proxy (maximum and mean log-transformed ODCL, using only marine species in the dataset) on the palaeotemperature proxies (δ^18^O data for tropical and temperate regions from Prokoph *et al*. (2008), and global δ^18^O data from Zachos *et al*. (2008)). Possible correlation was analysed using generalised least squares (GLS) regressions, incorporating a first-order autoregressive model, as well as ordinary least squares (OLS) regressions using untransformed data (assuming no serial correlation). *Significant at alpha = 0.05.

| Prokoph (Early Triassic -recent): tropical palaeotemperatures |  |  |  |  |  |  |  |  |  |  |  |  |  |  |  |  |
| --- | --- | --- | --- | --- | --- | --- | --- | --- | --- | --- | --- | --- | --- | --- | --- | --- |
| N | Maximum size |  |  |  |  |  |  |  | Mean size |  |  |  |  |  |  |  |
|  | GLS |  |  |  | OLS (untransformed) |  |  |  | GLS |  |  |  | OLS (untransformed) |  |  |  |
|  | Phi | Int. | Slope | AIC | R<sup>2</sup> | Int. | Slope | AIC | Phi | Int. | Slope | AIC | R<sup>2</sup> | Int. | Slope | AIC |
| 18 | 0.56 | 2.358 | -0.025<br>(0.536) | -25.542 | 0.201 | 2.276 | -0.11*<br>(0.035) | -24.167 | 0.014 | 2.239 | -0.017<br>(0.451) | -49.171 | -0.023 | 2.239 | -0.017<br>(0.448) | -51.167 |
| Prokoph (Early Triassic -recent): temperate palaeotemperatures |  |  |  |  |  |  |  |  |  |  |  |  |  |  |  |  |
| N | Maximum size |  |  |  |  |  |  |  | Mean size |  |  |  |  |  |  |  |
|  | GLS |  |  |  | OLS (untransformed) |  |  |  | GLS |  |  |  | OLS (untransformed) |  |  |  |
|  | Phi | Int. | Slope | AIC | R<sup>2</sup> | Int. | Slope | AIC | Phi | Int. | Slope | AIC | R<sup>2</sup> | Int. | Slope | AIC |
| 17 | 0.708 | 2.423 | 0.079<br>(0.059) | -23.953 | -0.066 | 2.398 | -0.002<br>(0.955) | -16.916 | 0.758 | 2.273 | 0.022<br>(0.058) | -66.294 | -0.027 | 2.273 | 0.011<br>(0.463) | -56.901 |
| Zachos (Late Cretaceous - recent): global palaeotemperatures |  |  |  |  |  |  |  |  |  |  |  |  |  |  |  |  |
| N | Maximum size |  |  |  |  |  |  |  | Mean size |  |  |  |  |  |  |  |
|  | GLS |  |  |  | OLS (untransformed) |  |  |  | GLS |  |  |  | OLS (untransformed) |  |  |  |
|  | Phi | Int. | Slope | AIC | R<sup>2</sup> | Int. | Slope | AIC | Phi | Int. | Slope | AIC | R<sup>2</sup> | Int. | Slope | AIC |
| 10 | -0.143 | 2.422 | -0.045<br>(0.054) | -21.432 | 0.22 | 2.417 | -0.042<br>(0.096) | -23.261 | 0.627 | 2.252 | -0.006<br>(0.654) | -39.327 | -0.088 | 2.241 | 0.005<br>(0.617) | -38.084 |

**Table S6.**
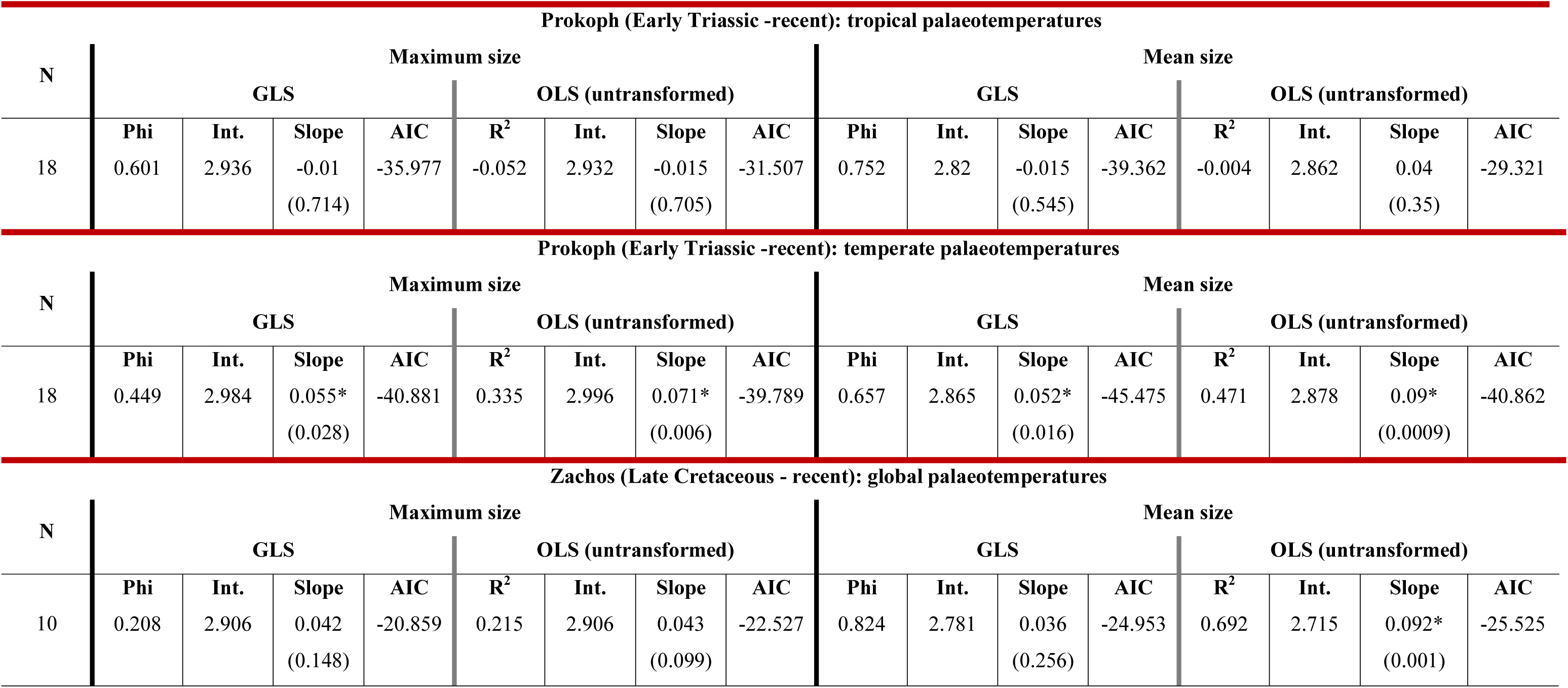
. Results of regressions of body size proxy (maximum and mean log-transformed DCL, using only marine species in the dataset) on the palaeotemperature proxies (δ^18^O data for tropical and temperate regions from Prokoph *et al*. (2008), and global δ^18^O data from Zachos *et al*. (2008)). Possible correlation was analysed using generalised least squares (GLS) regressions, incorporating a first-order autoregressive model, as well as ordinary least squares (OLS) regressions using untransformed data (assuming no serial correlation). *Significant at alpha = 0.05.

**Table S7.** Results of regressions of body size proxy (maximum and mean log-transformed ODCL, using only non-marine species in the dataset) on the palaeotemperature proxies (δ^18^O data for tropical and temperate regions from Prokoph *et al*. (2008), and global δ^18^O data from Zachos *et al*. (2008)). Possible correlation was analysed using generalised least squares (GLS) regressions, incorporating a first-order autoregressive model, as well as ordinary least squares (OLS) regressions using untransformed data (assuming no serial correlation). *Significant at alpha = 0.05.

| Prokoph (Early Triassic -recent): tropical palaeotemperatures |  |  |  |  |  |  |  |  |  |  |  |  |  |  |  |  |
| --- | --- | --- | --- | --- | --- | --- | --- | --- | --- | --- | --- | --- | --- | --- | --- | --- |
| N | Maximum size |  |  |  |  |  |  |  | Mean size |  |  |  |  |  |  |  |
|  | GLS |  |  |  | OLS (untransformed) |  |  |  | GLS |  |  |  | OLS (untransformed) |  |  |  |
|  | Phi | Int. | Slope | AIC | R<sup>2</sup> | Int. | Slope | AIC | Phi | Int. | Slope | AIC | R<sup>2</sup> | Int. | Slope | AIC |
| 26 | 0.553 | 2.32 | 0.043<br>(0.504) | 4.843 | 0.011 | 2.366 | 0.075<br>(0.264) | 11.094 | 0.64 | 1.978 | 0.029<br>(0.453) | -21.012 | 0.049 | 2.023 | 0.065<br>(0.142) | -11.564 |
| Prokoph (Early Triassic -recent): temperate palaeotemperatures |  |  |  |  |  |  |  |  |  |  |  |  |  |  |  |  |
| N | Maximum size |  |  |  |  |  |  |  | Mean size |  |  |  |  |  |  |  |
|  | GLS |  |  |  | OLS (untransformed) |  |  |  | GLS |  |  |  | OLS (untransformed) |  |  |  |
|  | Phi | Int. | Slope | AIC | R<sup>2</sup> | Int. | Slope | AIC | Phi | Int. | Slope | AIC | R<sup>2</sup> | Int. | Slope | AIC |
| 23 | 0.354 | 2.291 | -0.065<br>(0.232) | -6.825 | 0.022 | 2.291 | -0.06<br>(0.232) | -6.129 | 0.523 | 1.947 | -0.042<br>(0.299) | -21.071 | -0.037 | 1.967 | -0.017<br>(0.65) | -17.942 |
| Zachos (Late Cretaceous - recent): global palaeotemperatures |  |  |  |  |  |  |  |  |  |  |  |  |  |  |  |  |
| N | Maximum size |  |  |  |  |  |  |  | Mean size |  |  |  |  |  |  |  |
|  | GLS |  |  |  | OLS (untransformed) |  |  |  | GLS |  |  |  | OLS (untransformed) |  |  |  |
|  | Phi | Int. | Slope | AIC | R<sup>2</sup> | Int. | Slope | AIC | Phi | Int. | Slope | AIC | R<sup>2</sup> | Int. | Slope | AIC |
| 10 | 0.209 | 2.228 | 0.068<br>(0.366) | -0.829 | -0.011 | 2.236 | 0.06<br>(0.371) | -2.397 | -0.157 | 1.964 | 0.06*<br>(0.007) | -24.96 | 0.502 | 1.965 | 0.06*<br>(0.013) | -26.706 |

**Table S8.** Results of regressions of body size proxy (maximum and mean log-transformed DCL, using only non-marine species in the dataset) on the palaeotemperature proxies (δ^18^O data for tropical and temperate regions from Prokoph *et al*. (2008), and global δ^18^O data from Zachos *et al*. (2008)). Possible correlation was analysed using generalised least squares (GLS) regressions, incorporating a first-order autoregressive model, as well as ordinary least squares (OLS) regressions using untransformed data (assuming no serial correlation). *Significant at alpha = 0.05.

| Prokoph (Early Triassic -recent): tropical palaeotemperatures |  |  |  |  |  |  |  |  |  |  |  |  |  |  |  |  |
| --- | --- | --- | --- | --- | --- | --- | --- | --- | --- | --- | --- | --- | --- | --- | --- | --- |
| N | Maximum size |  |  |  |  |  |  |  | Mean size |  |  |  |  |  |  |  |
|  | GLS |  |  |  | OLS (untransformed) |  |  |  | GLS |  |  |  | OLS (untransformed) |  |  |  |
|  | Phi | Int. | Slope | AIC | R<sup>2</sup> | Int. | Slope | AIC | Phi | Int. | Slope | AIC | R<sup>2</sup> | Int. | Slope | AIC |
| 26 | 0.563 | 2.753 | 0.024<br>(0.763) | 15.858 | -0.011 | 2.82 | 0.069<br>(0.406) | 22.285 | 0.624 | 2.339 | 0.017<br>(0.725) | -8.74 | -0.018 | 2.366 | 0.04<br>(0.466) | 0.623 |
| Prokoph (Early Triassic -recent): temperate palaeotemperatures |  |  |  |  |  |  |  |  |  |  |  |  |  |  |  |  |
| N | Maximum size |  |  |  |  |  |  |  | Mean size |  |  |  |  |  |  |  |
|  | GLS |  |  |  | OLS (untransformed) |  |  |  | GLS |  |  |  | OLS (untransformed) |  |  |  |
|  | Phi | Int. | Slope | AIC | R<sup>2</sup> | Int. | Slope | AIC | Phi | Int. | Slope | AIC | R<sup>2</sup> | Int. | Slope | AIC |
| 23 | 0.317 | 2.76 | -0.071<br>(0.312) | 5.801 | 0.003 | 2.762 | -0.066<br>(0.309) | 5.997 | 0.518 | 2.32 | -0.046<br>(0.387) | -7.779 | -0.033 | 2.335 | -0.027<br>(0.6) | -4.075 |
| Zachos (Late Cretaceous - recent): global palaeotemperatures |  |  |  |  |  |  |  |  |  |  |  |  |  |  |  |  |
| N | Maximum size |  |  |  |  |  |  |  | Mean size |  |  |  |  |  |  |  |
|  | GLS |  |  |  | OLS (untransformed) |  |  |  | GLS |  |  |  | OLS (untransformed) |  |  |  |
|  | Phi | Int. | Slope | AIC | R<sup>2</sup> | Int. | Slope | AIC | Phi | Int. | Slope | AIC | R<sup>2</sup> | Int. | Slope | AIC |
| 10 | -0.083 | 2.633 | 0.095<br>(0.172) | 0.504 | 0.104 | 2.633 | 0.096<br>(0.189) | -1.426 | -0.089 | 2.345 | 0.07*<br>(0.027) | -16.045 | 0.376 | 2.346 | 0.07*<br>(0.034) | -18.272 |

**Table S9.** Results of regressions of body size proxy (maximum and mean log-transformed ODCL, using only crocodylian species in the dataset) on the palaeotemperature proxies (global δ^18^O data from Zachos *et al*. (2008), from the Late Cretaceous to Recent). Possible correlation was analysed using generalised least squares (GLS) regressions, incorporating a first-order autoregressive model, as well as ordinary least squares (OLS) regressions using untransformed data (assuming no serial correlation). *Significant at alpha = 0.05.

| Zachos (Late Cretaceous - recent): global palaeotemperatures |  |  |  |  |  |  |  |  |  |  |  |  |  |  |  |  |
| --- | --- | --- | --- | --- | --- | --- | --- | --- | --- | --- | --- | --- | --- | --- | --- | --- |
| N | Maximum size |  |  |  |  |  |  |  | Mean size |  |  |  |  |  |  |  |
|  | GLS |  |  |  | OLS (untransformed) |  |  |  | GLS |  |  |  | OLS (untransformed) |  |  |  |
|  | Phi | Int. | Slope | AIC | R<sup>2</sup> | Int. | Slope | AIC | Phi | Int. | Slope | AIC | R<sup>2</sup> | Int. | Slope | AIC |
| 10 | 0.19 | 2.133 | 0.121*<br>(0.017) | -11.989 | 0.554 | 2.124 | 0.127*<br>(0.008) | -13.662 | -0.297 | 1.98 | 0.075*<br>(0.0003) | -29.953 | 0.698 | 1.987 | 0.07*<br>(0.001) | -31.137 |

**Table S10.** Results of regressions of body size proxy (maximum and mean log-transformed DCL, using only crocodylian species in the dataset) on the palaeotemperature proxies (global δ^18^O data from Zachos *et al*. (2008), from the Late Cretaceous to Recent). Possible correlation was analysed using generalised least squares (GLS) regressions, incorporating a first-order autoregressive model, as well as ordinary least squares (OLS) regressions using untransformed data (assuming no serial correlation). *Significant at alpha = 0.05.

| Zachos (Late Cretaceous - recent): global palaeotemperatures |  |  |  |  |  |  |  |  |  |  |  |  |  |  |  |  |
| --- | --- | --- | --- | --- | --- | --- | --- | --- | --- | --- | --- | --- | --- | --- | --- | --- |
| N | Maximum size |  |  |  |  |  |  |  | Mean size |  |  |  |  |  |  |  |
|  | GLS |  |  |  | OLS (untransformed) |  |  |  | GLS |  |  |  | OLS (untransformed) |  |  |  |
|  | Phi | Int. | Slope | AIC | R<sup>2</sup> | Int. | Slope | AIC | Phi | Int. | Slope | AIC | R<sup>2</sup> | Int. | Slope | AIC |
| 10 | -0.215 | 2.618 | 0.165*<br>(0.001) | -10.724 | 0.632 | 2.627 | 0.157*<br>(0.003) | -12.355 | -0.235 | 2.386 | 0.105*<br>(0.0007) | -20.748 | 0.647 | 2.395 | 0.098*<br>(0.003) | -22.325 |

**Table S11.** Results of regressions of body size proxy (maximum and mean log-transformed ODCL, using only notosuchian species in the dataset) on the palaeotemperature proxies (tropical δ^18^O data from Prokoph *et al*. (2008), from the Aptian to the Eocene). Possible correlation was analysed using generalised least squares (GLS) regressions, incorporating a first-order autoregressive model, as well as ordinary least squares (OLS) regressions using untransformed data (assuming no serial correlation). *Significant at alpha = 0.05.

| Prokoph (Aptian - Eocene): tropical palaeotemperatures |  |  |  |  |  |  |  |  |  |  |  |  |  |  |  |  |
| --- | --- | --- | --- | --- | --- | --- | --- | --- | --- | --- | --- | --- | --- | --- | --- | --- |
| N | Maximum size |  |  |  |  |  |  |  | Mean size |  |  |  |  |  |  |  |
|  | GLS |  |  |  | OLS (untransformed) |  |  |  | GLS |  |  |  | OLS (untransformed) |  |  |  |
|  | Phi | Int. | Slope | AIC | R<sup>2</sup> | Int. | Slope | AIC | Phi | Int. | Slope | AIC | R<sup>2</sup> | Int. | Slope | AIC |
| 10 | 0.272 | 2.114 | -0.013<br>(0.812) | -5.557 | -0.115 | 2.118 | -0.014<br>(0.798) | -6.786 | 0.702 | 1.925 | -0.029<br>(0.472) | -11.724 | -0.122 | 1.957 | -0.005<br>(0.904) | -10.071 |

**Table S12.** Results of regressions of body size proxy (maximum and mean log-transformed DCL, using only notosuchian species in the dataset) on the palaeotemperature proxies (tropical δ^18^O data from Prokoph *et al*. (2008), from the Aptian to the Eocene). Possible correlation was analysed using generalised least squares (GLS) regressions, incorporating a first-order autoregressive model, as well as ordinary least squares (OLS) regressions using untransformed data (assuming no serial correlation). *Significant at alpha = 0.05.

| Prokoph (Aptian - Eocene): tropical palaeotemperatures |  |  |  |  |  |  |  |  |  |  |  |  |  |  |  |  |
| --- | --- | --- | --- | --- | --- | --- | --- | --- | --- | --- | --- | --- | --- | --- | --- | --- |
| N | Maximum size |  |  |  |  |  |  |  | Mean size |  |  |  |  |  |  |  |
|  | GLS |  |  |  | OLS (untransformed) |  |  |  | GLS |  |  |  | OLS (untransformed) |  |  |  |
|  | Phi | Int. | Slope | AIC | R<sup>2</sup> | Int. | Slope | AIC | Phi | Int. | Slope | AIC | R<sup>2</sup> | Int. | Slope | AIC |
| 10 | 0.06 | 2.622 | -0.014<br>(0.699) | -12.63 | -0.092 | 2.618 | -0.017<br>(0.64) | -14.601 | 0.758 | 2.313 | -0.055<br>(0.3) | -6.073 | -0.123 | 2.355 | -0.005<br>(0.928) | -3.54 |

**Table S13.** Results of regressions of body size proxy (maximum and mean log-transformed ODCL, using only thalattosuchian species in the dataset) on the palaeotemperature proxies (tropical δ^18^O data from Prokoph *et al*. (2008), for the Jurassic). Possible correlation was analysed using generalised least squares (GLS) regressions, incorporating a first-order autoregressive model, as well as ordinary least squares (OLS) regressions using untransformed data (assuming no serial correlation). *Significant at alpha = 0.05.

| Prokoph (Jurassic): tropical palaeotemperatures |  |  |  |  |  |  |  |  |  |  |  |  |  |  |  |  |
| --- | --- | --- | --- | --- | --- | --- | --- | --- | --- | --- | --- | --- | --- | --- | --- | --- |
| N | Maximum size |  |  |  |  |  |  |  | Mean size |  |  |  |  |  |  |  |
|  | GLS |  |  |  | OLS (untransformed) |  |  |  | GLS |  |  |  | OLS (untransformed) |  |  |  |
|  | Phi | Int. | Slope | AIC | R<sup>2</sup> | Int. | Slope | AIC | Phi | Int. | Slope | AIC | R<sup>2</sup> | Int. | Slope | AIC |
| 7 | 0.809 | 2.396 | -0.051<br>(0.308) | -5.062 | 0.059 | 2.322 | -0.11<br>(0.292) | -2.309 | -0.184 | 2.224 | -0.038<br>(0.455) | -10.311 | -0.09 | 2.232 | -0.033<br>(0.509) | -12.067 |
| Prokoph (Jurassic): temperate palaeotemperatures |  |  |  |  |  |  |  |  |  |  |  |  |  |  |  |  |
| N | Maximum size |  |  |  |  |  |  |  | Mean size |  |  |  |  |  |  |  |
|  | GLS |  |  |  | OLS (untransformed) |  |  |  | GLS |  |  |  | OLS (untransformed) |  |  |  |
|  | Phi | Int. | Slope | AIC | R<sup>2</sup> | Int. | Slope | AIC | Phi | Int. | Slope | AIC | R<sup>2</sup> | Int. | Slope | AIC |
| 7 | 0.808 | 2.526 | 0.074<br>(0.098) | -7.658 | 0.452 | 2.633 | 0.152<br>(0.058) | -6.096 | -0.369 | 2.366 | 0.082*<br>(0.003) | -22.184 | 0.778 | 2.369 | 0.086*<br>(0.005) | -23.214 |

**Table S14.** Results of regressions of body size proxy (maximum and mean log-transformed DCL, using only thalattosuchian species in the dataset) on the palaeotemperature proxies (tropical δ^18^O data from Prokoph *et al*. (2008), for the Jurassic). Possible correlation was analysed using generalised least squares (GLS) regressions, incorporating a first-order autoregressive model, as well as ordinary least squares (OLS) regressions using untransformed data (assuming no serial correlation). *Significant at alpha = 0.05.

| Prokoph (Jurassic): tropical palaeotemperatures |  |  |  |  |  |  |  |  |  |  |  |  |  |  |  |  |
| --- | --- | --- | --- | --- | --- | --- | --- | --- | --- | --- | --- | --- | --- | --- | --- | --- |
| N | Maximum size |  |  |  |  |  |  |  | Mean size |  |  |  |  |  |  |  |
|  | GLS |  |  |  | OLS (untransformed) |  |  |  | GLS |  |  |  | OLS (untransformed) |  |  |  |
|  | Phi | Int. | Slope | AIC | R<sup>2</sup> | Int. | Slope | AIC | Phi | Int. | Slope | AIC | R<sup>2</sup> | Int. | Slope | AIC |
| 7 | 0.661 | 2.856 | -0.054<br>(0.176) | -10.26 | 0.192 | 2.814 | -0.088<br>(0.179) | -9.432 | -0.124 | 2.727 | -0.042<br>(0.391) | -10.851 | -0.022 | 2.728 | -0.041<br>(0.394) | -12.753 |
| Prokoph (Jurassic): temperate palaeotemperatures |  |  |  |  |  |  |  |  |  |  |  |  |  |  |  |  |
| N | Maximum size |  |  |  |  |  |  |  | Mean size |  |  |  |  |  |  |  |
|  | GLS |  |  |  | OLS (untransformed) |  |  |  | GLS |  |  |  | OLS (untransformed) |  |  |  |
|  | Phi | Int. | Slope | AIC | R<sup>2</sup> | Int. | Slope | AIC | Phi | Int. | Slope | AIC | R<sup>2</sup> | Int. | Slope | AIC |
| 7 | 0.553 | 2.995 | 0.07<br>(0.069) | -12.556 | 0.563 | 3.046 | 0.107*<br>(0.031) | -13.734 | 0.582 | 2.852 | 0.072<br>(0.056) | -12.788 | 0.162 | 2.839 | 0.051<br>(0.201) | -14.155 |

**Table S15.** Results of regressions of body size proxy (maximum and mean log-transformed ODCL, using only tethysuchian species in the dataset) on the palaeotemperature proxies (tropical δ^18^O data from Prokoph *et al*. (2008), from the Late Jurassic to the Eocene). Possible correlation was analysed using generalised least squares (GLS) regressions, incorporating a first-order autoregressive model, as well as ordinary least squares (OLS) regressions using untransformed data (assuming no serial correlation). *Significant at alpha = 0.05.

| Prokoph (Late Jurassic – Eocene): tropical palaeotemperatures |  |  |  |  |  |  |  |  |  |  |  |  |  |  |  |  |
| --- | --- | --- | --- | --- | --- | --- | --- | --- | --- | --- | --- | --- | --- | --- | --- | --- |
| N | Maximum size |  |  |  |  |  |  |  | Mean size |  |  |  |  |  |  |  |
|  | GLS |  |  |  | OLS (untransformed) |  |  |  | GLS |  |  |  | OLS (untransformed) |  |  |  |
|  | Phi | Int. | Slope | AIC | R<sup>2</sup> | Int. | Slope | AIC | Phi | Int. | Slope | AIC | R<sup>2</sup> | Int. | Slope | AIC |
| 13 | -0.554 | 2.243 | -0.145*<br>(0.004) | -5.113 | 0.138 | 2.288 | -0.108<br>(0.115) | -2.267 | -0.448 | 2.096 | -0.154*<br>(0.0002) | -15.18 | 0.493 | 2.116 | -0.142*<br>(0.004) | -14.409 |
| Prokoph (Late Jurassic – Eocene): temperate palaeotemperatures |  |  |  |  |  |  |  |  |  |  |  |  |  |  |  |  |
| N | Maximum size |  |  |  |  |  |  |  | Mean size |  |  |  |  |  |  |  |
|  | GLS |  |  |  | OLS (untransformed) |  |  |  | GLS |  |  |  | OLS (untransformed) |  |  |  |
|  | Phi | Int. | Slope | AIC | R<sup>2</sup> | Int. | Slope | AIC | Phi | Int. | Slope | AIC | R<sup>2</sup> | Int. | Slope | AIC |
| 13 | -0.223 | 2.276 | -0.142<br>(0.051) | -1.969 | 0.202 | 2.27 | -0.15<br>(0.069) | -3.281 | 0.113 | 2.165 | -0.129<br>(0.063) | -7.028 | 0.226 | 2.163 | -0.127<br>(0.057) | -8.891 |

**Table S16.** Results of regressions of body size proxy (maximum and mean log-transformed DCL, using only tethysuchian species in the dataset) on the palaeotemperature proxies (tropical δ^18^O data from Prokoph *et al*. (2008), from the Late Jurassic to the Eocene). Possible correlation was analysed using generalised least squares (GLS) regressions, incorporating a first-order autoregressive model, as well as ordinary least squares (OLS) regressions using untransformed data (assuming no serial correlation). *Significant at alpha = 0.05.

| Prokoph (Late Jurassic – Eocene): tropical palaeotemperatures |  |  |  |  |  |  |  |  |  |  |  |  |  |  |  |  |
| --- | --- | --- | --- | --- | --- | --- | --- | --- | --- | --- | --- | --- | --- | --- | --- | --- |
| N | Maximum size |  |  |  |  |  |  |  | Mean size |  |  |  |  |  |  |  |
|  | GLS |  |  |  | OLS (untransformed) |  |  |  | GLS |  |  |  | OLS (untransformed) |  |  |  |
|  | Phi | Int. | Slope | AIC | R<sup>2</sup> | Int. | Slope | AIC | Phi | Int. | Slope | AIC | R<sup>2</sup> | Int. | Slope | AIC |
| 12 | 0.53 | 3.02 | -0.004<br>(0.914) | -11.772 | -0.061 | 2.993 | -0.03<br>(0.559) | -10.039 | 0.483 | 2.848 | -0.045<br>(0.327) | -10.28 | 0.075 | 2.814 | -0.07<br>(0.198) | -9.441 |
| Prokoph (Late Jurassic – Eocene): temperate palaeotemperatures |  |  |  |  |  |  |  |  |  |  |  |  |  |  |  |  |
| N | Maximum size |  |  |  |  |  |  |  | Mean size |  |  |  |  |  |  |  |
|  | GLS |  |  |  | OLS (untransformed) |  |  |  | GLS |  |  |  | OLS (untransformed) |  |  |  |
|  | Phi | Int. | Slope | AIC | R<sup>2</sup> | Int. | Slope | AIC | Phi | Int. | Slope | AIC | R<sup>2</sup> | Int. | Slope | AIC |
| 12 | 0.528 | 2.941 | -0.08<br>(0.15) | -14.367 | 0.105 | 2.941 | -0.083<br>(0.16) | -12.088 | 0.503 | 2.799 | -0.105<br>(0.081) | -12.934 | 0.221 | 2.782 | -0.114<br>(0.069) | -11.507 |

**Table S17.** Results of regressions of log-transformed body length proxy (using all species in the ODCL cranial measurement dataset) on the palaeolatitudinal data. Possible correlation was analysed using ordinary least squares (OLS) and phylogenetic generalised least squares (PGLS) regressions. *Significant at alpha = 0.05.

| N | OLS |  |  |  | PGLS |  |  |  |
| --- | --- | --- | --- | --- | --- | --- | --- | --- |
|  | R<sup>2</sup> | Intercept | Slope | AIC | R<sup>2</sup> | Intercept | Slope | AIC |
| 195 | 0.013 | 2.13 | -0.002 (0.059) | 43.284 | 0.003 | 1.77 | -0.001 (0.194) | -39.972 |

**Table S18.** Results of regressions of log-transformed body length proxy (using all species in the DCL cranial measurement dataset) on the palaeolatitudinal data. Possible correlation was analysed using ordinary least squares (OLS) and phylogenetic generalised least squares (PGLS) regressions. *Significant at alpha = 0.05.

| N | OLS |  |  |  | PGLS |  |  |  |
| --- | --- | --- | --- | --- | --- | --- | --- | --- |
|  | R<sup>2</sup> | Intercept | Slope | AIC | R<sup>2</sup> | Intercept | Slope | AIC |
| 178 | 0.022 | 2.595 | -0.004* (0.024) | 150.74 | 0.019 | 2.195 | -0.002* (0.034) | -19.379 |

**Table S19.** Results of regressions of log-transformed body length proxy (using only marine species in the ODCL cranial measurement dataset) on the palaeolatitudinal data. Possible correlation was analysed using ordinary least squares (OLS) and phylogenetic generalised least squares (PGLS) regressions. *Significant at alpha = 0.05.

| N | OLS |  |  |  | PGLS |  |  |  |
| --- | --- | --- | --- | --- | --- | --- | --- | --- |
|  | R<sup>2</sup> | Intercept | Slope | AIC | R<sup>2</sup> | Intercept | Slope | AIC |
| 48 | -0.019 | 2.289 | -0.0008 (0.739) | -21.925 | 0.035 | 2.289 | -0.003 (0.105) | -36.771 |

**Table S20.** Results of regressions of log-transformed body length proxy (using only marine species in the DCL cranial measurement dataset) on the palaeolatitudinal data. Possible correlation was analysed using ordinary least squares (OLS) and phylogenetic generalised least squares (PGLS) regressions. *Significant at alpha = 0.05.

| N | OLS |  |  |  | PGLS |  |  |  |
| --- | --- | --- | --- | --- | --- | --- | --- | --- |
|  | R<sup>2</sup> | Intercept | Slope | AIC | R<sup>2</sup> | Intercept | Slope | AIC |
| 43 | 0.014 | 2.873 | -0.002 (0.211) | -28.625 | -0.014 | 2.662 | 0.001 (0.53) | -45.625 |

**Table S21.** Results of regressions of log-transformed body length proxy (using only non-marine species in the ODCL cranial measurement dataset) on the palaeolatitudinal data. Possible correlation was analysed using ordinary least squares (OLS) and phylogenetic generalised least squares (PGLS) regressions. *Significant at alpha = 0.05.

| N | OLS |  |  |  | PGLS |  |  |  |
| --- | --- | --- | --- | --- | --- | --- | --- | --- |
|  | R<sup>2</sup> | Intercept | Slope | AIC | R<sup>2</sup> | Intercept | Slope | AIC |
| 147 | 0.037 | 2.09 | -0.003* (0.01) | 14.567 | 0.028 | 1.836 | -0.002* (0.023) | -48.394 |

**Table S22.** Results of regressions of log-transformed body length proxy (using only non-marine species in the DCL cranial measurement dataset) on the palaeolatitudinal data. Possible correlation was analysed using ordinary least squares (OLS) and phylogenetic generalised least squares (PGLS) regressions. *Significant at alpha = 0.05.

| N | OLS |  |  |  | PGLS |  |  |  |
| --- | --- | --- | --- | --- | --- | --- | --- | --- |
|  | R<sup>2</sup> | Intercept | Slope | AIC | R<sup>2</sup> | Intercept | Slope | AIC |
| 135 | 0.036 | 2.508 | -0.005* (0.014) | 102.424 | 0.06 | 2.259 | -0.004* (0.002) | 13.242 |

**Table S23.**
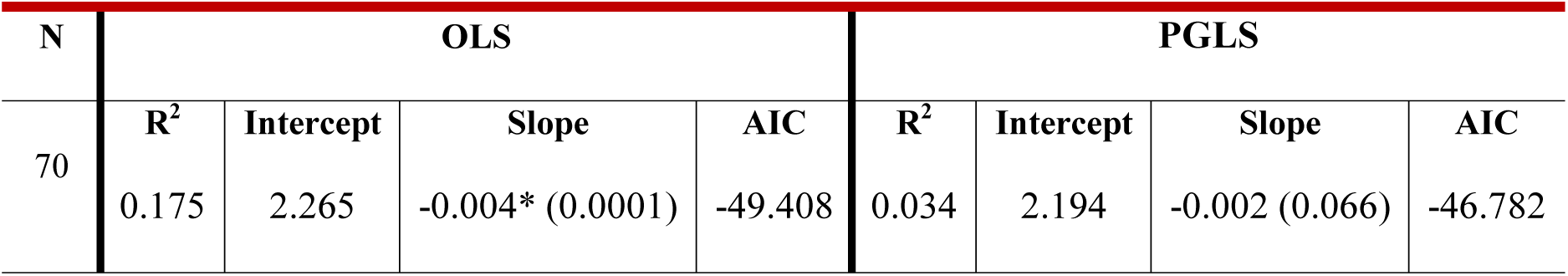
Results of regressions of log-transformed body length proxy (using only crocodylian species in the ODCL cranial measurement dataset) on the palaeolatitudinal data. Possible correlation was analysed using ordinary least squares (OLS) and phylogenetic generalised least squares (PGLS) regressions. *Significant at alpha = 0.05.

| N | OLS |  |  |  | PGLS |  |  |  |
| --- | --- | --- | --- | --- | --- | --- | --- | --- |
|  | R<sup>2</sup> | Intercept | Slope | AIC | R<sup>2</sup> | Intercept | Slope | AIC |
| 70 | 0.175 | 2.265 | -0.004* (0.0001) | -49.408 | 0.034 | 2.194 | -0.002 (0.066) | -46.782 |

**Table S24.** Results of regressions of log-transformed body length proxy (using only crocodylian species in the DCL cranial measurement dataset) on the palaeolatitudinal data. Possible correlation was analysed using ordinary least squares (OLS) and phylogenetic generalised least squares (PGLS) regressions. *Significant at alpha = 0.05.

| N | OLS |  |  |  | PGLS |  |  |  |
| --- | --- | --- | --- | --- | --- | --- | --- | --- |
|  | R<sup>2</sup> | Intercept | Slope | AIC | R<sup>2</sup> | Intercept | Slope | AIC |
| 64 | 0.178 | 2.81 | -0.007* (0.0003) | 8.976 | 0.045 | 2.744 | -0.004 (0.05) | -6.629 |

**Table S25.**
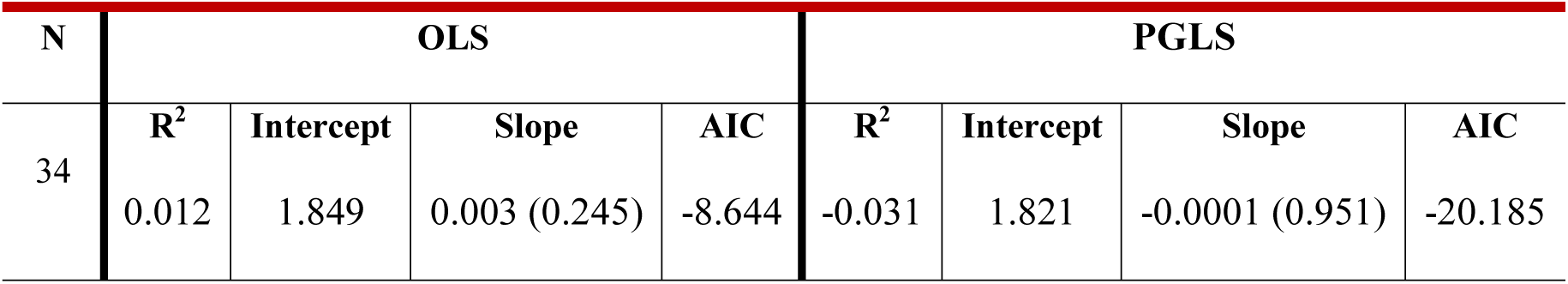
Results of regressions of log-transformed body length proxy (using only notosuchian species in the ODCL cranial measurement dataset) on the palaeolatitudinal data. Possible correlation was analysed using ordinary least squares (OLS) and phylogenetic generalised least squares (PGLS) regressions. *Significant at alpha = 0.05.

**Table S26.**
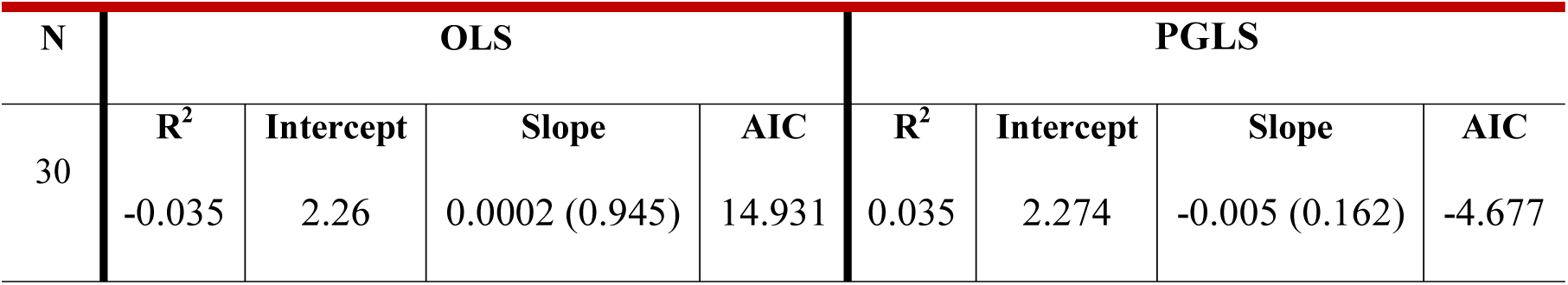
Results of regressions of log-transformed body length proxy (using only notosuchian species in the DCL cranial measurement dataset) on the palaeolatitudinal data. Possible correlation was analysed using ordinary least squares (OLS) and phylogenetic generalised least squares (PGLS) regressions. *Significant at alpha = 0.05.

**Table S27.** Results of regressions of log-transformed body length proxy (using only thalattosuchian species in the ODCL cranial measurement dataset) on the palaeolatitudinal data. Possible correlation was analysed using ordinary least squares (OLS) and phylogenetic generalised least squares (PGLS) regressions. *Significant at alpha = 0.05.

| N | OLS |  |  |  | PGLS |  |  |  |
| --- | --- | --- | --- | --- | --- | --- | --- | --- |
|  | R<sup>2</sup> | Intercept | Slope | AIC | R<sup>2</sup> | Intercept | Slope | AIC |
| 30 | -0.019 | 2.115 | 0.004 (0.509) | -4.607 | -0.035 | 2.156 | -0.0001 (0.977) | -19.849 |

**Table S28.** Results of regressions of log-transformed body length proxy (using only thalattosuchian species in the DCL cranial measurement dataset) on the palaeolatitudinal data. Possible correlation was analysed using ordinary least squares (OLS) and phylogenetic generalised least squares (PGLS) regressions. *Significant at alpha = 0.05.

| N | OLS |  |  |  | PGLS |  |  |  |
| --- | --- | --- | --- | --- | --- | --- | --- | --- |
|  | R<sup>2</sup> | Intercept | Slope | AIC | R<sup>2</sup> | Intercept | Slope | AIC |
| 26 | -0.004 | 2.579 | 0.004 (0.357) | -14.111 | 0.01 | 2.601 | 0.003 (0.273) | -28.503 |

**Table S29.** Results of regressions of log-transformed body length proxy (using only tethysuchian species in the ODCL cranial measurement dataset) on the palaeolatitudinal data. Possible correlation was analysed using ordinary least squares (OLS) and phylogenetic generalised least squares (PGLS) regressions. *Significant at alpha = 0.05.

| N | OLS |  |  |  | PGLS |  |  |  |
| --- | --- | --- | --- | --- | --- | --- | --- | --- |
|  | R<sup>2</sup> | Intercept | Slope | AIC | R<sup>2</sup> | Intercept | Slope | AIC |
| 16 | 0.251 | 2.468 | -0.009* (0.027) | -5.781 | 0.444 | 2.54 | -0.012* (0.002) | -6.853 |

**Table S30.** Results of regressions of log-transformed body length proxy (using only tethysuchian species in the DCL cranial measurement dataset) on the palaeolatitudinal data. Possible correlation was analysed using ordinary least squares (OLS) and phylogenetic generalised least squares (PGLS) regressions. *Significant at alpha = 0.05.

| N | OLS |  |  |  | PGLS |  |  |  |
| --- | --- | --- | --- | --- | --- | --- | --- | --- |
|  | R<sup>2</sup> | Intercept | Slope | AIC | R<sup>2</sup> | Intercept | Slope | AIC |
| 14 | -0.002 | 2.898 | -0.004 (0.345) | -1.387 | -0.084 | 2.904 | 0.001 (0.729) | -2.537 |

## References

1. Hutchinson GE, MacArthur RH. A theoretical ecological model of size distributions among species of animals. Am Nat. 1959;93:117–25.

2. Peters RH: The Ecological Implications of body size. New York: Cambridge University Press; 1983.

3. Calder WAI: Size, Function, and Life History. Cambridge: Harvard University Press; 1984.

4. Schmidt-Nielsen K: Scaling: Why is animal size so important? Cambridge: Cambridge University Press; 1984.

5. McKinney ML. Trends in body size evolution. In: McNamara KJ, editor. Evolutionary trends. Tucson: University of Arizona Press; 1990. p. 75–118.

6. McClain CR, Boyer AG. Biodiversity and body size are linked across metazoans. Proc R Soc B-Biol Sci. 2009;276:2209–15.

7. Cope ED. The origin of the fittest: essays on evolution. New York: D. Appleton and Company; 1887.

8. Cope ED. The primary factors of organic evolution. Chicago: Open Court Press; 1896.

9. Depéret CJJ. The transformations of the animal world. New York: D. Appleton and Company; 1909.

10. Newell ND. Phyletic size increase, an important trend illustrated by fossil invertebrates. Evolution. 1949;3:103–24.

11. Stanley SM. An explanation for Cope’s rule. Evolution. 1973;27:1–26.

12. Price SA, Hopkins SS. The macroevolutionary relationship between diet and body mass across mammals. Biol J Linnean Soc. 2015;115:173–84.

13. Raup DM. Testing the fossil record for evolutionary progress. In: Nitecki MH, editor. Evolutionary progress. Chicago: University of Chicago Press; 1988. p. 293–317.

14. Alroy J. Cope’s rule and the dynamics of body mass evolution in North American fossil mammals. Science. 1998;280:731–4.

15. Smith FA, Boyer AG, Brown JH, Costa DP, Dayan T, Ernest SM, Evans AR, Fortelius M, Gittleman JL, Hamilton MJ, et al. The evolution of maximum body size of terrestrial mammals. Science. 2010;330:1216–9.

16. Venditti C, Meade A, Pagel M. Multiple routes to mammalian diversity. Nature. 2011;479:393–6.

17. Heim NA, Knope ML, Schaal EK, Wang SC, Payne JL. Cope’s rule in the evolution of marine animals. Science. 2015;347:867–70.

18. Laurin M. The evolution of body size, Cope’s rule and the origin of amniotes. Syst Biol. 2004;53:594–622.

19. Benson RBJ, Frigot RA, Goswami A, Andres B, Butler RJ. Competition and constraint drove Cope’s rule in the evolution of giant flying reptiles. Nat Commun. 2014;5:3567.

20. Alberdi MT, Prado JL, Ortiz-Jaureguizar E. Patterns of body size changes in fossil and living Equini (Perissodactyla). Biol J Linnean Soc. 1995;54:349–70.

21. Smith FA, Lyons SK. How big should a mammal be? A macroecological look at mammalian body size over space and time. Philos Trans R Soc Lond B-Biol Sci. 2011;366:2364–78.

22. Saarinen JJ, Boyer AG, Brown JH, Costa DP, Ernest SM, Evans AR, Fortelius M, Gittleman JL, Hamilton MJ. Harding LE, et al. Patterns of maximum body size evolution in Cenozoic land mammals: eco-evolutionary processes and abiotic forcing. Proc R Soc B-Biol Sci. 2014;281:20132049.

23. Churchill M, Clementz MT, Kohno N. Cope’s rule and the evolution of body size in Pinnipedimorpha (Mammalia: Carnivora). Evolution. 2015;69:201–15.

24. Gearty W, McClain CR, Payne JL. Energetic tradeoffs control the size distribution of aquatic mammals. Proc Natl Acad Sci USA. 2018;115:4194–9.

25. Burness GP, Diamond J, Flannery T. Dinosaurs, dragons, and dwarfs: the evolution of maximal body size. Proc Natl Acad Sci USA. 2001;98:14518–23.

26. Hone DWE, Dyke GJ, Haden M, Benton MJ. Body size evolution in Mesozoic birds. J Evol Biol. 2008;21:618–24.

27. Carrano MT. Body-size evolution in the Dinosauria. In: Carrano MT, Blob RW, Gaudin T, Wibble JR, editors. Amniote paleobiology: perspectives on the evolution of mammals, birds, and reptiles. Chicago: University of Chicago Press; 2006. p. 225–68.

28. Turner AH, Pol D, Clarke JA, Erickson GM, Norell MA. A basal dromaeosaurid and size evolution preceding avian flight. Science. 2007;317:1378–81.

29. Butler RJ, Goswami A. Body size evolution in Mesozoic birds: little evidence for Cope’s rule. J Evol Biol. 2008;21:1673–82.

30. Lee MS, Cau A, Naish D, Dyke GJ. Sustained miniaturization and anatomical innovation in the dinosaurian ancestors of birds. Science. 2014;345:562–566.

31. Benson RBJ, Campione NE, Carrano MT, Mannion PD, Sullivan C, Upchurch P, Evans DC. Rates of dinosaur body mass evolution indicate 170 million years of sustained ecological innovation on the avian stem lineage. PLoS Biol. 2014;12:e1001853.

32. Carballido JL, Pol D, Otero A, Cerda IA, Salgado L, Garrido AC, Ramezani J, Cúneo NR, Krause JM. A new giant titanosaur sheds light on body mass evolution among sauropod dinosaurs. Proc R Soc B-Biol Sci. 2017;284:20171219.

33. Benson RBJ, Hunt G, Carrano MT, Campione N. Cope’s rule and the adaptive landscape of dinosaur body size evolution. Palaeontology. 2018;61:13–48.

34. Bronzati M, Montefeltro FC, Langer MC. A species-level supertree of Crocodyliformes. Hist Biol. 2012;24:598–606.

35. Bronzati M, Montefeltro FC, Langer MC. Diversification events and the effects of mass extinctions on Crocodyliformes evolutionary history. R Soc Open Sci. 2015;2:140385.

36. Mannion PD, Benson RBJ, Carrano MT, Tennant JP, Judd J, Butler RJ. Climate constrains the evolutionary history and biodiversity of crocodylians. Nat Commun. 2015;6:8438.

37. Markwick PJ. Crocodilian diversity in space and time: the role of climate in paleoecology and its implication for understanding K/T extinctions. Paleobiology. 1998;24:470–97.

38. Langston W. The crocodilian skull in historical perspective. In: Gans C, Parsons TS, editors. Biology of the Reptilia. London: Academic Press; 1973 p. 263–84.

39. Brochu CA. Crocodylian snouts in space and time: phylogenetic approaches toward adaptive radiation. Am Zool. 2001;41:564–85.

40. Sadleir RW, Makovicky PJ. Cranial shape and correlated characters in crocodilian evolution. J Evol Biol. 2008;21:1578–96.

41. Stubbs TL, Pierce SE, Rayfield EJ, Anderson PS. Morphological and biomechanical disparity of crocodile-line archosaurs following the end-Triassic extinction. Proc R Soc B-Biol Sci. 2013;280:20131940.

42. Toljagić, Butler RJ. Triassic–Jurassic mass extinction as trigger for the Mesozoic radiation of crocodylomorphs. Biol Lett. 2013;9:20130095.

43. Wilberg EW. Investigating patterns of crocodyliform cranial disparity through the Mesozoic and Cenozoic. Zool J Linn Soc. 2017;181:189–208.

44. Godoy PL, Ferreira GS, Montefeltro FC, Vila Nova BC, Butler RJ, Langer MC. Evidence for heterochrony in the cranial evolution of fossil crocodyliforms. Palaeontology. 2018;61:543–58.

45. Turner AH, Nesbitt SJ. Body size evolution during the Triassic archosauriform radiation. Geol Soc Spec Publ. 2013;379:573–97.

46. Young MT, Bell MA, Andrade MB, Brusatte SL. Body size estimation and evolution in metriorhynchid crocodylomorphs: implications for species diversification and niche partitioning. Zool J Linn Soc. 2011;163:1199–216.

47. Allsteadt J, Lang JW. Incubation temperature affects body size and energy reserves of hatchling American alligators (*Alligator mississippiensis*). Physiol Zool. 1995;68:76–97.

48. Markwick PJ. Fossil crocodilians as indicators of Late Cretaceous and Cenozoic climates: implications for using palaeontological data in reconstructing palaeoclimate. Palaeogeogr Palaeocl. 1998;137:205–71.

49. Delfino M, de Vos J. A giant crocodile in the Dubois Collection from the Pleistocene of Kali Gedeh (Java). Integr Zool. 2014;9:141–7.

50. Clark JM, Sues HD. Two new basal crocodylomorph archosaurs from the Lower Jurassic and the monophyly of the Sphenosuchia. Zool J Linn Soc. 2002;136:77–95.

51. Clark JM, Sues HD, Berman DS. A new specimen of *Hesperosuchus agilis* from the Upper Triassic of New Mexico and the interrelationships of basal crocodylomorph archosaurs. J Vertebr Paleontol. 2001;20:683–704.

52. Erickson GM, Brochu CA. How the ‘terror crocodile’ grew so big. Nature. 1999;398:205–6.

53. Sereno PC, Larsson HC, Sidor CA, Gado B. The giant crocodyliform *Sarcosuchus* from the Cretaceous of Africa. Science. 2001;294:1516–19.

54. Ross JP. Crocodiles: Status survey and conservation action plan 2nd ed. Gland (Switzerland), Cambridge (UK): IUCN/SSC Crocodile Specialist Group; 1998.

55. Grigg GC, Seebacher F, Franklin CE. Crocodilian Biology and Evolution. Chipping Norton: Surrey Beatty & Sons; 2001.

56. McShea DW. Mechanisms of large=scale evolutionary trends. Evolution. 1994;48:1747–63.

57. Felsenstein, J. Phylogenies and the comparative method. Am Nat. 1985;125:1– 15.

58. Hansen TF. Stabilizing selection and the comparative analysis of adaptation. Evolution; 1997;51:1341–51.

59. Pennell MW, Harmon LJ. An integrative view of phylogenetic comparative methods: connections to population genetics, community ecology, and paleobiology. Ann NY Acad Sci. 2013;1289:90–105.

60. MacFadden BJ. Fossil horses from “Eohippus” (*Hyracotherium*) to *Equus*: scaling, Cope’s law, and the evolution of body size. Paleobiology. 1986;12:355–69.

61. Butler MA, King AA. Phylogenetic comparative analysis: a modeling approach for adaptive evolution. Am Nat. 2004;164:683–95.

62. Hunt G, Carrano MT. Models and methods for analyzing phenotypic evolution in lineages and clades. In: Alroy J, Hunt G, editors. Quantitative methods in Paleobiology. New Haven: The Paleontological Society Papers; 2010. p. 245–69.

63. Hunt G. Measuring rates of phenotypic evolution and the inseparability of tempo and mode. Paleobiology. 2012;38:351–73.

64. Slater GJ. Phylogenetic evidence for a shift in the mode of mammalian body size evolution at the Cretaceous=Palaeogene boundary. Methods Ecol Evol. 2013;4:734–44.

65. Slater GJ. Iterative adaptive radiations of fossil canids show no evidence for diversity-dependent trait evolution. Proc Natl Acad Sci USA. 2015;112:4897–902.

66. Cooper N, Purvis A. Body size evolution in mammals: complexity in tempo and mode. Am Nat. 2010;175:727–38.

67. Sookias RB, Butler RJ, Benson RBJ. Rise of dinosaurs reveals major body-size transitions are driven by passive processes of trait evolution. Proc R Soc B-Biol Sci. 2012;279:2180–7.

68. Hunt G. Evolutionary patterns within fossil lineages: model-based assessment of modes, rates, punctuations and process. In: Bambach RK, Kelley PH, editors. From Evolution to geobiology: research questions driving Paleontology at the start of a new century. New Haven: The Paleontological Society Papers; 2008. p. 117–31.

69. Hunt G. Gradual or pulsed evolution: when should punctuational explanations be preferred? Paleobiology. 2008;34:360–77.

70. Hunt G, Hopkins MJ, Lidgard S. Simple versus complex models of trait evolution and stasis as a response to environmental change. Proc Natl Acad Sci USA. 2015;112:4885–90.

71. Mahler DL, Ingram T. Phylogenetic comparative methods for studying clade-wide convergence. In: Garamszegi LZ, editor. Modern phylogenetic comparative methods and their application in evolutionary biology. Berlin: Springer; 2014. p. 425– 50.

72. Khabbazian M, Kriebel R, Rohe K, Ané C. Fast and accurate detection of evolutionary shifts in Ornstein–Uhlenbeck models. Methods Ecol Evol. 2016;7:811–24.

73. Simpson GG. Tempo and mode in evolution. New York: Columbia University Press; 1944.

74. Simpson GG. Major features of evolution. New York: Columbia University Press; 1953.

75. Hansen TF. Adaptive landscapes and macroevolutionary dynamics. In: Svensson E, Calsbeek R, editors. The adaptive landscape in evolutionary biology. Oxford: Oxford University Press; 2012. p. 205–26.

76. Arnold SJ. Phenotypic evolution: the ongoing synthesis (American Society of Naturalists Address). Am Nat. 2014;183:729–46.

77. Arnold SJ, Pfrender ME, Jones AG. The adaptive landscape as a conceptual bridge between micro-and macroevolution. Genetica. 2001;112: 9–32.

78. Uyeda JC, Harmon LJ. A novel Bayesian method for inferring and interpreting the dynamics of adaptive landscapes from phylogenetic comparative data. Syst Biol. 2014;63:902–18.

79. Hansen TF, Martins EP. Translating between microevolutionary process and macroevolutionary patterns: the correlation structure of interspecific data. Evolution. 1996;50:1404–17.

80. Pagel M. Modelling the evolution of continuously varying characters on phylogenetic trees. In: MacLeod N, Forey PL, editors. Morphology, shape and phylogeny. London: Taylor & Francis; 2002. p. 269–86.

81. Felsenstein J. Phylogenies and quantitative characters. Annu Rev Ecol Syst. 1988;19:445–71.

82. Beaulieu JM, Jhwueng DC, Boettiger C, O’Meara BC. Modeling stabilizing selection: expanding the Ornstein–Uhlenbeck model of adaptive evolution. Evolution. 2012;66:2369–83.

83. Ingram T, Mahler DL. SURFACE: detecting convergent evolution from comparative data by fitting Ornstein=Uhlenbeck models with stepwise Akaike Information Criterion. Methods Ecol Evol. 2013;4:416–25.

84. Akaike H. A new look at the statistical model identification. IEEE Trans Autom Control. 1974;19:716–23.

85. Mahler DL, Ingram T, Revell LJ, Losos JB. Exceptional convergence on the macroevolutionary landscape in island lizard radiations. Science. 2013;341:292–5.

86. Davis AM, Unmack PJ, Pusey BJ, Pearson RG, Morgan DL. Evidence for a multi-peak adaptive landscape in the evolution of trophic morphology in terapontid fishes. Biol J Linnean Soc. 2014;113:623–34.

87. Brocklehurst N. Rates and modes of body size evolution in early carnivores and herbivores: a case study from Captorhinidae. PeerJ. 2016;4: e1555.

88. Young MT, Rabi M, Bell MA, Foffa D, Steel L, Sachs S, Peyer K. Big-headed marine crocodyliforms and why we must be cautious when using extant species as body length proxies for long-extinct relatives. Palaeontol Electron. 2016;19:1–14.

89. Webb GJW, Messel H. Morphometric analysis of *Crocodylus porosus* from the north coast of Arnhem Land, northern Australia. Aust J Zool. 1978;26:1–27.

90. Hall PM, Portier KM. Cranial morphometry of New Guinea crocodiles (*Crocodylus novaeguineae*): ontogenetic variation in relative growth of the skull and an assessment of its utility as a predictor of the sex and size of individuals. Herpetol Monogr. 1994;8:203–25.

91. Hurlburt GR, Heckert AB, Farlow JO. Body mass estimates of phytosaurs (Archosauria: Parasuchidae) from the Petrified Forest Formation (Chinle Group: Revueltian) based on skull and limb bone measurements. New Mex Mus Nat Hist Sci Bull. 2003;24:105–13.

92. Platt SG, Rainwater TR, Thorbjarnarson JB, Finger AG, Anderson TA, McMurry ST. Size estimation, morphometrics, sex ratio, sexual size dimorphism, and biomass of Morelet’s crocodile in northern Belize. Caribb J Sci. 2009;45:80–93.

93. Platt SG, Rainwater TR, Thorbjarnarson JB, Martin D. Size estimation, morphometrics, sex ratio, sexual size dimorphism, and biomass of *Crocodylus acutus* in the coastal zone of Belize. Salamandra. 2011;47:179–92.

94. Bustard HR, Singh LAK. Studies on the Indian Gharial *Gavialis gangeticus* (Gmelin) (Reptilia, Crocodilia) – I: Estimation of body length from scute length. Indian For. 1977;103:140–9.

95. Farlow JO, Hurlburt GR, Elsey RM, Britton AR, Langston W. Femoral dimensions and body size of Alligator mississippiensis: estimating the size of extinct mesoeucrocodylians. J Vertebr Paleontol. 2005;25:354–69.

96. Pol D, Leardi JM, Lecuona A, Krause M. Postcranial anatomy of *Sebecus icaeorhinus* (Crocodyliformes, Sebecidae) from the Eocene of Patagonia. J Vertebr Paleontol. 2012;32:328–54.

97. Godoy PL, Bronzati M, Eltink E, Marsola JCA, Cidade GM, Langer MC, Montefeltro FC. Postcranial anatomy of *Pissarrachampsa sera* (Crocodyliformes, Baurusuchidae) from the Late Cretaceous of Brazil: insights on lifestyle and phylogenetic significance. PeerJ. 2016;4:e2075.

98. Clark JM. Patterns of evolution in Mesozoic Crocodyliformes. In: Fraser NC, Sues HD, editors. In the shadow of the dinosaurs. early Mesozoic tetrapods. Cambridge: Cambridge University Press; 1994. p. 84–97.

99. Pol D, Gasparini Z. Skull anatomy of *Dakosaurus andiniensis* (Thalattosuchia: Crocodylomorpha) and the phylogenetic position of Thalattosuchia. J Syst Palaeontol. 2009;7:163–97.

100. Wilberg EW. What’s in an outgroup? The impact of outgroup choice on the phylogenetic position of Thalattosuchia (Crocodylomorpha) and the origin of Crocodyliformes. Syst Biol. 2015;64:621–37.

101. Herrera Y, Fernandez MS, Lamas SG, Campos L, Talevi M, Gasparini Z. Morphology of the sacral region and reproductive strategies of Metriorhynchidae: a counter-inductive approach. Earth Env Sci T R So Edinb. 2017;106:247–55.

102. Jouve S, Iarochene M, Bouya B, Amaghzaz M. A new species of *Dyrosaurus* (Crocodylomorpha, Dyrosauridae) from the early Eocene of Morocco: phylogenetic implications. Zool J Linn Soc. 2006;148:603–56.

103. Young MT, Andrade MB. What is *Geosaurus*? Redescription of *Geosaurus giganteus* (Thalattosuchia: Metriorhynchidae) from the Upper Jurassic of Bayern, Germany. Zool J Linn Soc. 2009;157:551–85.

104. Montefeltro FC, Larsson HC, França MA, Langer MC. A new neosuchian with Asian affinities from the Jurassic of northeastern Brazil. Naturwissenschaften. 2013;100:835–41.

105. Turner AH. A review of *Shamosuchus* and *Paralligator* (Crocodyliformes, Neosuchia) from the Cretaceous of Asia. PLoS One. 2015;10:e0118116.

106. Larsson HC, Sues HD. Cranial osteology and phylogenetic relationships of *Hamadasuchus rebouli* (Crocodyliformes: Mesoeucrocodylia) from the Cretaceous of Morocco. Zool J Linn Soc. 2007;149:533–67.

107. Bapst DW. Preparing paleontological datasets for phylogenetic comparative methods. In: Garamszegi LZ, editor. Modern phylogenetic comparative methods and their application in evolutionary biology. Berlin: Springer; 2014. p. 515–44.

108. Bapst DW. A stochastic rate=calibrated method for time=scaling phylogenies of fossil taxa. Methods Ecol Evol. 2013;4:724–33.

109. Bapst DW. Assessing the effect of time-scaling methods on phylogeny-based analyses in the fossil record. Paleobiology. 2014;40:331–51.

110. Stadler T. Sampling-through-time in birth–death trees. J Theor Biol. 2010;267:396–404.

111. Ronquist F, Klopfstein S, Vilhelmsen L, Schulmeister S, Murray DL, Rasnitsyn AP. A total-evidence approach to dating with fossils, applied to the early radiation of the Hymenoptera. Syst Biol. 2012;61:973–99.

112. Zhang C, Stadler T, Klopfstein S, Heath TA, Ronquist F. Total-evidence dating under the fossilized birth–death process. Syst Biol. 2015;65:228–49.

113. Matzke NJ, Wright A. Inferring node dates from tip dates in fossil Canidae: the importance of tree priors. Biol Lett. 2016;12:20160328.

114. Wright DF. Bayesian estimation of fossil phylogenies and the evolution of early to middle Paleozoic crinoids (Echinodermata). J Paleontol. 2017;91:799–814.

115. Ronquist F, Teslenko M, Van Der Mark P, Ayres DL, Darling A, Höhna S, Larget B, Liu L, Suchard MA, Huelsenbeck JP. MrBayes 3.2: efficient Bayesian phylogenetic inference and model choice across a large model space. Syst Biol. 2012;61:539–42.

116. Bapst DW. paleotree: an R package for paleontological and phylogenetic analyses of evolution. Methods Ecol Evol. 2012;3:803–07.

117. Irmis RB, Nesbitt SJ, Sues HD. Early Crocodylomorpha. Geol Soc Spec Publ. 2013;379:275–302.

118. Ezcurra MD, Butler RJ. The rise of the ruling reptiles and ecosystem recovery from the Permo-Triassic mass extinction Proc R Soc B-Biol Sci. 2018;285:20180361.

119. Bapst DW, Wright AM, Matzke NJ, Lloyd GT. Topology, divergence dates, and macroevolutionary inferences vary between different tip-dating approaches applied to fossil theropods (Dinosauria). Biol Lett. 2016;12:20160237.

120. Lloyd GT, Bapst DW, Friedman M, Davis KE. Probabilistic divergence time estimation without branch lengths: dating the origins of dinosaurs, avian flight and crown birds. Biol Lett. 2016;12:20160609.

121. R Core Team. R: a language and environment for statistical computing. Vienna: R Foundation for Statistical Computing; 2018. https://www.R-project.org/.

122. Harmon LJ, Weir JT, Brock CD, Glor RE, Challenger W. GEIGER: investigating evolutionary radiations. Bioinformatics. 2008;24:129–31.

123. Sugiura N. Further analysts of the data by Akaike’s information criterion and the finite corrections. Commun Stat–Theor M. 1978:7:13–26.

124. Burnham KP, Anderson DR. Model selection and multimodel inference: a practical information–theoretic approach. 2nd ed. New York: Springer; 2002.

125. Blomberg SP, Garland T, Ives AR. Testing for phylogenetic signal in comparative data: behavioral traits are more labile. Evolution. 2003;57:717–45.

126. Harmon LJ, Losos JB, Davies TJ, Gillespie RG, Gittleman JL, Bryan Jennings W, Kozak KH, McPeek MA, Moreno-Roark F, Near TJ, et al. Early bursts of body size and shape evolution are rare in comparative data. Evolution. 2010;64:2385–96.

127. Ho LST. Ané C. Intrinsic inference difficulties for trait evolution with Ornstein=Uhlenbeck models. Methods Ecol Evol. 2014;5:1133–46.

128. Cooper N, Thomas GH, Venditti C, Meade A, Freckleton RP. A cautionary note on the use of Ornstein Uhlenbeck models in macroevolutionary studies. Biol J Linn Soc. 2016;118:64–77.

129. Clavel J, Escarguel G, Merceron G. mvMORPH: an R package for fitting multivariate evolutionary models to morphometric data. Methods Ecol Evol. 2015;6:1311–9.

130. Beaulieu JM, O’Meara BC. OUwie: Analysis of Evolutionary Rates in an OU Framework. R package version 1.50. 2016. https://CRAN.R-project.org/package=OUwie.

131. Zachos JC, Dickens GR, Zeebe RE. An early Cenozoic perspective on greenhouse warming and carbon-cycle dynamics. Nature. 2008;451:279–83.

132. Prokoph A, Shields GA, Veize, J. Compilation and time-series analysis of a marine carbonate δ^18^O, δ^13^C, ^87^Sr/^86^Sr and δ^34^S database through Earth history. Earth– Sci Rev. 2008;87:113–33.

133. Hunt G, Cronin TM, Roy K. Species–energy relationship in the deep sea: a test using the Quaternary fossil record. Ecol Lett. 2005;8:739–47.

134. Marx FG, Uhen MD. Climate, critters, and cetaceans: Cenozoic drivers of the evolution of modern whales. Science. 2010;327:993–6.

135. Benson RBJ, Butler RJ. Uncovering the diversification history of marine tetrapods: ecology influences the effect of geological sampling biases. Geol Soc Spec Publ. 2011;358:191–208.

136. Wilberg EW, Turner AH, Brochu CA. Evolutionary structure and timing of major habitat shifts in Crocodylomorpha. Sci Rep. 2019;9:514.

137. Martins EP, Hansen TF. Phylogenies and the comparative method: a general approach to incorporating phylogenetic information into the analysis of interspecific data. Am Nat. 1997;149:646–67.

138. Pagel M. Inferring the historical patterns of biological evolution. Nature. 1999;401:877–84.

139. Orme CDL, Freckleton R, Thomas G, Petzoldt T, Fritz S, Isaac N, Pearse W. CAPER: comparative analyses of phylogenetics and evolution in R. R package version 1.0.1. 2018. https://CRAN.R-project.org/package=caper.

140. Garland T, Dickerman AW, Janis CM, Jones JA. Phylogenetic analysis of covariance by computer simulation. Syst Biol. 1993;42:265–92.

141. Pinheiro J, Bates D, DebRoy S, Sarkar D, R Core Team. nlme: linear and nonlinear mixed effects models. R package version 3.1–131. 2017. https://CRAN.R-project.org/package=nlme.

142. Revell LJ. phytools: an R package for phylogenetic comparative biology (and other things). Methods Ecol Evol. 2012;3:217–23.

143. Foote M. Discordance and concordance between morphological and taxonomic diversity. Paleobiology. 1993;19:185–204.

144. Foote M. The evolution of morphological diversity. Annu Rev Ecol Syst. 1997;28:129–52.

145. Wills MA. Morphological disparity: a primer. In: Adrain JM, Edgecombe GD, Lieberman BS, editors. Fossils, phylogeny, and form. Boston: Springer; 2001. p. 55–144.

146. Hopkins MJ, Gerber S. Morphological disparity. In: Nuño de la Rosa L, Müller GB, editors. Evolutionary Developmental Biology. Springer International Publishing; 2017. p. 1–12.

147. Burnham KP, Anderson DR, Huyvaert KP. AIC model selection and multimodel inference in behavioral ecology: some background, observations, and comparisons. Behav Ecol Sociobiol. 2011;65:23–35.

148. Marinho TS, Carvalho IS. An armadillo-like sphagesaurid crocodyliform from the Late Cretaceous of Brazil. J S Am Earth Sci. 2009;27:36–41.

149. Young MT, Tennant JP, Brusatte SL, Challands TJ, Fraser NC, Clark ND, Ross, DA. The first definitive Middle Jurassic atoposaurid (Crocodylomorpha, Neosuchia), and a discussion on the genus Theriosuchus. Zool J Linn Soc. 2016;176:443–62.

150. Brochu CA. A new alligatorid from the lower Eocene Green River Formation of Wyoming and the origin of caimans. J Vertebr Paleontol. 2010;30:1109–26.

151. Van Valen L. Adaptive zones and the orders of mammals. Evolution. 1971;25:420–8.

152. Landis MJ, Schraiber JG. Pulsed evolution shaped modern vertebrate body sizes. Proc Natl Acad Sci USA. 2017;114:13224–9.

153. Slater GJ, Pennell MW. Robust regression and posterior predictive simulation increase power to detect early bursts of trait evolution. Syst Biol. 2013;63:293–308.

154. Downhower JF, Blumer LS. Calculating just how small a whale can be. Nature. 1988;335:675.

155. Williams TM. The evolution of cost efficient swimming in marine mammals: limits to energetic optimization. Philos Trans R Soc Lond B-Biol Sci. 1999;354:193–201.

156. Vermeij GJ. Gigantism and its implications for the history of life. PLoS One. 2016;11:e0146092.

157. Ahlborn BK, Blake RW. Lower size limit of aquatic mammals. Am J Phys. 1999;67:920–2.

158. Smith EN. Heating and cooling rates of the American alligator, *Alligator mississippiensis*. Physiol Zool. 1976;49:37–48.

159. Zanno LE, Drymala S, Nesbitt SJ, Schneider VP. Early crocodylomorph increases top tier predator diversity during rise of dinosaurs. Sci Rep. 2015;5:9276.

160. Sookias RB, Benson RBJ, Butler RJ. Biology, not environment, drives major patterns in maximum tetrapod body size through time. Biol Lett. 2012;8:674–7.

161. Huttenlocker AK. Body size reductions in nonmammalian eutheriodont therapsids (Synapsida) during the end-Permian mass extinction. PLoS One. 2014;9:e87553.

162. Linnert C, Robinson SA, Lees JA, Bown PR, Pérez-Rodríguez I, Petrizzo MR, Falzoni F, Littler K, Arz JA, Russell EE. Evidence for global cooling in the Late Cretaceous. Nat Commun. 2014;5:4194.

163. Alroy J. A multispecies overkill simulation of the end-Pleistocene megafaunal mass extinction. Science. 2001;292:1893–6.

164. Johnson CN. Determinants of loss of mammal species during the Late Quaternary ‘megafauna’ extinctions: life history and ecology, but not body size. Proc R Soc B-Biol Sci. 2002;269:22213–7.

165. Fisher DO, Owens IP. The comparative method in conservation biology. Trends Ecol Evol. 2004;19:391–8.

166. Cardillo M, Mace GM, Jones KE, Bielby J, Bininda-Emonds OR, Sechrest W, Orme CDL, Purvis A. Multiple causes of high extinction risk in large mammal species. Science. 2005;309:1239–41.

167. Clauset A, Erwin DH. The evolution and distribution of species body size. Science. 2008;321:399–401.

168. Purvis A, Gittleman JL, Cowlishaw G, Mace GM. Predicting extinction risk in declining species. Proc R Soc B-Biol Sci. 2000;267:1947–52.

169. Van Valkenburgh B, Wang X, Damuth J. Cope’s rule, hypercarnivory, and extinction in North American canids. Science. 2004;306:101–4.

170. Tennant JP, Mannion PD, Upchurch P. Environmental drivers of crocodyliform extinction across the Jurassic/Cretaceous transition. Proc R Soc B-Biol Sci. 2016;283:20152840.

171. Tennant JP, Mannion PD, Upchurch P. Sea level regulated tetrapod diversity dynamics through the Jurassic/Cretaceous interval. Nat Commun. 2016;7:12737.

172. Tennant JP, Mannion PD, Upchurch P, Sutton MD, Price GD. Biotic and environmental dynamics through the Late Jurassic–Early Cretaceous transition: evidence for protracted faunal and ecological turnover. Biol Rev. 2017;92:776–814.

173. Fanti F, Miyashita T, Cantelli L, Mnasri F, Dridi J, Contessi M, Cau A. The largest thalattosuchian (Crocodylomorpha) supports teleosaurid survival across the Jurassic-Cretaceous boundary. Cretaceous Res. 2016;61:263–74.

174. Benson RBJ, Mannion PD, Butler RJ, Upchurch P, Goswami A, Evans SE. Cretaceous tetrapod fossil record sampling and faunal turnover: implications for biogeography and the rise of modern clades. Palaeogeogr Palaeocl. 2013;372:88–107.

175. Carvalho IS, Gasparini ZB, Salgado L, Vasconcellos FM, Marinho TS. Climate’s role in the distribution of the Cretaceous terrestrial Crocodyliformes throughout Gondwana. Palaeogeogr Palaeocl. 2010;297:252–62.

176. Pol D, Leardi JM. Diversity patterns of Notosuchia (Crocodyliformes, Mesoeucrocodylia) during the Cretaceous of Gondwana. In: Fernández M, Herrera Y, editors. Reptiles Extintos–Volumen en Homenaje a Zulma Gasparini. Buenos Aires: Asociación Paleontológica Argentina; 2015. p. 172–86

177. Brochu CA. Phylogenetic approaches toward crocodylian history. Annu Rev Earth Pl Sc. 2003;31:357–97.

178. Ősi A. The evolution of jaw mechanism and dental function in heterodont crocodyliforms. Hist Biol. 2014;26:279–414.

179. Pol D, Nascimento PM, Carvalho AB, Riccomini C, Pires-Domingues RA, Zaher H. A new notosuchian from the Late Cretaceous of Brazil and the phylogeny of advanced notosuchians. PLoS One. 2014;9:e93105.

180. Russell AP, Wu X. The Crocodylomorpha at and between geological boundaries. Zoology. 1997;100:164–82.

181. Jouve S, Bardet N, Jalil NE, Suberbiola XP, Bouya, Amaghzaz, M. The oldest African crocodylian: phylogeny, paleobiogeography, and differential survivorship of marine reptiles through the Cretaceous-Tertiary boundary. J Vertebr Paleontol. 2008;28:409–21.

182. Wilson GP. Mammals across the K/Pg boundary in northeastern Montana, USA: dental morphology and body-size patterns reveal extinction selectivity and immigrant-fueled ecospace filling. Paleobiology. 2013;39:429–69.

183. Longrich NR, Bhullar BAS, Gauthier JA. Mass extinction of lizards and snakes at the Cretaceous–Paleogene boundary. Proc Natl Acad Sci USA. 2012;109:21396–401.

## Supplementary references

Andrade MB, Edmonds R, Benton MJ, Schouten R. 2011A new Berriasian species of *Goniopholis* (Mesoeucrocodylia, Neosuchia) from England, and a review of the genus. Zoological Journal of the Linnean Society, 163S1, S66–S108.

Bapst DW. 2012. paleotree: an R package for paleontological and phylogenetic analyses of evolution. Methods in Ecology and Evolution, 3(5), 803–07.

Bapst DW. 2013. A stochastic rate calibrated method for time scaling phylogenies of fossil taxa. Methods in Ecology and Evolution, 4(8), 724–33.

Bapst DW. 2014. Preparing paleontological datasets for phylogenetic comparative methods. In: Garamszegi LZ (ed.) Modern phylogenetic comparative methods and their application in evolutionary biology. Berlin: Springer. p. 515–44.

Bapst DW. 2014. Assessing the effect of time-scaling methods on phylogeny-based analyses in the fossil record. Paleobiology, 40(3), 331–51.

Bates KT, Manning PL, Hodgetts D, Sellers WI. 2009. Estimating mass properties of dinosaurs using laser imaging and 3D computer modelling. PLoS One, 4, e4532.

Benson RBJ, Campione NE, Carrano MT, Mannion PD, Sullivan C, Upchurch P, Evans DC. 2014. Rates of dinosaur body mass evolution indicate 170 million years of sustained ecological innovation on the avian stem lineage. PLoS Biology, 12(5), e1001853.

Benson RBJ, Hunt G, Carrano MT, Campione N. 2018. Cope’s rule and the adaptive landscape of dinosaur body size evolution. Palaeontology, 61(1), 13–48.

Brochu CA. 2012. Phylogenetic relationships of Palaeogene ziphodont eusuchians and the status of *Pristichampsus* Gervais, 1853. Earth and Environmental Science Transactions of the Royal Society of Edinburgh, 103(3-4), 521–550.

Brochu CA, Parris DC, Grandstaff BS, Denton Jr RK, Gallagher WB. 2012. A new species of *Borealosuchus* (Crocodyliformes, Eusuchia) from the Late Cretaceous–early Paleogene of New Jersey. Journal of Vertebrate Paleontology, 32(1), 105–116.

Brocklehurst N. 2017. Rates of morphological evolution in Captorhinidae: an adaptive radiation of Permian herbivores. PeerJ, 5, e3200.

Bronzati M, Montefeltro FC, Langer MC. 2015. Diversification events and the effects of mass extinctions on Crocodyliformes evolutionary history. Royal Society Open Science, 2, 140385.

Buscalioni ÁD. 2017. The Gobiosuchidae in the early evolution of Crocodyliformes. Journal of Vertebrate Paleontology, 37(3), e1324459.

Bustard HR, Singh LAK. 1977. Studies on the Indian Gharial *Gavialis gangeticus* (Gmelin) (Reptilia, Crocodilia) – I: Estimation of body length from scute length. Indian Forester, 103(2), 140–149.

Campione NE, Evans DC. 2012. A universal scaling relationship between body mass and proximal limb bone dimensions in quadrupedal terrestrial tetrapods. BMC Biology, 10(1), 60.

Carballido JL, Pol D, Otero A, Cerda IA, Salgado L, Garrido AC, Ramezani J, Cúneo NR, Krause JM. 2017. A new giant titanosaur sheds light on body mass evolution among sauropod dinosaurs. Proceedings of the Royal Society B: Biological Sciences, 284(1860), 20171219.

Clark JM. 1994. Patterns of evolution in Mesozoic Crocodyliformes. In: Fraser NC, Sues HD (eds.) In the Shadow of Dinosaurs. Cambridge: Cambridge University Press. p. 84–97.

Clark JM. 2011. A new shartegosuchid crocodyliform from the Upper Jurassic Morrison Formation of western Colorado. Zoological Journal of the Linnean Society, 163S1, S152–S172.

Colbert EH. 1962. The weights of dinosaurs. American Museum Novitates, (2076), 1–16.

Currie PJ. 1978. The orthometric linear unit. Journal of Paleontology, 52, 964–971.

Farlow JO, Hurlburt GR, Elsey RM, Britton AR, Langston W. 2005. Femoral dimensions and body size of *Alligator mississippiensis*: estimating the size of extinct mesoeucrocodylians. Journal of Vertebrate Paleontology, 25(2), 354–369.

Godoy PL, Bronzati M, Eltink E, Marsola JCA, Cidade GM, Langer MC, Montefeltro FC. 2016. Postcranial anatomy of *Pissarrachampsa sera* (Crocodyliformes, Baurusuchidae) from the Late Cretaceous of Brazil: insights on lifestyle and phylogenetic significance. PeerJ, 4, e2075.

Hastings AK, Bloch JI, Jaramillo CA. 2015. A new blunt-snouted dyrosaurid, *Anthracosuchus balrogus* gen. et sp. nov. (Crocodylomorpha, Mesoeucrocodylia), from the Palaeocene of Colombia. Historical Biology, 27(8), 998–1020.

Hedman MM. 2010. Constraints on clade ages from fossil outgroups. Paleobiology, 36(1), 16– 31.

Herrera Y, Gasparini Z, Fernández MS. 2015. *Purranisaurus potens* Rusconi, an enigmatic metriorhynchid from the Late Jurassic–Early Cretaceous of the Neuquén Basin. Journal of Vertebrate Paleontology, 35(2), e904790.

Hall PM, Portier KM. 1994. Cranial morphometry of New Guinea crocodiles (*Crocodylus novaeguineae*): ontogenetic variation in relative growth of the skull and an assessment of its utility as a predictor of the sex and size of individuals. Herpetological Monographs, 203– 225.

Hurlburt G. 1999. Comparison of body mass estimation techniques, using recent reptiles and the pelycosaur *Edaphosaurus boanerges*. Journal of Vertebrate Paleontology, 19(2), 338–350.

Hurlburt GR, Heckert AB, Farlow JO. 2003. Body mass estimates of phytosaurs (Archosauria: Parasuchidae) from the Petrified Forest Formation (Chinle Group: Revueltian) based on skull and limb bone measurements. New Mexico Museum of Natural History and Science Bulletins, 24, 105–13.

Laurin M. 2004. The evolution of body size, Cope’s rule and the origin of amniotes. Systematic Biology, 53(4), 594–622.

Leardi JM, Pol D, Clark JM. 2017. Detailed anatomy of the braincase of *Macelognathus vagans* Marsh, 1884 (Archosauria, Crocodylomorpha) using high resolution tomography and new insights on basal crocodylomorph phylogeny. PeerJ, 5, e2801.

Lloyd GT, Bapst DW, Friedman M, Davis KE. 2016. Probabilistic divergence time estimation without branch lengths: dating the origins of dinosaurs, avian flight and crown birds. Biology letters, 12(11), 20160609.

Maddison WP, Maddison DR. 2018. Mesquite: a modular system for evolutionary analysis. Version 3.40. http://mesquiteproject.org

Martin JE, De’lfino M, Smith T. 2016. Osteology and affinities of Dollo’s goniopholidid (Mesoeucrocodylia) from the Early Cretaceous of Bernissart, Belgium. Journal of Vertebrate Paleontology, 36(6), e1222534.

Meunier LMV, Larsson HCE. 2017. Revision and phylogenetic affinities of *Elosuchus* (Crocodyliformes). Zoological Journal of the Linnean Society, 179, 169–200.

Montefeltro FC, Larsson HCE, França, MAG, Langer MC. 2013. A new neosuchian with Asian affinities from the Jurassic of northeastern Brazil. Naturwissenschaften, 100(9), 835–841.

Motani R. 2001. Estimating body mass from silhouettes: testing the assumption of elliptical body cross-sections. Paleobiology, 27, 735–50.

Narváez I, Brochu CA, Escaso F, Pérez-García A, Ortega F. 2015. New Crocodyliforms from Southwestern Europe and Definition of a Diverse Clade of European Late Cretaceous Basal Eusuchians. PLoS ONE, 10(11), e0140679.

Platt SG, Rainwater TR, Thorbjarnarson JB, Finger AG, Anderson TA, McMurry ST. 2009. Size estimation, morphometrics, sex ratio, sexual size dimorphism, and biomass of Morelet’s crocodile in northern Belize. Caribbean Journal of Science, 45(1), 80–94.

Platt SG, Rainwater TR, Thorbjarnarson JB, Martin D. 2011. Size estimation, morphometrics, sex ratio, sexual size dimorphism, and biomass of *Crocodylus acutus* in the coastal zone of Belize. Salamandra; 47, 179–92.

Pol D, Gasparini Z. 2009. Skull anatomy of *Dakosaurus andiniensis* (Thalattosuchia: Crocodylomorpha) and the phylogenetic position of Thalattosuchia. Journal of Systematic Palaeontology, 7(2), 163–197.

Pol D, Leardi JM, Lecuona A, Krause M. 2012. Postcranial anatomy of *Sebecus icaeorhinus* (Crocodyliformes, Sebecidae) from the Eocene of Patagonia. Journal of Vertebrate Paleontology, 32(2), 328–354.

Pol D, Rauhut OWM, Lecuona A, Leardi JM, Xu X, Clark JM. 2013. A new fossil from the Jurassic of Patagonia reveals the early basicranial evolution and the origins of Crocodyliformes. Biological Reviews. 88, 862–872.

Pol D, Nascimento PM, Carvalho AB, Riccomini C, Pires-Domingues RA, Zaher H. 2014. A New Notosuchian from the Late Cretaceous of Brazil and the Phylogeny of Advanced Notosuchians. PLoS ONE, 9(4), e93105.

R Core Team. 2018. R: a language and environment for statistical computing. Vienna: R Foundation for Statistical Computing. https://www.R-project.org/

Ristevski J, Young MT, Andrade MB, Hastings AK. 2018. A new species of *Anteophthalmosuchus* (Crocodylomorpha, Goniopholididae) from the Lower Cretaceous of the Isle of Wight, United Kingdom, and a review of the genus. Cretaceous Research, 84, 340–383.

Romer AS, Price LW. 1940. Review of the Pelycosauria. Geological Society of America Special Papers, 28, 1–534.

Scheyer TM, Aguilera OA, Delfino M, Fortier DC, Carlini AA, Sánchez R, Carrillo-Briceño JD, Quiroz L, Sánchez-Villagra MR. 2013. Crocodylian diversity peak and extinction in the late Cenozoic of the northern Neotropics. Nature communications, 4, 1907.

Schwarz D, Raddatz M, Wings O. 2017. *Knoetschkesuchus langenbergensis* gen. nov. sp. nov., a new atoposaurid crocodyliform from the Upper Jurassic Langenberg Quarry (Lower Saxony, northwestern Germany), and its relationships to Theriosuchus. PLoS ONE, 12(2), e0160617.

Sellers WI, Hepworth-Bell J, Falkingham PL, Bates KT, Brassey CA, Egerton VM, Manning PL. 2012. Minimum convex hull mass estimations of complete mounted skeletons. Biology Letters, 8(5), 842–845.

Sereno PC, Larsson HC, Sidor CA, Gado B. 2001. The giant crocodyliform *Sarcosuchus* from the Cretaceous of Africa. Science, 294(5546), 1516–1519.

Tennant JP, Mannion PD, Upchurch P. 2016. Evolutionary relationships and systematics of Atoposauridae (Crocodylomorpha: Neosuchia): implications for the rise of Eusuchia. Zoological Journal of the Linnean Society, 177(4), 854–936.

Turner AH. 2015. A review of *Shamosuchus* and *Paralligator* (Crocodyliformes, Neosuchia) from the Cretaceous of Asia. PLoS ONE, 10(2), e0118116.

Turner AH, Pritchard AC. 2015. The monophyly of Susisuchidae (Crocodyliformes) and its phylogenetic placement in Neosuchia. PeerJ, 3, e759.

Webb GJW, Messel H. 1978. Morphometric analysis of *Crocodylus porosus* from the north coast of Arnhem Land, northern Australia. Australian Journal of Zoology, 26(1), 1–27.

Wilberg E. 2015. What’s in an Outgroup? The Impact of Outgroup Choice on the Phylogenetic Position of Thalattosuchia (Crocodylomorpha) and the Origin of Crocodyliformes. Systematic Biology, 64(4), 621–37.

Young MT. 2014. Filling the “Corallian Gap”: re-description of a metriorhynchid crocodylomorph from the Oxfordian (Late Jurassic) of Headington, England. Historical Biology, 26, 80–90.

Young MT, Bell MA, Andrade MB, Brusatte SL. 2011. Body size estimation and evolution in metriorhynchid crocodylomorphs: implications for species diversification and niche partitioning. Zoological Journal of the Linnean Society, 163(4), 1199–1216.

Young MT, Rabi M, Bell MA, Foffa, D, Steel L, Sachs S, Peyer K. 2016. Big-headed marine crocodyliforms and why we must be cautious when using extant species as body length proxies for long-extinct relatives. Palaeontologia Electronica, 19(3), 1–14.

Young MT, Hastings AK, Allain R, Smith TJ. 2017. Revision of the enigmatic crocodyliform *Elosuchus felixi* de Lapparent de Broin, 2002 from the Lower-Upper Cretaceous boundary of Niger: potential evidence for an early origin of the clade Dyrosauridae. Zoological Journal of the Linnean Society, 179, 377–403.

